# Cannabidiol targets a modulatory system for excitatory-inhibitory synaptic coordination, contributing to its anti-seizure action

**DOI:** 10.1101/2022.09.27.509638

**Authors:** Evan Rosenberg, Simon Chamberland, Michael Bazelot, Erica R. Nebet, Xiaohan Wang, Sam McKenzie, Swati Jain, Stuart Greenhill, Max Wilson, Alejandro Salah, Shanice Bailey, Pabitra Hriday Patra, Rebecca Rose, Nicolas Chenouard, Simon D. Sun, Drew Jones, György Buzsáki, Orrin Devinsky, Gavin Woodhall, Helen Scharfman, Benjamin Whalley, Richard Tsien

## Abstract

Cannabidiol (CBD), a non-euphoric component of cannabis, reduces seizures in multiple forms of pediatric epilepsy, but the mechanism(s) of anti-seizure action remain unclear. In one leading model, CBD acts at glutamatergic axon terminals, blocking pro-excitatory actions of an endogenous membrane phospholipid, lysophosphatidylinositol (LPI), at the G protein-coupled receptor GPR55. However, the impact of LPI-GPR55 signaling at inhibitory synapses and in epileptogenesis remains underexplored. We found that LPI transiently increased hippocampal CA3→CA1 excitatory presynaptic release probability and evoked synaptic strength in WT mice, while attenuating inhibitory postsynaptic strength by decreasing GABA_A_Rγ_2_ and gephyrin puncta. Effects of LPI at both excitatory and inhibitory synapses were eliminated by CBD pretreatment and absent after GPR55 deletion. Acute pentylenetrazole-induced seizures elevated levels of GPR55 and LPI, and chronic lithium pilocarpine-induced epileptogenesis potentiated the pro-excitatory effects of LPI. We propose that CBD exerts potential therapeutic effect both by blocking synaptic effects of LPI and dampening hyperexcitability.

## Introduction

Neuronal circuits require coordination between synaptic excitation (E) and inhibition (I) for proper function (Liu, 2004; Xue et al., 2014), as disruptions in the E:I ratio contribute to disorders such as epilepsy (Paz and Huguenard, 2015), autism spectrum disorders (Rubenstein and Merzenich, 2003), and schizophrenia (Lewis et al., 2012). G protein-coupled receptors (GPCRs) tightly regulate the interplay between E and I by linking agonists such as endocannabinoids to downstream signaling (Katona and Freund, 2008; Yamashita et al., 2013). However, in many cases, the exact nature of GPCR signaling remains obscure. Understanding the effects and interactions of the agonists and antagonists of these receptors, their signaling pathways, and downstream effector mechanisms would greatly enhance our understanding of CNS networks.

(-)-Trans-cannabidiol (CBD), a major non-psychoactive component derived from cannabis, can reduce seizure activity in multiple animal models (Rosenberg et al., 2017) and in patients with treatment-resistant forms of epilepsy. Promising results from double-blind, placebo-controlled phase III clinical trials in Dravet Syndrome (Devinsky et al., 2017) Lennox-Gastaut Syndrome (Devinsky et al., 2018a), and Tuberous Sclerosis (Thiele et al., 2021) have contributed to disease-specific FDA approval of highly purified, plant-derived cannabidiol (Epidiolex® in the U.S.) for multiple clinical populations. Preclinical experiments verify that CBD reduces spontaneous recurrent seizures in a chronic epilepsy model (Patra et al., 2019) and suggest that CBD modulates E:I coordination (Kaplan et al., 2017; Khan et al., 2018), but the molecular signaling underlying CBD’s anti-seizure actions is not clearly defined (Gray and Whalley, 2020).

Multiple molecular targets of CBD have been proposed as mediators of its therapeutic action, including ion channels, transporters, and transmembrane signaling proteins (Ibeas Bih et al., 2015). Among the proposed candidates are two G protein-coupled receptors: the cannabinoid receptor CB_1_R (Laprairie et al., 2015; Straiker et al., 2018) and the de-orphanized receptor GPR55 (Oka et al., 2007; Sylantyev et al., 2013). While CBD has minimal activity at the CB_1_R orthosteric binding site (Bisogno et al., 2001; Jones et al., 2010; Pertwee, 2007; Thomas et al., 1998), it has been proposed to act as a CB_1_R negative allosteric modulator (Laprairie et al., 2015; Straiker et al., 2018). The relevance of such allosteric modulation to *in vivo* epilepsy remains uncertain, as CB_1_R antagonists are mostly proconvulsive (Rosenberg et al., 2017).

CBD also operates as a GPR55 antagonist, blocking the effects of the lysophosphatidylinositol (LPI), an endogenous lipid agonist of GPR55 (Oka et al., 2007; Ryberg et al., 2007). LPI drives GPR55-mediated Ca^2+^ flux at neuronal cell bodies (Lauckner et al., 2008) and presynaptic CA3-CA1 excitatory terminals in the hippocampus, producing a pro-excitatory effect. However, CBD blocks LPI-mediated presynaptic Ca^2+^ rises, preventing an elevation in glutamate release (Sylantyev et al., 2013). LPI effects were completely absent in GPR55 KO mice, suggesting agonist-receptor selectivity (Sylantyev et al., 2013). Further, a synthetic GPR55 inverse agonist occluded CBD action to reduce hippocampal excitability (Kaplan et al., 2017); see also (Khan et al., 2018). Collectively, these observations implicate GPR55 as a substrate for mediating some of the anti-seizure actions of CBD.

However, these studies leave many questions unanswered. If CBD acts on multiple types of neurons, how do the responses converge to affect excitability? Are the critical CBD effect(s) alterations of intrinsic excitability and/or synaptic strength, and if the latter, at pre- or postsynaptic loci? How do the cellular mechanism(s) impact spike transmission and information flow in circuits, and how does the underlying signaling change with repeated seizures and chronic epilepsy?

To address these questions, we focused on the LPI-GPR55 lipid signaling system as a candidate mediator of the anti-seizure action of CBD, and investigated how this system might regulate excitatory and inhibitory synaptic function. We observed that LPI triggers a GPR55-dependent, dual mechanism to elevate network excitability: a transient elevation in presynaptic excitatory release probability (Sylantyev et al., 2013), followed by a sustained down-regulation of inhibitory synaptic strength. The attenuation of synaptic inhibition, a novel role for LPI, was mediated by a slowly developing downregulation of clusters of γ_2_-containing postsynaptic GABA_A_R and gephyrin scaffolding. At synaptic and circuit levels, both LPI-mediated effects on synaptic transmission were blocked by GPR55 deletion or pre-treatment with CBD. Further, we found that acute seizures upregulate GPR55 expression and LPI production, thus markedly potentiating their combined pro-excitatory effects. Taken together, our observations reveal a positive feedback loop whereby LPI elevates excitability, which in turn increases expression of its target receptor, GPR55. We propose that elevated LPI-GPR55 signaling contributes a potential target for CBD action in reducing seizures, complementary to ion channels supporting neuronal excitability (Ghovanloo et al., 2018; Kaplan et al., 2017; Khan et al., 2018; Patel et al., 2016; Zhang and Bean, 2021).

## Results

### Hippocampal GPR55 expression and LPI-mediated enhancement of spike throughput

We studied GPR55-mediated mechanisms of CBD action on the hippocampus, whose circuitry contributes to temporal lobe seizures (Alexander et al., 2016). GPR55 was strongly expressed in the pyramidal layer of areas CA1 and CA3 of hippocampus derived from male wild-type C57Bl/6J mice, with lower expression in area CA2 and dentate gyrus granule layer (Fig. 1A, B: n=8 slices, CA1 vs CA2 p=0.033, CA1 vs DG p=0.0002, CA3 vs DG p=0.0006). In confocal images at higher resolution, punctate GPR55 labeling also occurs in the CA1 *stratum radiatum*, consistent with localization at incoming CA3 Schaffer Collateral axon terminals (Sylantyev et al., 2013) (Suppl. Fig. 3A), although glial and postsynaptic expression cannot be excluded. Anti-GPR55 antibody labeling was greatly reduced in images from GPR55 KO mice (male *B6;129S-Gpr55^tm1Lex^/Mmnc*) at both low (10x) and high (63x) magnification (Fig. 1A, B: n=8 WT vs n=3 KO slices, p<0.0001 for all regions), demonstrating the specificity of the anti-GPR55 antibody for immunocytochemistry. These data suggested that GPR55 is well-expressed at CA3 to CA1 inputs, poised to regulate synaptic strength at this connection.

**Fig. 1:**
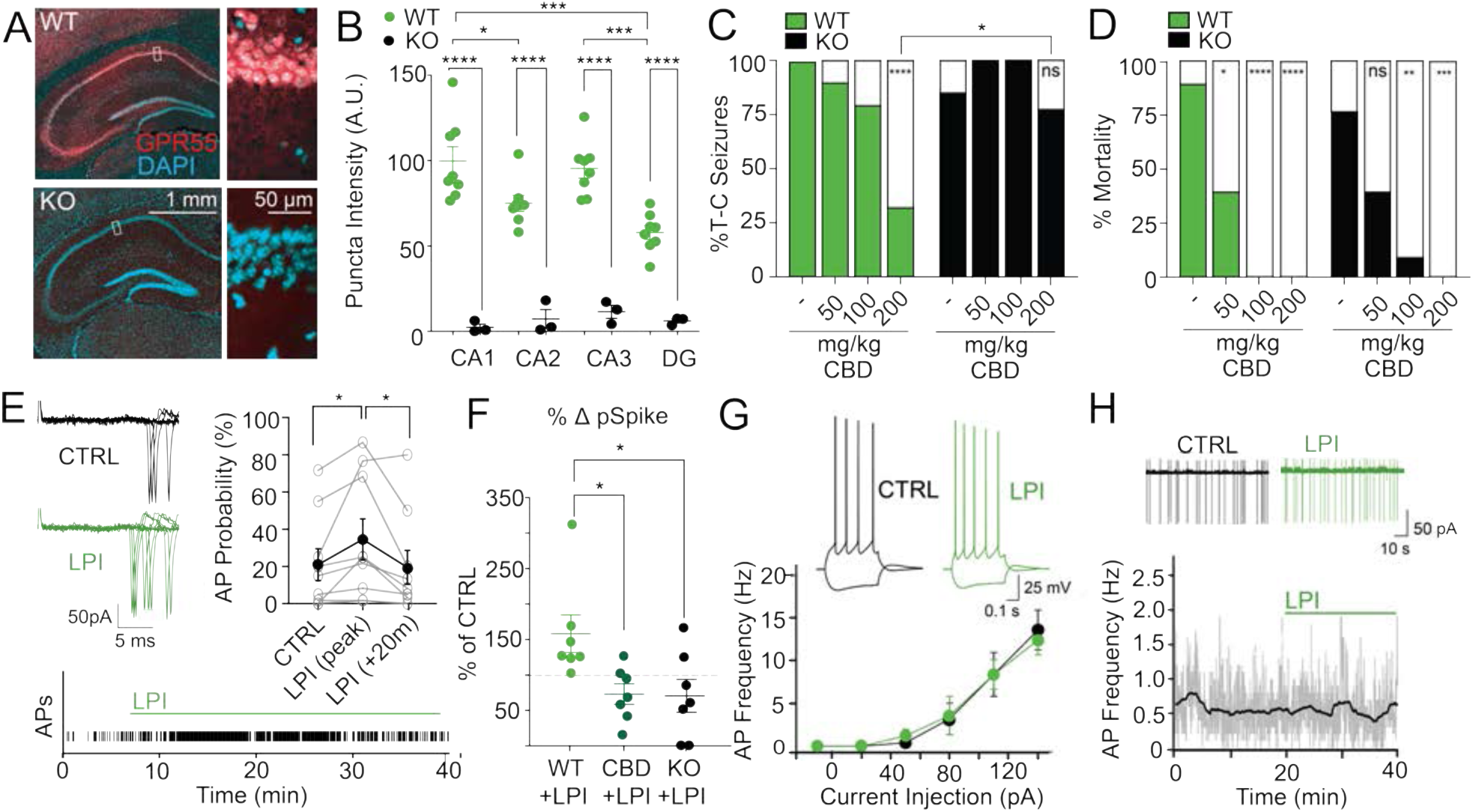
CBD reduces seizures and LPI-driven CA3 →CA1 spike enhancement via GPR55. (A) GPR55 expression in hippocampal slices at 10x and 63x resolution is absent in slices from the GPR55 KO mouse. (B) GPR55 expression (20x) was greater in the pyramidal layer of area CA1 and CA3 than the pyramidal layer of CA2 or granule layer of DG (CA1 vs CA2 p=0.030, CA1 vs DG p=0.0002, CA3 vs DG p=0.0006). GPR55 expression was significantly reduced in all regions in slices from GPR55 KO littermate controls (p<0.0001). (C) CBD exerted a dose-dependent reduction in seizures generated by the chemoconvulsant pentylenetetrazole (PTZ, 105 mg/kg), with significant effects at 200 mg/kg, ip. (p<0.0001). However, CBD did not reduce PTZ-induced seizures in GPR55 KO mice (200 mg/kg CBD: p>0.999 vs KO veh controls, p=0.03 vs. WT CBD 200 mg/kg). (D) CBD decreased mortality in seizures induced by PTZ in both WT (50 mg/kg p=0.011, 100mg/kg p=0.0008, and 200 mg/kg p<0.0001 vs vehicle controls) and GPR55 KO mice (100 mg/kg p=0.0028, 200 mg/kg p=0.0001 vs vehicle controls). (E-F) In cell-attached recordings, LPI acutely elevated the probability of evoking an action potential induced by Schaffer Collateral synaptic stimulation (5x 50Hz at 5s increments) (p=0.038), an effect reversed 20 min following onset of LPI treatment (p=0.034 vs peak). The LPI-induced effect was blocked by pre-treatment with 1 μM CBD (p=0.025) and absent in GPR55 KO slices (p=0.021). (G) The current-frequency (F-I) properties and (H) cell-attached firing rate of CA1 pyramidal neurons remain unchanged upon addition of LPI (4 μM). For this and all figures hereafter, graph values and error bars represent mean of independent samples ± standard error of mean; *p≤0.05, **p≤0.01, ***p≤0.001, ****p<0.0001.

### GPR55 is implicated in the seizure-reducing properties of CBD

To investigate GPR55 as a potential target for the anticonvulsant effects of CBD, we first studied an *in vivo* model of acute, generalized seizures in mice, using the chemoconvulsant pentylenetetrazole (PTZ, 105 mg/kg, i.p.) (Fig. 1C). Administration of CBD (50, 100, 200 mg/kg, i.p., 1 h prior to PTZ injection) (Deiana et al., 2012) produced a dose-dependent reduction in tonic-clonic seizures, with a significant effect at 200 mg/kg, i.p. (n=18/18 WT Veh vs n=5/15 WT CBD 200 mg/kg, p<0.0001). In contrast, the seizure-reducing effects of CBD were absent in GPR55 KO mice, even at the highest dose of 200 mg/kg (n=11/13 KO Veh vs n=10/13 KO CBD 200 mg/kg, p>0.9999). At 200 mg/kg, the proportion of mice with tonic-clonic seizures was significantly higher in GPR55 KO mice than in WT (p=0.03). Thus, dampening of behavioral seizures appeared to be GPR55-dependent.

CBD also reduced mortality in both WT (Fig. 1D: n=16/18 death in WT Veh vs 0/15 WT CBD 200 mg/kg, p<0.0001) and KO mice (n=10/13 KO Veh vs n=0/13 KO CBD mg/kg, p=0.0001), suggesting a potential GPR55-independent effect of CBD. To confirm that our behavioral assay was not missing covert seizure activity that could contribute to mortality, we performed tungsten electrode recordings from CA1 (n=6 mice) and verified that electrographically monitored seizure activity was well correlated to behavioral assays in either WT (n=3) or GPR55 KO (n=3) genotypes (Suppl. Fig. 1A-C). Overall, these results suggested that GPR55 might play a key role in the anticonvulsive effect of CBD, but potentially not seizure-induced mortality, prompting further investigation into the interplay between CBD and GPR55 function in seizure-prone neuronal circuits.

The striking effects of deleting the GPR55 receptor suggest that its endogenous agonist, LPI, promotes seizure-like activity (Sylantyev et al., 2013). To explore this, we first tested the effect of LPI on synaptically driven excitation by stimulating the CA3→CA1 (Schaffer Collateral) input. Exposure to LPI transiently increased the probability of observing a spike (Fig. 1E: 21±9% baseline vs 35±11% peak, n=9, p=0.038). This enhancement of excitation-spike coupling (Fig. 1F: 155±26% relative to baseline) was completely prevented by pre-treatment with 1 μM CBD (Fig. 1F: CBD+LPI 71±14%, n=7, p=0.025) and absent in slices from GPR55 KO mice (Fig. 1F: 68±22%, n=7, p=0.021, see also Suppl. Fig. 1D).

We next looked for changes in principal cell intrinsic properties that would contribute to excess circuit excitability. Exposing hippocampal slices to LPI (4 μM) did not change the intrinsic current-frequency (F-I) relation (Fig. 1G, n=6) or spontaneous firing rate of CA1 pyramidal cells (Fig. 1H, Suppl. Fig. 1E: baseline 0.68±0.13 Hz vs LPI 0.72±0.15 Hz, n=11, p=0.74). Lacking a detectable change in intrinsic firing, the elevated responsiveness of pyramidal cells to Schaffer Collateral input (Fig. 1E, F) suggested that LPI strengthens the net synaptic drive and that CBD, acting as a GPR55 antagonist, opposes this effect to dampen feedforward CA3→CA1 signaling. This reasoning prompted further study of CBD, LPI, and GPR55 at the synaptic level.

### CBD blocks the pro-excitatory and disinhibitory effects of LPI

Coupling between axonal inputs and postsynaptic firing depends on both monosynaptic excitation (E→E) and disynaptic feedforward inhibition (E→ I→ E), prompting separate examination of excitatory and inhibitory synaptic transmission. In excitatory boutons, LPI releases Ca^2+^ from presynaptic stores and transiently enhances glutamatergic transmission in a GPR55-dependent and CBD-sensitive manner (Sylantyev et al., 2013). We also found that LPI (4 μM) enhanced spontaneous neurotransmission, studied by whole cell recordings of miniature excitatory postsynaptic currents (mEPSCs, in 1 μM TTX) in CA1 pyramids (Fig. 2A_1_-C_1_, light green traces; Suppl. Fig. 2A, green: 155±9% 5-10 min post LPI application vs baseline, n=10 neurons, p=0.0001), with no associated changes in mEPSC amplitude (97±3% vs baseline, n=10, p=0.30).

**Fig. 2:**
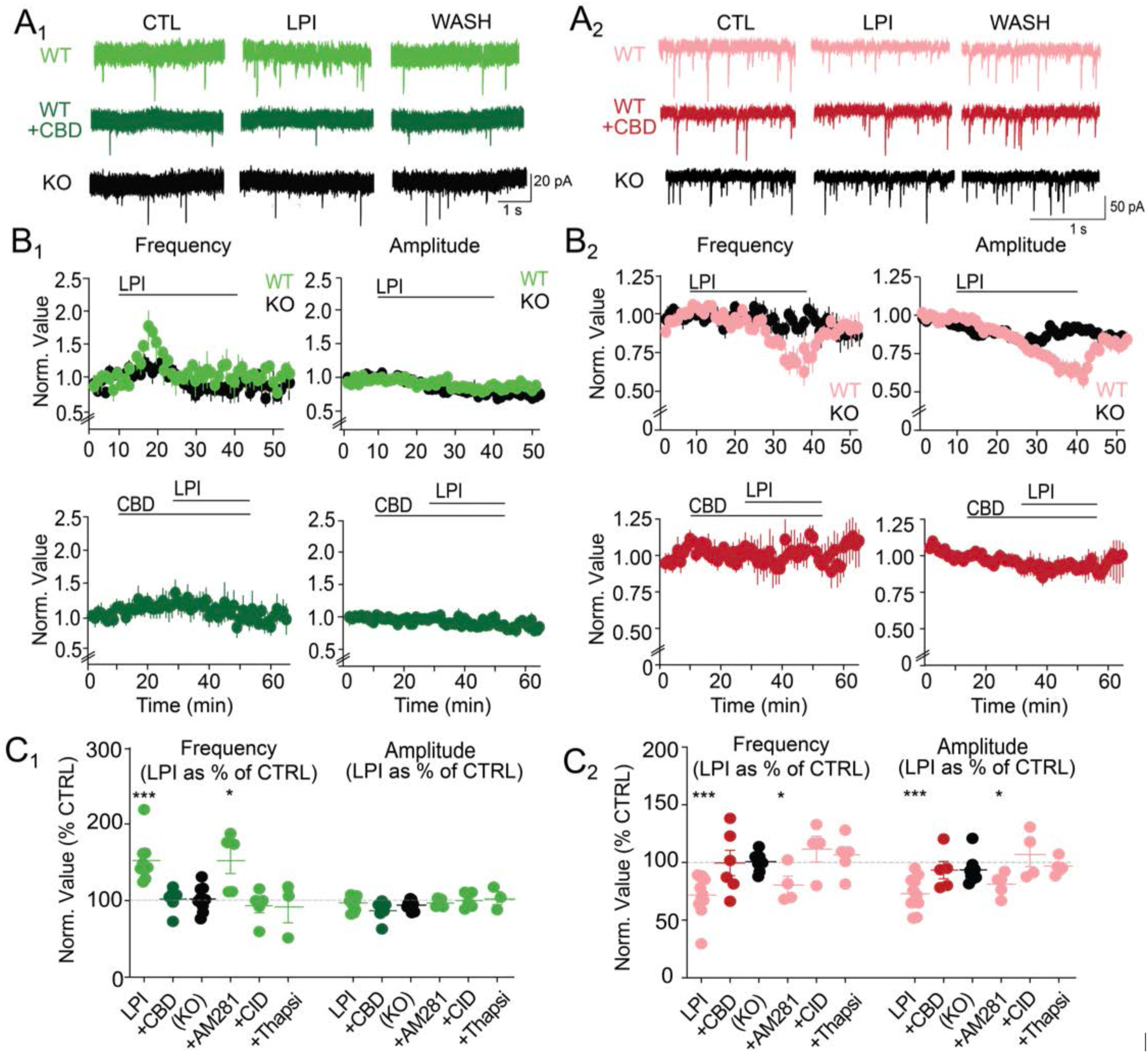
CBD blocks the pro-excitatory and anti-inhibitory effects of LPI on spontaneous synaptic transmission. (A_1-2_) Representative electrophysiological traces indicating that LPI (4 μM) reversibly elevated mEPSC frequency, without changing mEPSC amplitude (A_1_, light green traces). Contrarily, LPI diminished both the frequency and the amplitude of mIPSCs, annulled during the wash period (A_2_, pink traces). Pre-treatment of slices with CBD (1 μM) prevented LPI-mediated increases in mEPSC frequency (A_1_, dark green) and LPI-driven decreases in mIPSC amplitude and frequency (A_2_, red). Additionally, effects of LPI were absent in slices from GPR55 KO mice (A_1-2_, black traces). (B_1-2_) Time courses of LPI-mediated effects demonstrated a transient rise in mEPSCs (∼ 5 min to peak, light green, upper panel, left), and a slow, gradual reduction in mIPSC frequency and amplitude (20-30 min, pink, upper panels, right). These effects were not apparent in slices from GPR55 KO mice (black symbols, rear). CBD pre-treatment (B, lower panels) completely prevented the LPI-driven rise in mEPSC frequency (B_1_, dark green) and reduction in mIPSC frequency and amplitude (B_2_, red). Circles and bars represent mean±standard error of mean. (C_1-2_) Quantification of the peak effect of LPI, sampled during a 5 min time window, 5 min post LPI (mEPSCs, C_1_) or 25 min post LPI (mIPSCs, C_2_). Effect of LPI to both elevate mEPSC frequency (left, p=0.0001 vs baseline) and lower mIPSC frequency (right, p=0.0003) was absent following CBD application (dark green, C_1_; red, C_2_) and ablated in GPR55 KO mice (black). However, effects persisted in the presence of the CB_1_R antagonist AM281 (1 μM) (p=0.048 for mEPSCs, p=0.034 for mIPSCs). LPI-mediated changes were also blocked by the synthetic GPR55 inverse agonist CID16020046 (CID, 2.5 μM) and the SERCA pump inhibitor thapsigargin (Thapsi, 10 μM).

We next explored a potential role of LPI in regulating inhibitory signaling, as CBD rescues deficits caused by Nav1.1 sodium channel mutations in GABAergic neurons (Kaplan et al., 2017), that might contribute to its anti-seizure effect in patients with Dravet Syndrome (Devinsky et al., 2017). Miniature inhibitory postsynaptic currents (mIPSCs) recorded at V_rest_ in the presence of TTX (see Fig. 2 legend for details) were unchanged immediately following LPI application, but after 20-30 min of continuous exposure, exhibited a substantial decrease in both mIPSC frequency and amplitude (Fig. 2A_2_-C_2_, pink traces). Both the decrease in mIPSC rate (Fig. 2A_2_-C_2_, Suppl. Fig. 2C, D: 72±5%, 25-30 min post LPI vs baseline, n= 10, p=0.0002) and the attenuation of mIPSC size (72±5% vs baseline, n=10, p=0.0003) were reversed by removing LPI from the bathing medium (Fig. 2B_2_, pink traces, Suppl. Fig. 2D).

Uncovering a delayed disinhibitory action of LPI led us to examine the interplay between LPI and CBD. CBD exposure alone without LPI did not change mEPSC frequency relative to baseline (Fig. 2B_1_ dark green:108±14%, n=6, p=0.51). However, 1 μM CBD pre-treatment of slices completely prevented the pro-excitatory effects of LPI (Fig. 2A_1_-C_1_: 102±6% CBD+LPI vs baseline, n=6, p=0.82). Likewise, CBD without LPI did not affect mIPSC frequency (Fig. 2B_2_ red: 101±7% vs baseline, n=5, p=0.65) or amplitude (99±4%, n=5, p=0.94). As with mEPSCs, 1 μM CBD pre-treatment of slices fully blocked the LPI-mediated decrease in mIPSC frequency and amplitude (Fig. 2A_2_-C_2_: Freq: 103±13% vs baseline, n=5, p=0.97; Ampl: 90±8% vs baseline, n=5, p=0.32).

GPR55 receptor deletion eliminated both the pro-excitatory and anti-inhibitory effects induced by LPI (Fig. 2, black traces, Suppl. Fig. 2). Slices from GPR55 KO mice showed no LPI-mediated increase in mEPSC frequency (102±6% vs baseline, n=9, p=0.54) nor LPI-driven reductions in either mIPSC frequency (101±3% vs baseline, n=8, p=0.47) or amplitude (92±5% vs baseline, n=8, p=0.19). Thus, elimination of GPR55 had the same effect as CBD in almost all aspects of unitary synaptic transmission. A possible exception was basal mEPSC amplitude (Suppl. Fig. 2A) which appeared slightly larger in neurons from GPR55 KO mice (black, 26±1 pA) than in WT littermate controls (green, 22±1 pA, p=0.018), suggesting that GPR55 receptor deletion could have a developmental effect on excitatory synapses (Cherif et al., 2015). In contrast, basal mIPSC amplitude (Suppl. Fig. 2C) was not different in GPR55 KO (27±3 pA) and WT recordings (30±2 pA, p=0.37).

The dramatic effect of GPR55 removal leaves open the possibility that CB_1_Rs could modulate LPI signaling through partial colocalization and possible heterodimerization with GPR55 (Kargl et al., 2012). We tested this using AM281, an antagonist of CB_1_Rs, with minimal activity at GPR55 at 1 μM (Ryberg et al., 2007) (Fig. 2C_1_). However, LPI-mediated enhancement of mEPSC frequency persisted with 1 μM AM281 (140±17% vs baseline, n=5, p=0.048), similar to previous data (Sylantyev et al., 2013). Similarly, the LPI-mediated reduction in mIPSC amplitude and frequency remained robust with AM281 (Fig. 2C_2_, Freq: 80±8% vs baseline, n=4, p=0.034; Ampl: 81±4%, n=4, p=0.016), suggesting that LPI is likely not acting through CB_1_Rs.

Because CBD has several proposed non-GPR55 targets (Ibeas Bih et al., 2015), we tested further for GPR55 involvement using the synthetic GPR55-selective inverse agonist CID16020046 (Fig. 2C_1_). As expected, pre-treatment of slices with CID16020046 (2.5 μM) abolished the LPI-induced elevation of mEPSC frequency (84±13% vs baseline, n=5, p=0.10). Similarly, CID16020046 pretreatment prevented the anti-inhibitory effect of LPI on mIPSC frequency (Fig. 2C_2:_112±11% vs baseline, n=4, p=0.44) and amplitude (108±10%, n=4, p=0.52).

Although the pro-excitatory and anti-inhibitory effects of LPI displayed different time courses, the possibility remained that both effects on synaptic signaling might be facilitated by a GPR55-mediated discharge of intracellular calcium stores. To test this, we exposed slices to thapsigargin (10 μM) to pre-deplete Ca^2+^ stores, and found a complete block of the LPI-induced responses (Fig. 2C_1-2_), including mEPSC frequency (92±21% vs baseline, n=3, p=0.76), mIPSC frequency (106±8% vs baseline, n=5, p=0.58) and mIPSC amplitude (96±3%, n=5, p=0.44). These data indicated that LPI acts through CBD-sensitive GPR55 receptors to release intracellular Ca^2+^ and thereby modify both excitatory and inhibitory synapses.

We used Ca^2+^ imaging to localize critical Ca^2+^ stores, as LPI-mediated increases in presynaptic Ca^2+^ and mEPSC frequency were prevented by thapsigargin treatment in hippocampal slices (Fig. 2C_1_; Sylantyev et al. 2013). Finding that LPI does not increase mIPSC frequency (Fig. 2B_2_) suggested that Ca^2+^ stores in inhibitory presynaptic boutons might react differently. We performed two-photon Ca^2+^ imaging in GABAergic terminals from GCaMP6f-expressing, PV+ interneurons (male PV-Cre x Ai148 mice) to assay presynaptic Ca^2+^ regulation by LPI-GPR55 signaling. We focused on inhibitory presynaptic terminals in the perisomatic region of CA1 pyramidal neurons which predominantly emanate from PV+ interneurons. Applying LPI (4 μM) for 3 min did not affect presynaptic Ca^2+^ in the GABAergic terminals whereas subsequent exposure to [K]-rich solution reliably elevated Ca^2+^ (Suppl. Fig. 2E). Thus, inhibitory presynaptic boutons differ from their excitatory counterparts: LPI failed to discharge presynaptic Ca^2+^ stores of inhibitory terminals, suggesting that it more likely acts postsynaptically (Lauckner et al., 2008; Sylantyev et al., 2013).

### GPR55 expression at excitatory and inhibitory synapses

Given the effects of LPI on excitatory and inhibitory synaptic transmission in CA1, we next performed light microscopic immunocytochemistry on hippocampal slices (Suppl. Fig. 3A) to assess the co-localization of GPR55 with presynaptic and postsynaptic markers for excitatory synapses (the vesicular glutamate transporter 1 (Vglut1) and post-synaptic density protein (Psd-95)) and for inhibitory synapses (vesicular GABA transporter (Vgat) and post-synaptic Gephyrin). In both *stratum radiatum* (S.R.) and *stratum pyramidale* (S.P.), GPR55 puncta colocalized more strongly with Vglut1 than with the inhibitory presynaptic marker Vgat (Suppl. Fig. 3A S.R. p=0.044, S.P. p=0.020) and much more strongly with Gephyrin than Vgat (S.R. p=0.014, S.P. p=0.027). These observations suggest that GPR55 is likely situated at postsynaptic sites but shows less colocalization with inhibitory presynaptic terminals, providing a rationale for the absence of LPI-induced Ca^2+^ transient (Suppl. Fig. 2E) and the lack of LPI-induced elevation of mIPSC frequency (Fig. 2B_2_). However, our colocalization analyses are limited to the spatial resolution imposed by three-dimensional light microscopy (∼200 nm XY), and electron microscopy analyses will be required to determine exact pre- and postsynaptic GPR55 locations.

For better spatial resolution and less background fluorescence, we next examined GPR55 localization at hippocampal synapses in dissociated cell cultures using high-resolution confocal microscopy. Immunostaining of hippocampal cultures from GPR55 KO animals demonstrated specificity of GPR55 immunostaining (Suppl. Fig. 3B: GPR55 intensity WT vs KO p<0.0001). As with acute slice, GPR55 colocalized with a greater proportion of synapses labeled with Vglut1 than Vgat (Suppl. Fig. 3C, D, G, p=0.0219), and with more postsynapses labeled with Gephyrin than presynaptic terminals with Vgat (Suppl Fig. 3D, G: p<0.0001). Additionally, GPR55 overlapped with Gephyrin to a greater degree than Psd-95 (p=0.0092). To determine if GPR55 puncta colocalized with synaptic markers to a greater degree than chance alone, we scrambled GPR55 pixels along a 2D axon and compared colocalization levels to non-randomized conditions (Suppl. Fig. 3G, see Materials and Methods for details). Using this analysis, GPR55 colocalized better than chance with Vglut1 (p=0.0002), Psd-95 (p<0.0001), and Gephyrin (p<0.0001), but not Vgat (p=0.93). Thus, GPR55 colocalizes with excitatory synaptic markers and postsynaptic inhibitory markers, in line with the effects of GPR55 ligands on synaptic function.

We next looked for colocalization between GPR55 and another presynaptically localized GPCR sensitive to cannabinoids, CB_1_R (Dudok et al., 2015), of known importance for synaptic regulation (Castillo et al., 2012). CB_1_R can heterodimerize with GPR55 to reciprocally regulate cannabinoid signaling (Kargl et al., 2012). GPR55 appeared to alternate expression along a single axon (Suppl. Fig. 3E, linescan), and colocalized with only a small minority of total CB_1_R puncta in both acute slice and cell culture (Suppl. Fig. 3A, E, G). However, colocalization of GPR55 with CB_1_R was greater than chance when compared with randomized GPR55 pixels along an axon (Suppl. Fig. 3G, p=0.0061). Thus, GPR55 and CB_1_R likely occupy complementary structural regions in a single axon, but might colocalize or heterodimerize in some receptors.

### LPI modulates evoked transmission by enhancing excitation and reducing inhibition

After establishing that LPI modulates unitary synaptic currents via synaptic GPR55, we next studied action potential-evoked transmission in the canonical CA3 → CA1 microcircuit, recording from CA1 pyramidal neurons while stimulating Schaffer Collateral axons in S.R. (Fig 3A_1_, 10×10Hz). Whole-cell voltage clamp recordings of isolated inward excitatory currents were performed with cells held at E_Cl_ (∼ −70 mV) to eliminate contribution from GABAergic IPSCs. Exposure to LPI (4 μM) caused a reversible 22±6% elevation in the amplitude of evoked EPSCs during a 5 min window beginning 5 min post LPI application (Fig. 3A_1-3_: baseline 163±14 pA vs LPI 191±11 pA, n=5, p=0.0214). Concurrently, the paired pulse ratio (PPR), obtained as the ratio of amplitudes of the second and first evoked EPSCs, fell by 17±3% (baseline 1.6±0.2 vs LPI 1.3±0.1, n=5, p=0.0092). This combination of effects suggested that LPI transiently elevates the probability of evoked excitatory transmitter release, consistent with observations that LPI elevates mEPSC frequency (Sylantyev et al., 2013) (Fig. 2B_1_), field EPSP strength (Sylantyev et al., 2013), and long-term potentiation (Hurst et al., 2017). Pre-treatment with 1 μM CBD prevented the increased eEPSC amplitude (p=0.0057) and the reduced PPR (p=0.042, n=5 for all conditions) (Fig. 3A_3_, Suppl. Fig. 4). Further, GPR55 deletion produced similar effects (Fig. 3A_3_, Suppl. Fig. 4): eliminating the LPI-driven changes in eEPSC amplitude (p=0.0277), and in PPR (p=0.048, n=5 for all conditions).

**Fig. 3:**
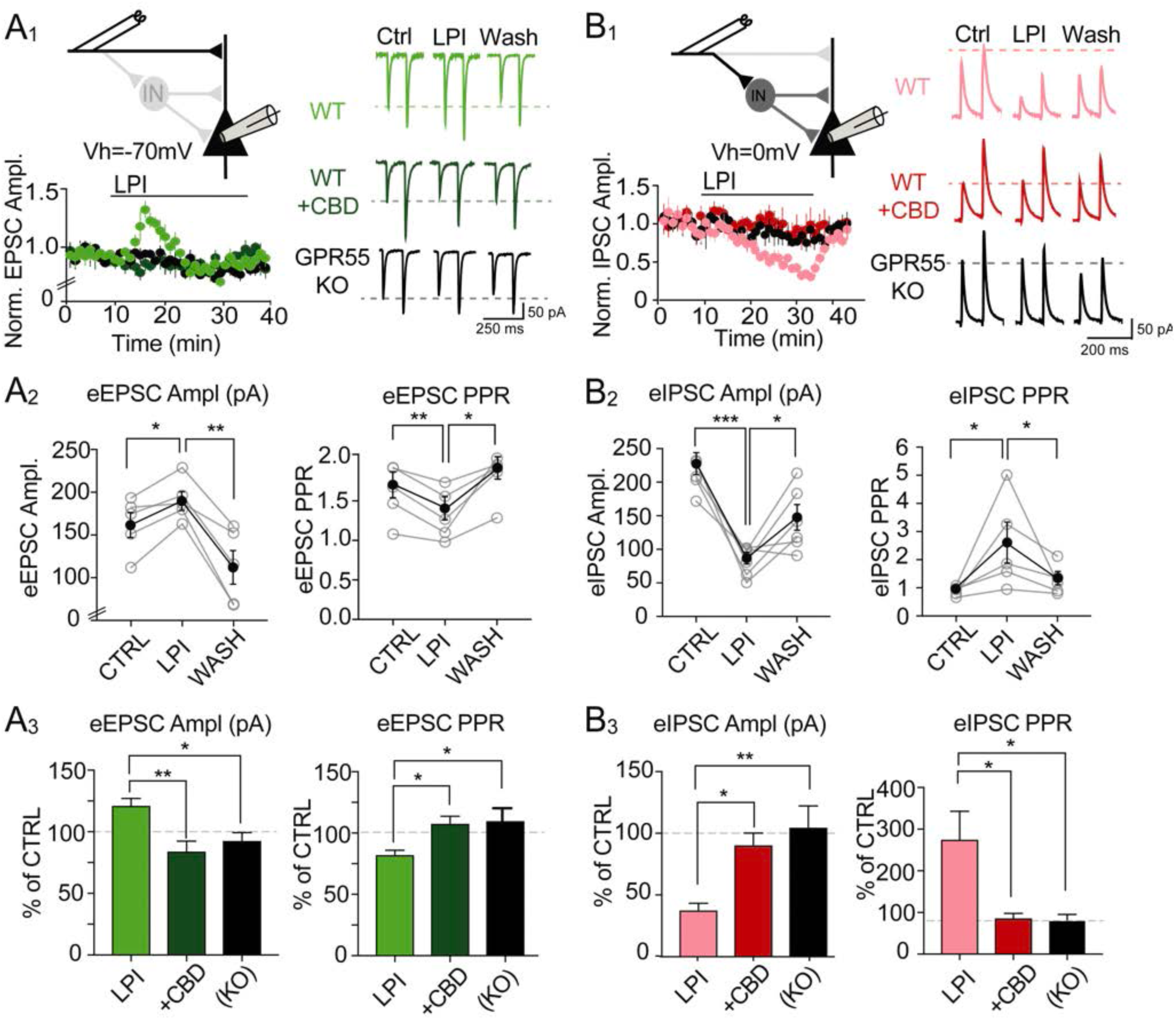
LPI regulates synaptically evoked transmission in the CA3-CA1 microcircuit by augmenting excitation and diminishing disynaptic inhibition. (A_1_) Recordings in CA1 pyramidal neurons (V_hold_ = −70 mV), with stimulating electrodes placed in the *s. radiatum* to activate Schaffer Collateral axons. LPI (4 μM) caused a transient, reversible elevation in synaptically evoked EPSCs (eEPSCs) in WT (light green) slices, blocked by pre-treatment with CBD (dark green), and absent in GPR55 KO mice (black). (A_2-3_) LPI elevated eEPSC amplitude (p=0.021 LPI vs CTRL, p=0.0039 LPI vs WASH, 5 min window, 5 min post LPI application), that corresponded with a drop in the paired pulse ratio (PPR) between the second and first events (p=0.0092 LPI vs CTRL, p=0.020 LPI vs WASH). The LPI-mediated rise in eEPSC amplitude and decrease in PPR was blocked by 1 μM CBD (Ampl: p=0.0057, PPR: p=0.042), and deficient in GPR55 KO mice (Ampl: p=0.028, PPR: p=0.042). (B_1_) To isolate inhibitory currents, CA1 neurons were held at ∼E_Glu_=0 mV. Schaffer collateral stimulation evoked presumed disynaptic (E→I→E) inhibitory currents (Suppl. Fig 4C). LPI produced a gradual reduction in eIPSC amplitude (pink), prevented by pre-treatment with CBD (red), and deficient in slices from GPR55 KO mice (black). (B_2-3_) eIPSC amplitude was significantly lowered by application of LPI (p=0.0006 CTRL vs LPI, p=0.032 LPI vs WASH, 5 min window, 25 min post LPI). LPI also elevated the PPR during the same time period (p=0.039). LPI-mediated effects were impeded by CBD pre-treatment (Ampl: p=0.013, PPR: p=0.021), and absent in GPR55 KO mice (Ampl: p=0.0017, PPR: p=0.013).

We next asked how LPI affects feedforward inhibition, focusing on the disynaptic circuit in CA1 (CA3-derived Schaffer collaterals→ interneurons→ CA1 pyramidal cells). Inhibitory synaptic currents (evoked IPSCs, eIPSCs) were isolated by holding CA1 pyramidal cells at the reversal potential for glutamatergic transmission (E_exc_ = ∼0 mV), stimulating Schaffer Collaterals, and recording outward GABAergic currents (Fig. 3B_1_, 10×10Hz). Application of glutamatergic synaptic blockers (10 μM NBQX + 50 μM APV) abolished eIPSCs, confirming that the outward events were disynaptic (Suppl. Fig. 4C: p=0.0051). After 25 min of exposure, LPI strikingly reduced eIPSC amplitude (Fig. 3B_1-3_: baseline 223±16 pA vs LPI 82±9 pA, n=6, p=0.0006) by 60-70%, while dramatically elevating the eIPSC PPR (baseline 0.9±0.1 vs LPI 2.5±0.7, n=6, p=0.039). The LPI-mediated decrease in eIPSC amplitude was prevented by 1 μM CBD (Fig. 3B_3_, Suppl. Fig. 4D: n=4, p=0.013) and was absent in GPR55 KO slices (n=5 KO, p=0.0017). Furthermore, the increase in eIPSC PPR was blocked by CBD (n=4, p=0.021) and not apparent in GPR55 KO slices (n=5, p=0.013). The disparity between early transient elevation of EPSC amplitude (Fig. 3A_1_) and the slowly developing attenuation of IPSC amplitude (Fig. 3B_1_) resembled the contrast between time courses of mEPSC frequency (Fig. 2B_1_) and mIPSC modulation (Fig. 2B_2_). A similar pattern was also found in recordings of compound (E+I) synaptic currents (n=9, Suppl. Fig. 4A), corroborating observations of EPSCs or IPSCs in isolation.

Taken together, our experiments showed clear effects of LPI on evoked synaptic excitation and, for the first time, feedforward inhibition. LPI initiated an early rise in evoked excitatory transmission due to elevated release probability, followed by a dramatic reduction in inhibitory synaptic strength after 20-30 min post-exposure. In combination, the LPI effects strongly shifted the excitatory-to-inhibitory ratio of the CA3→CA1 microcircuit towards hyperexcitability. Both actions of LPI were eliminated by CBD pre-treatment or by genetic deletion of GPR55.

### LPI-CBD opposition at E→I and I→E connections along the disynaptic pathway

Recognizing that LPI and CBD strongly modulate overall feedforward inhibition, we dissected the LPI sensitivity of individual synaptic elements in the disynaptic circuit (Fig. 4). Parvalbumin-positive (PV+) interneurons served as a principal target based on their primary role in feedforward inhibition and gating of CA1 pyramidal cell firing (Hu et al., 2014; Pouille and Scanziani, 2001). GPR55 was more strongly colocalized with tdTomato-labeled PV+ interneurons than with tdTomato-labeled SST+ interneurons or CCK+ interneurons (E.C. Rosenberg and E. Nebet, unpublished). We targeted tdTomato-labeled PV+ neurons in the CA1 pyramidal layer of slices from PV-Ai9 mice (Fig. 4A_1_, Suppl. Fig. 5A, male *B6;129P2-Pvalb^tm1(cre)Arbr^/ B6.Cg-Gt(ROSA)26Sor^tm9(CAG-tdTomato)Hze^*), verifying that these inhibitory neurons exhibited high frequency spiking patterns and brief membrane time constants (Fig. 4A_1_, Suppl. Fig. 5A). In voltage clamp recordings from PV+ neurons, LPI elevated the frequency of mEPSCs after 5 min drug exposure, but not in the presence of CBD (Suppl. Fig. 5B). LPI also increased Schaffer Collateral-evoked EPSC amplitude (Fig. 4A_1-3_: baseline 120±25 pA vs LPI 162±27 pA, n=5, p=0.0007), and decreased the eEPSC PPR (Fig. 4A_2_: baseline 1.6±0.2 vs LPI 0.8±0.1, n=5, p=0.022). Notably, pre-treatment with CBD (Fig. 4A_3_, Suppl. Fig. 5D) prevented the ability of LPI to increase eEPSC amplitude (LPI 150±19% of baseline vs CBD+LPI 78±7%, n=5 for both conditions, p=0.035), and to reduce eEPSC PPR (LPI 55±10% of baseline vs CBD+LPI 103±4%, n=5 for both conditions, p=0.0020). These data indicated that LPI enhanced release probability at Schaffer Collateral synapses onto PV+ interneurons, an effect abolished by CBD, in line with some colocalization of GPR55 with Vglut1 near PV+ somata (Suppl. Fig. 5C), and consistent with observations at excitatory synapses onto pyramidal neurons (Fig. 3A).

**Fig. 4:**
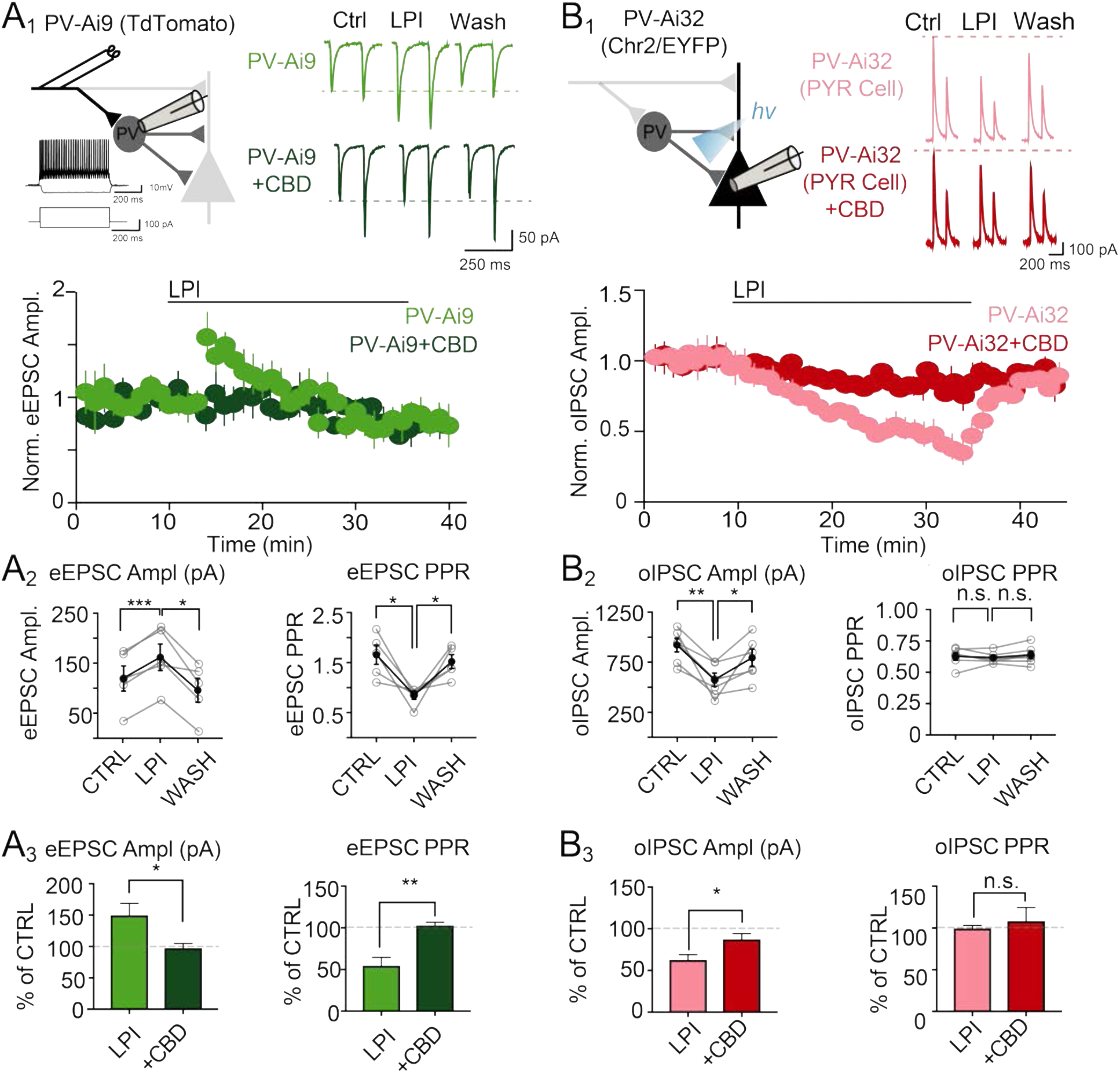
LPI transiently strengthens inputs onto PV neurons while attenuating their inhibitory output. (A_1-3_) Recording in genetically labeled PV+ interneurons (PV-Cre x Ai9), 4 μM LPI elevated the Schaffer collateral-evoked (e) EPSC amplitude (light green, p=0.0007 vs baseline), reversible during the wash period (p=0.012). Concurrently, LPI reduced the PPR (p=0.022 vs baseline; p=0.031 vs wash). CBD pre-treatment prevented the effects of LPI on amplitude (dark green, p=0.035) and PPR (p=0.0020). Analysis was performed in a 5 min window, 5 min post LPI application. (B_1-3_) LPI reduced the amplitude of optogenetically evoked monosynaptic IPSCs (oIPSCs) in slices from PV-Cre x Ai32/ChR2 mice (pink, n=6, p=0.0052 vs baseline, p=0.039 vs wash), but did not change the PPR (p=0.71). CBD blocked effects of LPI on IPSCs (red, p=0.028). Analysis was performed over a 5 min window, 25 min post LPI application.

To isolate the effect of LPI on I→E, PV+ interneuron synapses onto pyramidal neurons, we used slices from PV-Cre/Ai32/ChR2 mice that expressed channelrhodopsin in PV+ terminals (Fig. 4B_1_, male *B6;Cg-Gt(ROSA)26Sor^tm32(CAG-COP4*H134R/EYFP)Hze^/ B6;129P2-Pvalb^tm1(cre)Arbr^*). Blue light activation of PV fibers in the CA1 region of the hippocampus (10×10Hz) caused an outward, optically-evoked IPSC (oIPSC). Exposure to LPI for 25 min reduced the monosynaptic IPSC amplitude (Fig. 4B_1-3_: baseline 909±68 pA vs LPI 563±66 pA, n=6, p=0.0052), without a significant change in the oIPSC PPR (baseline 0.62±0.03 vs LPI 0.62±0.02, n=6, p=0.71). Pre-treatment with CBD blocked the effects of LPI on oIPSC amplitude (Fig. 4B_3_, Suppl. Fig. 5E: LPI 63±6% of baseline vs CBD+LPI 87±7%, n=5 CBD+LPI, p=0.028), also with no net effect on oIPSC PPR (Suppl. Fig. 5E: LPI 100±4% of baseline vs CBD+LPI 108±17%, n=5 CBD+LPI, p=0.59). Thus, LPI reduces the monosynaptic impact of PV+ interneurons likely through a postsynaptic effect, thereby contributing to the overall reduction in disynaptic inhibition onto CA1 pyramids. The time course and magnitude of the LPI-mediated attenuation of I→E contribute to the overall reduction in disynaptic inhibition, although a role for altered PV+ neuron excitability might also reflect direct effects on ion channel conductances (Ghovanloo et al., 2018; Kaplan et al., 2017; Khan et al., 2018; Patel et al., 2016; Zhang and Bean, 2021).

### LPI gradually attenuates clustering of GABA_A_R γ_2_ and gephyrin at inhibitory postsynaptic sites

To clarify the molecular underpinnings of LPI and CBD actions at GABAergic inhibitory synapses, we examined potential signaling targets by combining immunocytochemical and biochemical approaches. GABA_A_Rs are hetero-pentameric protein complexes whose activity-dependent trafficking to postsynaptic synapses is enhanced by inclusion of the γ_2_ subunit (Jacob et al., 2005) and anchored by the scaffolding protein gephyrin (Jacob et al., 2008). We immunostained these proteins in hippocampal cell cultures to track fine-scale synaptic changes. Sprague-Dawley rat cultures (both sexes) were exposed to 4 μM LPI, fixed 30 and 60 min later, and imaged with MAP2 as a dendritic marker. Relative to vehicle treatment controls, LPI significantly reduced the intensity of GABA_A_R γ_2_ puncta at both 30 and 60 min post application (Fig. 5A: normalized intensity relative to vehicle condition: VEH 100±2%, n=10 neurons; LPI-30 min 72±3%, n=7; LPI-60 min 78±5%, n=7; one-way ANOVA: p<0.0001, VEH vs LPI-30 p=0.0001, VEH vs LPI-60 p=0.0003). The LPI-mediated decrease in GABA_A_R γ_2_ puncta intensity reversed after 10 minutes of ACSF washout (Suppl. Fig. 6A: p=0.014 relative to LPI). The density of GABA_A_R γ_2_ puncta per unit area of dendrite was reduced at 60 min but not at 30 min post-LPI (Suppl. Fig. 6B: LPI-60 vs VEH p=0.029).

**Fig. 5:**
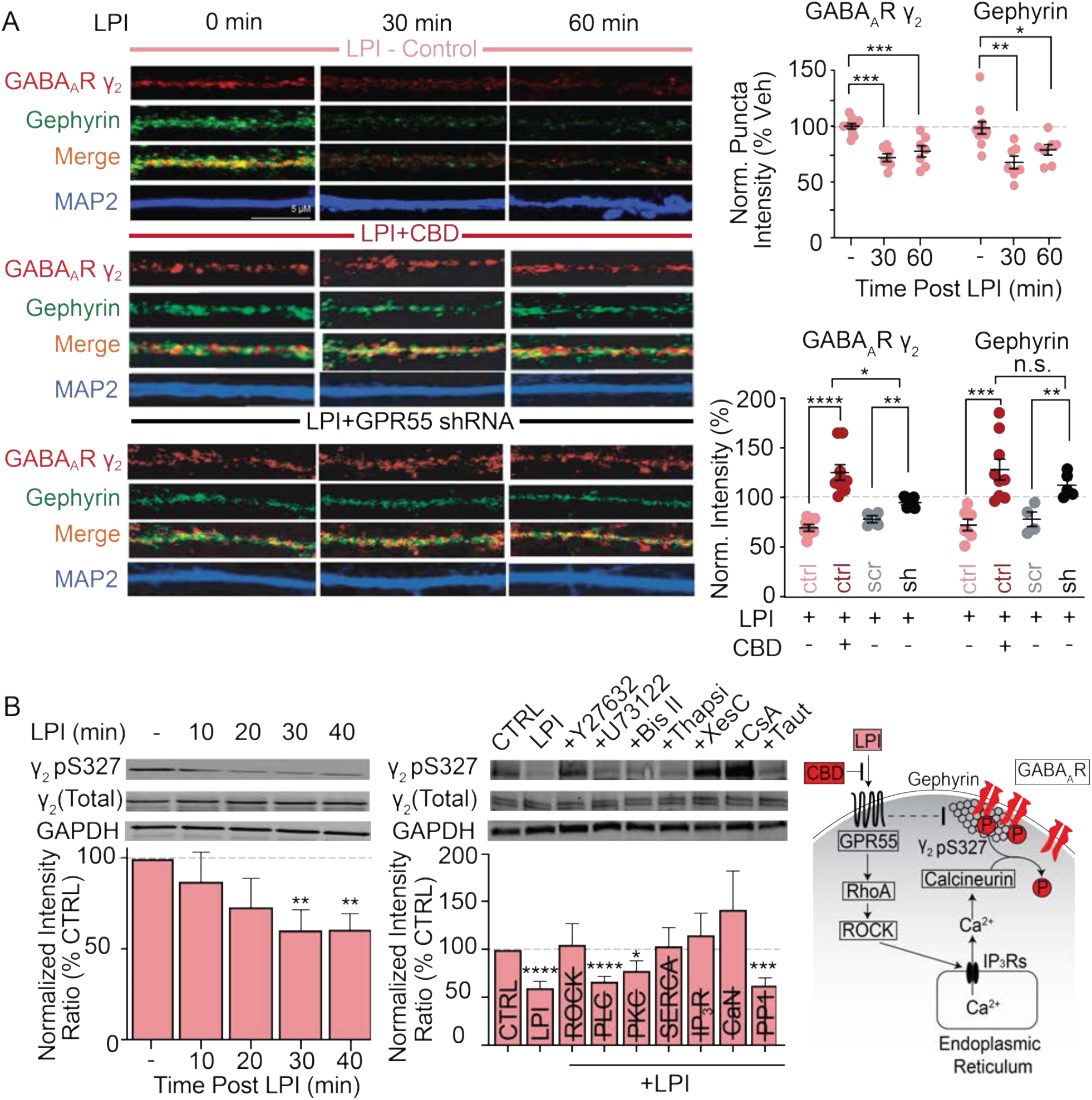
LPI mediates a GPR55-dependent, progressive downregulation of GABA_A_R γ_2_ and gephyrin clustering, via S327 dephosphorylation. (A) 4 μM LPI reduced the intensity of GABA_A_R γ_2_ puncta at 30- and 60-min post-application (one-way ANOVA with post-hoc Dunnett’s multiple comparisons: VEH vs LPI-30 p=0.0001, VEH vs LPI-60 p=0.0003). LPI also diminished the intensity of gephyrin at 30 and 60 min post-LPI (VEH vs LPI-30 p=0.0011, VEH vs LPI-60 p=0.032). 1 μM CBD (red) prevented both LPI-mediated decreases in GABA_A_R γ_2_ (p<0.0001) and gephyrin (p=0.0007) puncta at 30 min. Similarly, lentiviral shRNA-mediated knockdown of GPR55 (black, sh) deterred effects of LPI on GABA_A_R γ_2_ (p=0.0052) and gephyrin (p=0.0064) in comparison to scrambled (scr) shRNA controls (gray). CBD co-application with LPI produced a higher γ_2_ level than LPI+shRNA alone (p=0.017). (B) LPI reduced (γ_2_ phospho-S327)/(total γ_2_ expression) in whole cell hippocampal lysates (phospho-/total γ_2_ as % of VEH, LPI-30 min 61±11% p=0.0099; LPI-40 min 61±9%, p=0.0029). LPI-mediated S327 dephosphorylation was reversed by inhibitors of RhoA-activated protein kinase (ROCK) (10 μM YM-27632), IP_3_Rs (1 μM xestospongin C), and calcineurin (10 μM cyclosporin), as well as depletion of intracellular calcium stores via 10 μM thapsigargin. However, LPI-mediated effects remained robust in the presence of inhibitors of phospholipase C (10 μM U73122), protein kinase C (0.1 μM bisindolylmaleimide II), and PP1α (5 nm tautomycetin). Right, biochemical scheme based on literature, consistent with pharmacological data shown.

A similar pattern was observed for gephyrin: LPI diminished the intensity of gephyrin puncta at 30 and 60 min post-LPI (Fig. 5A: VEH 100±6%, n=10; LPI-30 69±6%, n=7; LPI-60 80±5%, n=7; one-way ANOVA p=0.0016: VEH vs LPI-30 p=0.0011, VEH vs LPI-60 p=0.032). Following a 10 min wash with ACSF, gephyrin intensity increased to greater than baseline levels (Suppl. Fig. 6A: VEH 100±4%; LPI-30 min 74±5%; WASH 135±22%; one-way ANOVA p=0.0002: VEH vs LPI p=0.028, LPI vs WASH p=0.0002, VEH vs WASH p=0.026). LPI reduced gephyrin puncta density only at 60 min post-treatment (Suppl. Fig. 6B: VEH vs LPI-60 p=0.042). The percentage of receptor puncta colocalized with gephyrin remained unchanged after applying LPI (Suppl. Fig. 6C-D), suggesting that gephyrin and γ_2_ intensity were reduced in parallel. Collectively, these data support a scenario wherein LPI diminishes both GABA_A_Rs and the associated gephyrin scaffold in synaptic puncta (consistent with a reduction in mIPSC amplitude, Fig. 2B_2_), before reducing puncta density and thus contributing to a reduction in mIPSC frequency (Fig. 2B_2_).

Consistent with our findings with synaptic transmission (Fig. 3), pre-treatment of cultures with 1 μM CBD prevented the LPI-mediated reduction in GABA_A_R γ_2_ intensity (Fig. 5A: normalized intensity relative to condition-matched VEH treated coverslips: LPI 72±3%, n=7 vs CBD+LPI 128±8% n=8, p=<0.0001; p=0.016 CBD+LPI vs no LPI baseline) and GABAR γ_2_ puncta density per unit area (Suppl. Fig. 6E: p=0.0001 CBD+LPI vs LPI; p=0.014 CBD+LPI vs no LPI baseline). Furthermore, 1 μM CBD blocked the LPI-driven decrease in gephyrin intensity (Fig. 5A: LPI 69±6%, n=7 vs CBD+LPI 124±10%, n=8, p=0.0007; p=0.72 CBD+LPI vs baseline) and gephyrin puncta per unit area (Suppl. Fig. 6E: p=0.0005 CBD+LPI vs LPI; p=0.096 CBD+LPI vs baseline).

To test whether LPI effects on inhibitory synaptic proteins were mediated by GPR55, we transfected cell cultures with GPR55-targeted shRNA and verified that GPR55 protein expression was significantly reduced by shRNA (Suppl. Fig. 6F: relative GPR55 intensity: untransfected 100±10%, n=8 vs shRNA 57±6%, n=5; p=0.034), but not with scrambled shRNA controls (91±12%, n=6; p=0.89). LPI failed to reduce GABA_A_R γ_2_ intensity in cultures transfected with GPR55 shRNA (Fig. 5A: normalized intensity relative to condition-matched VEH treated coverslips: LPI-scr 81±4%, n=5 vs LPI-shRNA 97±3%, n=5; p=0.0052) and γ_2_ puncta density (Suppl. Fig. 6G: p=0.0033). Similarly, shRNA knockdown of GPR55 blocked LPI-mediated decreases in gephyrin intensity relative to scrambled shRNA controls (Fig. 5A: LPI-scr 74±7%, n=5 vs LPI-shRNA 108±5%, n=5; p=0.0064) and in gephyrin puncta per unit area (Suppl. Fig. 6G: p=0.017). Comparing the effects of CBD and GPR55 shRNA, CBD co-application with LPI (Fig. 5A, middle right, red) produced a higher γ_2_ level than LPI+shRNA alone (black, p=0.017). Together, these observations indicate that LPI decreased the postsynaptic expression of GABA_A_R γ_2_ and its scaffolding partner gephyrin within 30 min, with reduced puncta density after 60 min. These actions were mediated by GPR55 and blocked by CBD.

### Possible signaling mechanisms to link GPR55 to GABA_A_R density

A potential biochemical control point for LPI effects on GABA_A_R γ_2_ postsynaptic localization is phosphorylation of γ_2_ serine 327, a key regulator of γ_2_ clustering. Activity-dependent dephosphorylation of γ_2_ S327 increases lateral mobility of GABA_A_Rs away from the gephyrin scaffold and destabilizes inhibitory synapses (Garcia-Morales et al., 2015; Muir et al., 2010). Exposure to LPI gradually reduced phosphorylation of γ_2_ S327 in western blots of whole cell lysates, reaching significance at 30 and 40 min post LPI application (Fig. 5B: LPI-30 min 61±11% phospho-/total γ_2_ as % of VEH, p=0.0099; LPI-40 min 61±9% of VEH, p=0.0033; n=8 for all). In contrast, LPI did not significantly alter β_3_ S408/409 phosphorylation (Suppl. Fig. 6H), an activity-dependent regulator of GABA_A_R endocytosis (Kittler et al., 2005), suggesting that LPI effects on GABA_A_R might be subunit-specific. Treatment of cultures with 1 μM CBD blocked the LPI-mediated decrease in GABA_A_R γ_2_ S327 dephosphorylation (Suppl. Fig. 6I: VEH 100±3%; LPI 60±8%; LPI+CBD 122±20%; one-way ANOVA p=0.0030: VEH vs. LPI p=0.047, LPI vs. CBD+LPI p=0.0027, VEH vs. CBD+LPI p=0.50).

We tested a potential signaling mechanism linking GPR55 to γ_2_ S327 dephosphorylation. GPR55 is a G_q_-linked (Lauckner et al., 2008) or G_α12/13_-linked (Henstridge et al., 2009; Ryberg et al., 2007) GPCR; when signaling via G_α13_, GPR55 communicates to RhoA and Rho-Associated Protein Kinase (ROCK) to release intracellular Ca^2+^ from internal stores (Ross, 2009). Blockade of ROCK with YM-27632 (10 μM) prevented the LPI-mediated dephosphorylation of γ_2_ S327 (Fig 5B: LPI+ YM-27632, 105±21% vs vehicle, n=4 for all hereafter, p=0.76). LPI–induced S327 dephosphorylation remained robust with blockade of PLC (LPI+10 μM U73122 67±5%, p<0.0001) and PKC (LPI+0.1 μM bisindolylmaleimide II 78±10%, p=0.024). However, LPI γ_2_ S327 dephosphorylation was blocked by depleting intracellular Ca^2+^ stores with thapsigargin (10 μM; LPI+Thapsi 104±19% vs vehicle, p=0.76), inhibition of IP_3_Rs with xestospongin C (1 μm; 115±22%, p=0.41), and reduced calcineurin activity (10 μM cyclosporine A, 142±40%, p=0.22), but not by PP1α inhibition (5 nM tautomycetin, 63±7%, p=0.0002). Jointly, these findings suggest that LPI, communicating via GPR55 and ROCK, signals in a PLC-independent manner to release intracellular Ca^2+^ stores, activate calcineurin, and dephosphorylate GABA_A_R γ_2_ S327, thus dispersing GABA_A_R clusters.

### Seizures elevate GPR55 expression and potentiate pro-excitatory effects of LPI

If LPI-GPR55 signaling shifts the E:I ratio towards hyperexcitability, potentially favoring seizure activity, could seizures reciprocally regulate the LPI-GPR55 axis? To address this, we assessed the expression of GPR55, ∼30 min following acute PTZ-induced seizures in WT mice. Injection of PTZ (105 mg/kg, i.p.) caused a prominent, region-specific elevation in GPR55 expression (Fig. 6A,B), with the highest increases in area CA1 (Non-Seizure 100±22% vs PTZ+VEH 204±34%, p=0.031) and CA3 (Non-Seizure 100±17% vs PTZ+VEH 221±42%, p=0.044), and a non-significant elevation in DG (Non-Seizure 100±22% vs PTZ+VEH 152±36%, p=0.30; n=6 for all conditions, 2 slices from each of 3 animals). *In vivo* pre-treatment with CBD (200 mg/kg, i.p.), 1 h prior to PTZ seizure induction completely prevented the seizure-induced rise in GPR55 expression (Fig. 6A, B: CA1 PTZ+CBD 82±23% GPR55 expression relative to non-seizure, p=0.0069 vs PTZ+VEH; CA3 PTZ+CBD 99±29%, p=0.034 vs PTZ+VEH). To crosscheck immunostaining data, we probed levels of hippocampal *Gpr55* mRNA by quantitative PCR using newly designed primers that showed specificity in KO controls and no associated genomic DNA contamination (Suppl. Fig. 7A,B). Assessment of *Gpr55* mRNA by qPCR supported immunostaining data, with a 60% increase in transcripts following PTZ treatment (Fig. 6C: Non-Seizure+VEH 1.0±0.1, n=7 vs PTZ+Veh 1.6±0.2, n=7; p=0.021), fully reversed at the least by CBD pre-treatment (PTZ+CBD 0.19±0.1, n=4; p<0.0001 vs PTZ+VEH; p=0.0060 vs Non-Seizure+VEH). Thus, immunocytochemistry and qPCR both showed that seizures elevate GPR55 expression within an hour following seizure induction, effects blocked by CBD pre-treatment.

**Fig. 6:**
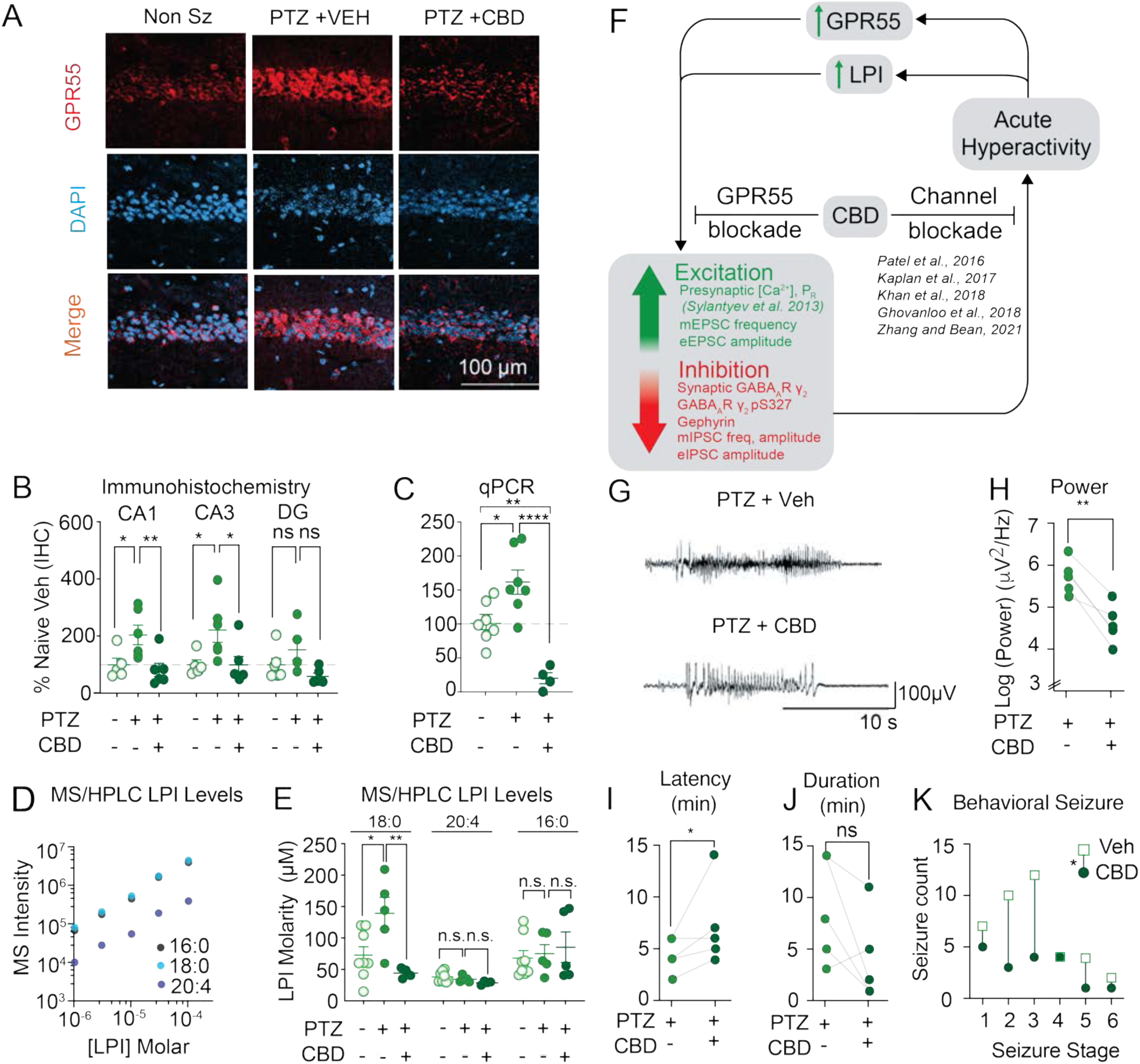
Acute seizures elevate GPR55 expression and increase levels of 18:0 LPI, prevented by CBD pre-treatment. (A-B) Seizures induced by pentylenetetrazole (PTZ, 105 mg/kg, i.p.) mediated a prominent, region-specific elevation in GPR55 expression, with the greatest increases in area CA1 (p=0.031, representative images) and CA3 (p=0.044). The seizure-induced elevation in GPR55 was blocked by *in vivo* pre-treatment with 200 mg/kg, i.p. CBD, 1 h prior to PTZ seizure induction (CA1 p=0.0069 PTZ+VEH vs PTZ+CBD; CA3 p=0.034 PTZ+VEH vs PTZ+CBD). (C) PTZ seizures increase GPR55 mRNA expression, prevented by 200 mg/kg, i.p. CBD administration 1 h prior to PTZ injection (ANOVA p<0.0001: Non-Seizure+VEH vs PTZ+VEH p=0.020, PTZ+VEH vs PTZ+CBD p<0.0001). CBD treated animals had a significantly lower GPR55 mRNA expression in comparison to non-seizure, vehicle controls (p=0.0060). (D) Linear relationship in standard control curves for 3 LPI isoforms (16:0 black triangles, 18:0 light blue squares, 20:4 purple circles) across increasing concentrations of LPI. See Materials and Methods for more details. (E) PTZ significantly elevated levels of the 18:0 isoform of LPI, as assayed by HPLC-MS (one-way ANOVA p=0.0086: Non-Seizure+VEH vs PTZ+VEH p=0.031, PTZ+VEH vs PTZ+CBD p=0.0097). (F) Simplified depiction of major points of this study. Proposed scenario wherein CBD interrupts a hyperactivity-induced LPI-GPR55 positive feedback loop. LPI promotes excitability by a dual mechanism: stimulating transient glutamate release and restructuring the inhibitory postsynapse to downregulate the strength of inhibition. The net effect of LPI-induced changes is to shift the E:I ratio towards acute hyperexcitability and elevate evoked principal cell firing. Seizures in turn upregulate GPR55 and its agonist, LPI. As an intervention, CBD strategically blocks the GPR55-dependent pro-excitatory and anti-inhibitory effects of LPI, and dampens spike firing via direct effects on ion channels (Ghovanloo et al., 2018; Kaplan et al., 2017; Khan et al., 2018; Patel et al., 2016; Zhang and Bean, 2021) thus inhibiting a positive feedback loop that helps drive hyper-excitability. (G-K) CBD action on electrographic seizures. Representative EEG traces (G) and quantification demonstrating that CBD (100 mg/kg, i.p.) administered 1 hour prior to PTZ (60 mg/kg, i.p.)-induced seizures reduces electrographic seizure average power (H, p=0.0036), increases latency to first electrographic seizure (I, p=0.040), and produces a non-significant trend towards reduced EEG seizure duration (J, p=0.11). (K) CBD decreases behaviorally observed PTZ-induced seizures at Racine stages 1-6 relative to matched vehicle treated controls (paired t-test, p=0.047).

To assay levels of LPI following seizures, we used liquid chromatography-mass spectrometry targeting various LPI isoforms (Fig. 6D-E and Suppl. Fig. 7C, D, Suppl. Table 1). The level of 1-stearoyl LPI (C18:0), the predominant LPI component in brain (Oka et al., 2009), was increased 30 min post-PTZ-induced seizure (ANOVA p=0.0086, Non-Seizure+VEH 73±13 μM, n=8 vs PTZ+VEH 139±25 μM, n=5; p=0.031), an elevation prevented by CBD pre-treatment 1 h prior to PTZ administration (PTZ+CBD 44±4 μM, n=4; p=0.0097 vs PTZ+VEH, p=0.51 vs Non-Seizure+VEH). Levels of other LPI isoforms (20:4 and 16:0) remained unchanged. Our findings of hyperactivity-induced elevations in GPR55 and its endogenous ligand raise the possibility of a positive feedback loop that upregulates excitatory synapses and downregulates inhibitory ones (Fig. 6F). The synaptic effects could provide an autocatalytic mechanism to promote circuit excitability that becomes dangerous in the context of overactivity. The resulting elevation of GPR55 expression, and LPI levels, would drive enhancement of excitation and attenuation of inhibition, promoting hyperexcitability. By directly antagonizing LPI activation of GPR55, CBD would counteract LPI’s pro-excitability action, lower GPR55 and LPI levels as observed, and thus hold the positive feedback in check (Fig. 6F). Additionally, CBD could also dampen excitability through a GPR55-independent mechanism such as a direct effect on excitatory ion channels (Ghovanloo et al., 2018; Patel et al., 2016; Zhang and Bean, 2021), which in turn could restrain activity-dependent GPR55 gene expression and LPI release (Sylantyev et al., 2013) (Fig. 6F). These putative mechanisms, operating at different stages of the overall loop, would be inherently complementary. In either case, we predict that CBD would act strategically to attenuate electrographic seizure activity.

To assess the role of CBD in electrographic seizures, we employed a PTZ model as previously described (Vilela et al., 2017), in a refinement of our earlier protocol (Fig. 1 C-D, Suppl. Fig. 1). Here we placed electrodes to capture a larger spatial range of electrographic activity in the frontal, temporal (hippocampus), and occipital lobes, and used a lower concentration of PTZ (60 mg/kg) as a milder stimulus that would not cause status epilepticus. We found that pre-treatment with 200 mg/kg CBD decreased average EEG power (Fig. 6G,H; Supplemental Fig. 7E, F: log(power): PTZ + VEH 5.7±0.2, n=5 vs PTZ + CBD 4.6±0.2, n=5; paired t-test p=0.0036), increased the latency to first electrographic seizures (Fig. 6I, PTZ + VEH 4.4±0.7 min vs PTZ + CBD 7.2± 0.8 min; p=0.04), and gave a non-significant trend towards abbreviation of EEG seizures (Fig. 6G,J, PTZ + VEH 8.8±2.3 min vs PTZ + CBD 4.0±1.9 min; p=0.11). These findings demonstrated that CBD reduces electrographically recorded seizures in a manner consistent with our working hypothesis (Fig. 6F). In the same animals, CBD also limited behaviorally observed seizures rated over Racine stages 1-6 (Fig. 6K, p=0.047).

### LPI-GPR55 signaling enhanced in a model of temporal lobe epilepsy

We next asked if a seizure-induced rise in GPR55 would potentiate the impact of LPI in a chronic model of epilepsy, in which epileptogenesis occurs for months following induction of status epilepticus. We used a low mortality, high morbidity model of lithium-pilocarpine (Li-PLC)-induced *status epilepticus* in male Wistar-Kyoto rats (Modebadze et al., 2016), in which seizure burden is reduced by CBD delivery during the epileptogenic period (Patra et al., 2019) (protocol outlined in Fig. 7A; see Materials and Methods). Following behaviorally-verified induction of status epilepticus, epileptogenesis was determined using a previously validated post-seizure behavioral battery (PSBB) test, and only animals with PSBB scores >10 following a 10-week period were used for qPCR, immunohistochemistry, and electrophysiology (Modebadze et al., 2016). At 3-6 months after Li-PLC induced epileptogenesis, there was a hint of an increase in *Gpr55* mRNA (55±29%) relative to non-epileptic age-matched controls (Fig. 7B, n=9 non-epileptic controls, n=11 epilepsy + vehicle) that failed to reach statistical significance (p=0.19), possibly due to experimental variability. There was no significant difference in *Gpr55* mRNA expression in animals treated with chronic CBD (200 mg/kg *p.o.*) following establishment of epileptogenesis vs non-epileptic controls or epileptic animals + vehicle. (Fig. 7B, epileptic + CBD 1.0±0.2 relative to non-seizure controls, n=7). However, GPR55 protein expression, as assessed by IHC at 20x resolution, roughly doubled in CA1, CA3, and DG following Li-PLC epileptogenesis (Fig. 7C: CA1 control 100±28%, n=9 vs Li-PLC 217±43%, n=7, p=0.020; CA3 control 100±26%, n=9 vs Li-PLC 220±49%, n=8, p=0.028; DG control 100±23%, n=8 vs Li-PLC 178±16%, n=7, p=0.014). Oral (*Per orem, p.o.*) intake of CBD 200 mg/kg during the weeks following confirmation of epileptogenesis completely prevented GPR55 elevation in all three areas (Fig. 7C: CA1 Li-PLC+CBD *p.o.* 119±28%, n=12, p=0.042 vs Li-PLC; CA3 Li-PLC+CBD *p.o.* 113±18%, n=11, p=0.042 vs Li-PLC; DG Li-PLC+CBD *p.o.* vs Li-PLC 99±14%, n=10, p=0.0096). In CA1 *stratum radiatum*, a locus of synaptic inputs to CA1 neurons, the normalized intensity of GPR55 colocalized with excitatory nerve terminals marked by Vglut1 was enhanced ∼4-fold following Li-PLC induction (Fig. 7D: control 100±17%, n=8 vs Li-PLC 405±111%, n=8; p=0.006), but not if CBD was given orally following the period of epileptogenesis (Li-PLC+CBD *p.o.* 159±30%, n=10; p=0.022). A similar pattern was seen in *stratum pyramidale* (Suppl. Fig. 8A) where axons originating in area CA3 also course (Ropireddy et al., 2011). Li-PLC epileptogenesis also induced a 27±8% increase in GPR55 puncta intensity at presumed postsynaptic inhibitory sites labeled with gephyrin in *stratum pyramidale*, an effect prevented by chronic *p.o.* CBD treatment (Fig. 7E, ANOVA p=0.0007; control 100±20%, n=7 vs Li-PLC + Veh 127±8%, n=6, p=0.04; Li-PLC+CBD 82±5%, n=9, p=0.0004 vs Li-PLC +Veh). A similar effect was seen in the *stratum radiatum* (Suppl. Fig. 8B). The intensity of GPR55 puncta colocalized with Vgat, a presynaptic marker of inhibitory synapses, was nearly doubled following Li-PLC administration (Suppl. Fig. 8C, D), effects prevented by CBD *p.o.* in S.R. (Suppl. Fig. 8C), but not in S.P. (Suppl. Fig. 8D). At the same time, levels of the major LPI isoforms (18:0, 20:4, and 16:0) neither rose nor fell following Li-PLC epileptogenesis (Suppl. Fig. 8E-F, Suppl. Table 1, ANOVA comparing control, Li-PLC, Li-PLC+CBD changes in each isoform: 18:0 p=0.20; 20:4 p=0.45, 16:0 p=0.76). In hippocampal tissue, the concentration of 18:0 LPI [LPI] was ∼0.1-1 μM in rat, roughly similar to prior reports of rat [LPI] (Oka et al., 2009), while roughly 100 to 1000-fold lower than [LPI] detected in mouse (10-100 μM, Fig. 6D-E, Suppl. Table 1). Taken together, our data provide evidence that levels of GPR55 protein, but not the agonist LPI, were elevated several months following Li-PLC induced epileptogenesis; these increases were prevented by chronic treatment with oral CBD.

**Fig. 7:**
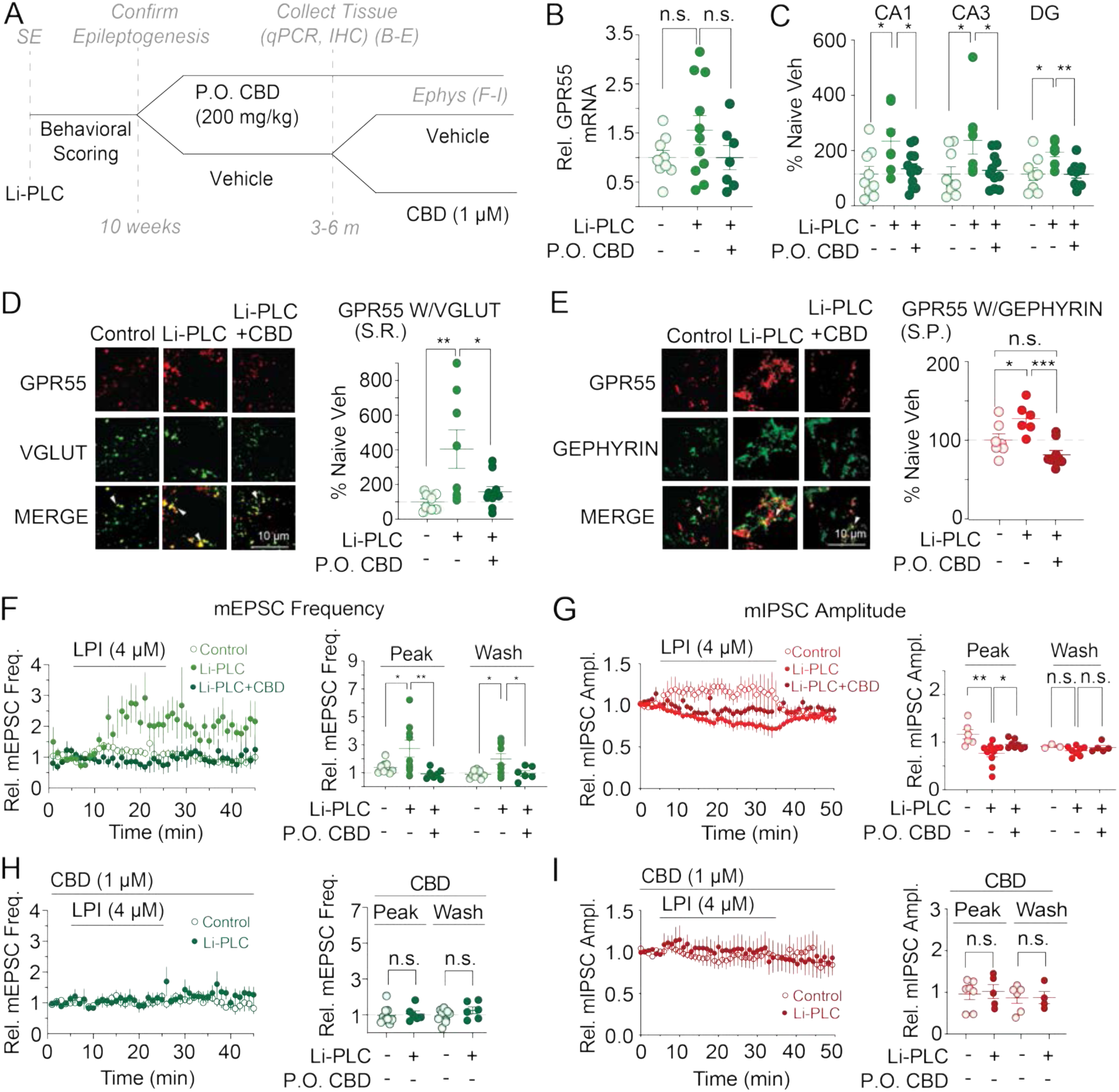
CBD prevents Li-Pilocarpine (Li-PLC) epileptogenesis-induced potentiation of GPR55 protein expression and enhanced LPI activity. (A) Experimental paradigm for Li-PLC epileptogenesis (see Methods and (Modebadze et al., 2016; Patra et al., 2019)). (B) Li-PLC epileptogenesis slightly elevates hippocampal GPR55 mRNA expression but without reaching statistical significance (ANOVA p=0.1907). (C) Hippocampal GPR55 protein expression (IHC, 20x resolution) elevated in *ex vivo* slices from rats following Li-PLC epileptogenesis in area CA1 (p=0.020 vs age-matched, non-seizure controls), CA3 (p=0.028), and DG (p=0.014). CBD, administered 200 mg/kg *p.o.* after the period of epileptogenesis, prevents the rise in GPR55 expression in CA1 (p=0.042), CA3 (p=0.042), and DG (p=0.0096). (D) Li-PLC epileptogenesis increases intensity of GPR55 puncta colocalized with Vglut1 at putative presynaptic excitatory terminals in the CA1 *stratum radiatum* (S.R., 63x), prevented by chronic *p.o.* CBD treatment. Left: representative images, Right: ANOVA p=0.0069: control vs Li-PLC p=0.0063, Li-PLC vs Li-PLC+CBD *p.o.* p=0.020). (E) GPR55 intensity increases at putative post-synaptic sites of inhibitory synapses labeled with gephyrin (S.R., 63x), following Li-PLC-induced epileptogenesis, prevented by CBD treatment after the period of epileptogenesis. Left: representative images, Right: one-way ANOVA p=0.0007: control vs Li-PLC p=0.035, Li-PLC vs Li-PLC+CBD *p.o.*, p= 0.0004). (F) Pro-excitatory effects of the GPR55 agonist LPI were potentiated in *ex vivo* slices from rats following Li-PLC-induced epileptogenesis (filled light green circles) in comparison to non-epileptic controls (unfilled light green circles), effects prevented by chronic *in vivo* treatment with CBD *p.o.* (filled dark green circles). Vertical axis, normalized values relative to pre-LPI baseline; mEPSC freq: (ANOVA p=0.0099: control vs Li-PLC p=0.038; Li-PLC + CBD *p.o.* vs Li-PLC p=0.0087). The potentiated effects of LPI also persisted following washout (ANOVA p=0.01: control vs Li-PLC p=0.012; Li-PLC + CBD *p.o.* vs Li-PLC p=0.034) (G) LPI produced a greater anti-inhibitory response in mIPSC amplitude in epileptic (filled light red circles) vs. non-epileptic controls (unfilled light red circles), effects prevented by *in vivo* CBD *p.o.* treatment (filled dark red circles, ANOVA p=0.0032: control vs Li-PLC p=0.0017; Li-PLC + CBD *p.o.* vs Li-PLC 0.047). (H-I) Acute CBD (1 μM) blocked the effects of LPI on both mEPSC frequency (H) and mIPSC amplitude (I) in both Li-PLC slices and non-epileptic controls.

The pro-excitatory effects of exogenous LPI, examined in acute slices taken ∼3 months after induction of recurrent seizures, were strongly potentiated relative to non-epileptic controls (Fig. 7F). The LPI-mediated rise in mEPSC frequency was greater in slices from Li-PLC-treated animals, an increase prevented by chronic *in vivo* treatment with CBD 200 mg/kg given orally (*p.o.*) following epileptogenesis (Fig. 7F: ANOVA 0.0099: control 1.4±0.1-fold relative to baseline, n=9 vs Li-PLC 2.7±0.6, n=9, p=0.038; Li-PLC + CBD *p.o.* 0.9±0.1, n=7 vs Li-PLC p=0.0087). The potentiated effects of LPI also persisted for at least 20 min following washout (Fig. 7F: ANOVA p=0.01: control 0.88±0.1 vs Li-PLC 2.0±0.4, p=0.012; Li-PLC + CBD *p.o.* 0.9±0.2 vs Li-PLC p=0.034). The mEPSC amplitude remained unchanged throughout (Suppl. Fig. 8G-H, ANOVA p=0.26). LPI produced a greater dis-inhibitory response in mIPSC amplitude in epileptic vs. non-epileptic controls, effects prevented by *in vivo* CBD *p.o.* treatment (Fig. 7G: ANOVA 0.0032: control 1.2±0.1x, n=6 vs Li-PLC 0.8±0.1x, n=10; p=0.0017; Li-PLC + CBD *p.o.* 0.95±0.03, n=8 vs Li-PLC 0.047), whereas LPI did not diminish mIPSC frequency (Suppl. Fig. 8I-J, ANOVA p=0.95). Each of the effects of LPI on excitatory and inhibitory currents were completely blocked by *ex vivo* acute pre-treatment of slices with 1 μM CBD (mEPSCs, Fig. 7H: control+LPI+CBD, n=9 vs Li-PLC+LPI+CBD, n=6: p=0.76; mIPSCs, Fig. 7I: control+LPI+CBD, n=7 vs Li-PLC+LPI+CBD, n=5: p=0.77). Taken together, our data indicate that Li-PLC epileptogenesis increases GPR55 protein expression, potentiates synaptic responses to LPI, and prolongs the effects of LPI on mEPSC frequency. All of these post-epilepsy changes were abolished by orally delivered CBD (Fig. 7F,G), in accord with block of the Li-PLC-induced elevation of GPR55 (Fig. 7D). This is as expected if the hypothetical positive feedback loop (Fig. 6F) operated *in vivo*.

### CBD prevents “two pulse” potentiation of KA-induced seizures

For more precise temporal control and assessment of biochemical changes and their impact on seizure susceptibility, we examined GPR55 changes 4 and 48 h following injection with kainic acid, a chemoconvulsant model of temporal lobe seizures (Iyengar et al., 2015) (Materials and Methods). We chose a ‘two-pulse’ protocol to determine if a peri-threshold dose for seizures, slightly below that required to consistently elicit seizures, might lead to reliable seizures if delivered again days later. We hypothesized that the first dose (“KA1”) might initiate a measurable rise in GPR55, as predicted by the proposed positive feedback loop (Fig. 6F), which could in turn lower the seizure threshold for an identical KA dose days after the first (“KA2”).

Indeed, hippocampal GPR55 mRNA expression increased by 70% at 48 h following the first largely subconvulsive dose of kainic acid (KA1, 24 mg/kg, s.c.) (Fig. 8A: ANOVA p=0.0176; post-hoc comparisons: non-seizure+VEH1 1.0±0.1, n=8 vs KA1+VEH1 1.7±0.3, n=8; p=0.039). In contrast, no significant change was seen 4 h after KA1 (1.2±0.1, n=4, p=0.86). Administration of CBD (200 mg/kg) 1 h prior to KA1 prevented the rise in GPR55 mRNA at 48 h (0.9±0.1, n=5, p=0.039 vs KA1+VEH1). In the same animals used for qPCR, there was no significant difference in KA1 seizure incidence following administration of CBD1 vs VEH1 (3/6 animals each, p=0.11). Consistent with qPCR findings, IHC revealed an increase in GPR55 intensity in hippocampal CA1 and CA3, 48 hours post KA1 administration, effects prevented by CBD treatment 1 hour prior to KA1 (Fig. 8B: n=11 for all, CA1 ANOVA p= 0.02, Non Sz 100±9% vs KA1 142±11% p=0.040, KA1+CBD1 101±14% vs KA1 p=0.048; CA3 ANOVA p<0.0001 Non Sz 100±6% vs KA1 158±12% p=0.0006, KA1+CBD1 91±12% vs KA1 p= 0.0001; DG ANOVA p=0.09).

**Fig. 8:**
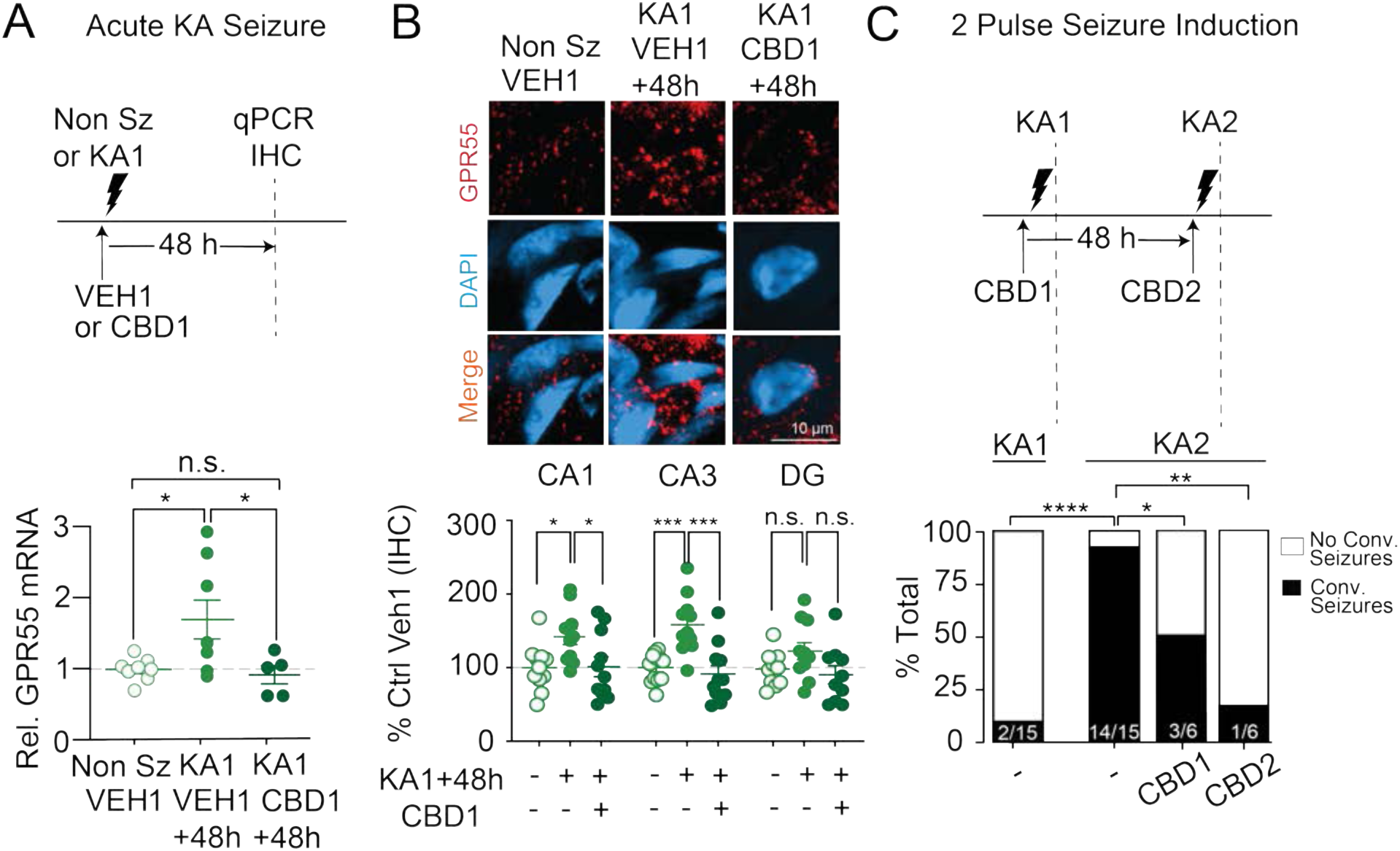
CBD decreases “second-dose” kainic acid (KA)-induced seizures. (A) GPR55 mRNA expression was significantly increased 48 h following kainic acid injection (KA1, 24 mg/kg s.c.) relative to vehicle-treated, non-seizure controls (ANOVA p=0.017: Non-Seizure+VEH vs KA+VEH1 p=0.039). However, pre-treatment with CBD (200 mg/kg, s.c.) 1 h prior to KA1 injection prevented the rise in GPR55 mRNA at 48 h (p=0.039 relative to KA1+VEH1). (B) GPR55 protein increases in CA1 and CA3 hippocampus 48 hours post KA1 administration, effects prevented by CBD 200 mg/kg 1 hour prior to KA1 (CA1 ANOVA p=0.02: Non Sz vs KA1 p=0.040, KA1 vs KA1+CBD1 p=0.048; CA3 ANOVA p<0.0001: Non Sz vs KA1 p=0.0006, KA1 vs KA1+CBD1 p= 0.0001; DG ANOVA p=0.09). (C) Effects of 2^nd^ dose of KA administered 48 h post KA1 (KA2, 24 mg/kg, s.c.). Behavioral seizures assessed via modified Racine Scale 2 h after KA injection (time of assay denoted by dotted vertical lines), followed by injection with diazepam 10 mg/kg, s.c. While 2/15 animals demonstrated convulsive seizures (as defined as 3-4 on Racine Scale) following KA1, a significantly greater proportion (14/15, p<0.0001, Fisher’s Exact Test) displayed KA-induced seizures after the KA2 challenge. CBD (200 mg/kg, s.c.) reduced the frequency of convulsive seizures following KA2 when CBD was administered 1 h prior to KA1 injection (CBD1, p=0.05) or 1 h prior to KA2 (CBD2, p=0.0017). There was no significant difference between KA1 seizure incidence with CBD1 or VEH1 administration (p=0.11, data not shown).

In light of the elevated GPR55 expression 2 days after the initial KA challenge, we tested if a second KA dose (KA2) would induce more convulsive seizures as hypothesized (Fig. 6F). Indeed, a greater proportion of animals exhibited tonic-clonic seizures after the second KA dose (KA2, 14/15) than after the initial KA pulse (KA1, 2/15, p<0.0001, Fig. 8C). CBD administration reduced the incidence of second pulse seizures when given 1 hr prior to the second KA2 pulse (CBD2: 1/6, p=0.0017), potentially due to an acute anti-seizure effect of CBD. However, CBD also reduced KA2 seizure incidence when given prior to the first priming, subconvulsive KA1 dose, 48 hours prior to the second KA dose (CBD1: 3/6, p=0.05). The effect of CBD1 on second pulse seizures (48 hours later) is most likely due to an enduring effect on cellular signaling, not an acute dampening of excitability (Fig. 6F) that would readily reverse over the two days prior to KA2, given the *in vivo* half-life of CBD given i.p. (estimated about 4-9 h) (Deiana et al., 2012). Collectively, these results support the hypothesis that seizure-induced increases in GPR55 might lower the threshold for subsequent seizures (Figure 6F), and CBD1 blockade of increased GPR55 (Fig. 8A) could provide one potential mechanism for its anti-seizure action (Fig. 8C). This reasoning leaves open the further possibility of GPR55-independent effects on excitability.

## Discussion

### CBD dampens E:I ratio and opposes a seizure-induced LPI-GPR55-mediated positive feedback loop

Our experiments provide new insights on synaptic mechanisms that contribute to the anti-seizure effects of CBD. We determined that CBD regulates excitability in part by preventing the actions of the GPR55 lipid agonist, LPI, at both excitatory and inhibitory synapses. LPI shifted the excitatory-to-inhibitory ratio towards hyperexcitability via two mechanisms: a fast rise in presynaptic excitatory strength (Sylantyev et al., 2013) followed by a prolonged gradual reduction in inhibitory strength, mediated by lowered clustering of GABA_A_R γ_2_ and its scaffolding partner gephyrin. LPI pushed the circuit towards hyperexcitability by enhancing the probability of synaptically-driven firing of CA1 pyramidal cells, without changing their basal intrinsic firing. CBD blocked these LPI-promoted, GPR55-mediated changes. Further, we discovered that seizures potentiate the GPR55-LPI pathway through elevation of GPR55 expression, increased or at least maintained LPI levels, and exaggerated LPI action. This modulatory but potentially pathogenic positive feedback loop (Fig. 6F, Fig. 8) is strategically interrupted by CBD, likely contributing to its therapeutic action *in vivo* (Fig. 7).

### LPI enhances circuit gain, potentially opposing stabilizing actions of endocannabinoids and LPA

The pro-excitatory effects of LPI and their sensitivity to CBD blockade prompted exploration into the control of endogenous LPI levels. We found that acute PTZ-induced seizures increased levels of the major LPI isoform (18:0) (Oka et al., 2009), as assessed by direct MS/HPLC. This aligns with evidence for activity-dependent enhancement of LPI generation and release: (1) LPI synthesis is partially dependent on secretory phospholipase A2 (PLA2) (Yamashita et al., 2013), a Ca^2+^-dependent enzyme (Burke and Dennis, 2009) whose levels are elevated by seizures (Yegin et al., 2002). (2) PLA2-driven LPI synthesis/release is likely elevated by brief bursts of presynaptic firing, as indicated by the effects of CBD and inhibitors of PLA2 (Sylantyev et al., 2013). (3) Efficacy of CBD action increases with graded neuronal depolarization, consistent with depolarization-induced release of LPI (Khan et al., 2018). Together, these data suggest that stimulus-driven, phasic release of LPI provides a short-lasting signal that in moderation acts to upregulate information transfer through feedforward circuits. However, following seizure-induced elevations of GPR55, the same neuromodulatory control system could be hijacked by a pathological positive feedback loop (Fig. 6F) that magnifies and greatly prolongs the pro-excitatory effect of LPI (Fig. 7F). Supportive genetic evidence for LPI’s pathological potential arises from inactivating mutations in LPI acyltransferase, the enzyme that catalyzes LPI conversion to its PI precursor, which cause epilepsy and intellectual disability in humans (Johansen et al., 2016). Further work is needed to determine the control of LPI synthesis and release, its relevance to other disorders associated with disrupted E:I ratio (Lewis et al., 2012; Rubenstein and Merzenich, 2003), and its possible value as a biomarker of CBD responsiveness. CBD pre-treatment could prevent the activity-dependent elevation in 16:0 LPI if hyperactivity were dampened in either a GPR55-dependent or independent manner (Fig. 6F). We also cannot exclude a direct action of CBD on PLA2 activity, while noting that *in vivo* concentrations of CBD used in this study (200 mg/kg, equivalent to ∼ 10 μM) (Deiana et al., 2012) are >10-fold lower than concentrations that inhibit PLA2 *in vitro* (IC_50_ 134 μm) (Evans et al., 1987; White and Tansik, 1980).

Less is known about the LPI-GPR55 axis than the endocannabinoid (eCB) signaling system, an endogenous synaptic “circuit breaker” that maintains stability despite elevated network activity (Katona and Freund, 2008). Activity-dependent release of eCB compounds (Castillo et al., 2012) activates presynaptic GPCRs, predominantly CB_1_Rs (Kreitzer and Regehr, 2001), that also bind Δ^9^-tetrahydrocannabinol (THC) and other cannabinoids. This retrograde signaling can reduce the presynaptic release of glutamate in depolarization-induced suppression of excitation (DSE) (Llano et al., 1991) or presynaptic release of GABA in depolarization-induced suppression of inhibition (DSI) (Pitler and Alger, 1992). Similarly, a related bioactive lipid, lysophosphatidic acid (LPA) depresses presynaptic excitatory release by reducing docked vesicles at the active zone and reduces inhibitory tone by decreasing postsynaptic GABA_A_R aggregation (Garcia-Morales 2015). We found that LPI complements and contrasts with both eCBs and LPA in its pattern of action at excitatory and inhibitory synapses. Whereas eCBs and LPA generally weaken excitation and inhibition together, LPI reduces postsynaptic inhibitory weights while elevating glutamate release, strongly increasing the E:I ratio as we show here. This combination of actions enables LPI to powerfully affect net synaptic excitation, amplifying incoming signals and elevating excitability. Insofar as eCBs, LPI, and LPA are lipid products of a shared biochemical network (Yamashita et al 2013), and acute seizures elevate levels of eCB (Farrell et al., 2021; Marsicano et al., 2003; Wallace et al., 2003), activity-dependent shifts in enzymatic conversion from 2-AG/LPA to LPI might provide a strategic control point to regulate brain excitability and a potential therapeutic target.

### Evidence that elevated LPI-GPR55 signaling is a potential target of CBD’s anti-seizure action

In this study, we utilized several models of both acute seizures and chronic epileptogenesis to evaluate GPR55 as a potential target for the anti-seizure mechanism of CBD. Using Li-PLC administration in rats, a well-described method for inducing spontaneously recurrent seizures, we demonstrated that GPR55 protein expression is persistently elevated several months following the initial epileptogenic insult (Fig. 7C). GPR55 levels were roughly doubled across multiple hippocampal regions relative to age-matched controls, but not if CBD was given orally during the epileptogenic period, at doses sufficient to reduce chronic seizure burden (Patra et al. 2018). Consistent with elevation of GPR55 at loci identified with excitatory and inhibitory synaptic markers (Fig. 7D,E, Suppl. Fig. 8A-D), respective synaptic responses to exogenous LPI application in slices were strikingly enhanced (Fig. 7F,G). Despite the upregulation of LPI-GPR55 signaling, CBD remained capable of preventing the responses at both excitatory and inhibitory synapses in slices, assessed with sodium channels blocked (Fig. 7H,I). We applied CBD at 1 μM, sufficient to antagonize GPR55 (reported IC_50_ of 445 ± 67 nM) (Ryberg et al., 2007), and in line with peak plasma levels (>1 μM) in patients with Dravet Syndrome receiving CBD long term (20 mg/kg x 21 days) (Devinsky et al., 2018b). Limitations of the rat Li-PLC model are that animals are euthanized several months following initial epileptic insult, agnostic to the last acute seizure, so that transient activity-dependent changes in acute LPI release might escape capture. Indeed, we found that levels of LPI detected in the hippocampus of epileptic rats were not significantly different than non-epileptic controls (Suppl. Fig. 8F). Of note, recent studies suggest that seizure-induced elevation in endocannabinoids are acute, and renormalize within 1 hour post seizure (Farrell et al., 2021; Marsicano et al., 2003; Sugaya and Kano, 2021). This highlights a potential similarity with our findings if post-seizure changes in LPI are similarly fleeting. Furthermore, levels of both *Gpr55* mRNA and [LPI] are significantly lower in rats in comparison to mice (Supplemental Table 1, Fig. 7B), which likely leads to increased variability in the [LPI] assays (e.g. Fig. 7B, Suppl. Fig. 8F), and a less robust effect of LPI on inhibitory mIPSC amplitude (Fig. 7G). Despite this more variable response in some aspects of this model of long-term model epilepsy, we predict that the robust effects of epileptogenesis on GPR55 protein expression (Fig. 7C-E) and synaptic function (Fig. 7F-I) will exert secondary effects on hyperexcitability.

To further explore a mechanistic role for GPR55 in acute seizures, we employed proconvulsants PTZ (Fig. 1, 6), a GABA_A_R antagonist to model disinhibition, and kainic acid (Fig. 8), a GluR agonist, to model temporal lobe seizures. Acute PTZ induction of seizures up-regulated GPR55 at both mRNA and protein levels, 30 min following PTZ injection (Fig. 6B, C), and acutely elevated the 18:0 isoform of LPI (Fig. 6E). Further, GPR55 mRNA increased 48 h following a single dose of kainic acid, in accord with an elevation in “second pulse” seizure susceptibility at this time (Fig. 8). Seizure-induced increases in GPR55 expression were prevented by CBD pre-treatment at a dose that yielded anti-seizure effects *in vivo* (200 mg/kg, Fig. 1C). A relatively higher dose of CBD was selected due to multiple factors including pharmacokinetics following acute delivery, rapid distribution of lipid soluble CBD into hydrophobic tissue, and a species difference in brain access from the circulation, evident in attenuation of brain:plasma concentration ratio in mice relative to other species (see: Epidyolex EMA Assessment 2019)(Deiana et al., 2012).

CBD’s anti-convulsive effects on PTZ-induced seizures were absent in GPR55 KO mice (Fig. 1C), consistent with GPR55’s role in seizure generation and CBD antagonism of GPR55 (Fig. 6F). A potential confound is that PTZ-induced seizures were more prevalent in GPR55 KO mice than expected in comparison to peak CBD suppression in WT animals (Fig. 1C). This difference might arise from the absence of GPR55 during development, when LPI-GPR55 signaling contributes to axon pathfinding and synapse maturation (Cherif et al., 2015; Guy et al., 2015). Hints of possible compensatory effects were evident in unitary excitatory synaptic weights, which were greater in KO than WT slices (Suppl Fig. 2A). Future studies should examine conditional deletion of GPR55 later in development and seizure susceptibility of GPR55 KO mice in additional models of acute seizure (e.g. 4-AP, kindling) and chronic epilepsy.

Our findings complement and extend earlier *in vitro* data demonstrating that CBD (1) blocks LPI-mediated increases in excitatory synaptic weights (Sylantyev et al., 2013) and (2) upregulates intrinsic firing of PV+ interneurons and thus spontaneous IPSC frequency (Kaplan et al., 2017; Khan et al., 2018). We found that CBD can extinguish a potentially regenerative loop in which hyperactivity enhances LPI-GPR55 signaling, further shifting the E:I ratio (Fig. 6F). The schema also alludes to additional non-GPR55-dependent actions, such as a role for CBD reducing excitability via direct effects on ion channels (Ghovanloo et al., 2018; Patel et al., 2016; Zhang and Bean, 2021). We also note that CBD reduced seizure-induced mortality in both WT and GPR55 KO mice (Fig. 1D) and exerted effects on GABA_A_Rs above that of GPR55 knockdown alone (Fig. 5A), suggesting potential direct effects of CBD on GABA_A_Rs (Bakas et al., 2017). Further candidates for CBD’s anti-seizure effects (Gray and Whalley, 2020; Ibeas Bih et al., 2015) include agonism of transient receptor potential (TRP) channels (TRPV1, TRPV2, TRPA1) (Bisogno et al., 2001; Costa et al., 2004; De Petrocellis et al., 2011; Qin et al., 2008), negative allosteric modulation of CB_1_Rs (Laprairie et al., 2015; Straiker et al., 2018), as well as inhibition of the ENT-1 transporter of adenosine (Carrier et al., 2006), CaV3.3 channels (Ross et al., 2008), mitochondrial VDAC1 channels (Rimmerman et al., 2013), and regulation of non-neuronal cells (e.g. glia). In addition, GPR55 could also play a non-synaptic role in regulating ion channel conductances in interneurons (Khan et al., 2018), especially PV+ interneurons (Chamberland et al., 2019; Kaplan et al., 2017; Khan et al., 2018) through effects of Na_V_ channels as listed above. We propose that CBD might act on non-GPR55 targets to regulate network excitability, opposing potential stimuli that “engage” the hyperactivity-induced feedback loop as described in Fig. 6F. However, effects on such non-GPR55 targets that rely on the immediate presence of CBD are unlikely to explain partial efficacy against KA seizures that persisted 48 h after animals are given CBD (Fig. 8C), long after drug levels in the brain have decayed. Nonetheless, these candidate targets call for further evaluation at clinically relevant doses of CBD.

### LPI mediates, and CBD prevents, a reduction in inhibitory synaptic weight: a mechanism for CBD synergy with GABA_A_R modulation

We found that the effects of LPI at inhibitory synapses were absent in GPR55 KO slices, prevented by CBD, and dependent on intracellular calcium stores. In these respects, inhibitory synapses resembled their excitatory counterparts. However, in contrast to GPR55-mediated enhancement of excitatory transmission, which was presynaptically expressed and strikingly transient (Sylantyev et al., 2013) (Figs. 2, 3), GPR55 activation downregulated inhibitory postsynaptic strength over a much slower time course. The reduction of inhibitory postsynaptic weights developed gradually, over 25-30 min post-LPI application, and would easily be missed with briefer recordings. The differences in synaptic locus and dynamics focus attention on underlying distinctions in mechanism.

GPR55 activation reduced phosphorylation of GABA_A_R at γ_2_ S327, a well-known phosphorylation site that regulates γ_2_ lateral movement from the gephyrin scaffold to extrasynaptic sites (Garcia-Morales et al., 2015; Jacob et al., 2005; Muir et al., 2010). We observed a slow, progressive decrease in mIPSC amplitude and frequency reflected by reduced GABA_A_R γ_2_ puncta intensity and γ_2_ S327 phosphorylation 30 min post application and decreased GABA_A_R synapse density at 60 min. Our data align with findings that slow γ_2_ dephosphorylation by calcineurin supports a form of inhibitory LTD (Lu et al., 2000; Wang et al., 2003), and that NMDAR stimulation causes gradual dephosphorylation of γ_2_ S327 and GABA_A_R dispersal (Muir et al., 2010). We also observed that S327 dephosphorylation depended on calcineurin, as well as RhoA/ROCK kinase and IP_3_R-mediated release of calcium from intracellular stores (Garcia-Morales et al., 2015; Muir et al., 2010). However, in contrast to previous instances of mediation by phospholipase C (Henstridge et al., 2009), we found that GPR55-triggered dephosphorylation was not prevented by PLC blockade (Fig. 5B). As an alternative to the canonical pathway wherein PIP_2_ cleavage drives IP_3_ formation, ROCK can also activate IP_3_Rs by direct interaction (Mehta et al., 2003). This would drive Ca^2+^ release and CaN activation in a chain of events consistent with our observations.

Beyond effects on γ_2_, LPI also reduced the number and intensity of gephyrin puncta; gephyrin scaffolding favors the synaptic localization of GABA_A_R γ_2_-containing heteromers (Essrich et al., 1998; Kneussel et al., 1999). Immunolabeled puncta of γ_2_ and gephyrin were diminished together, suggesting that LPI-induced changes in γ_2_ and gephyrin operate coordinately to disperse inhibitory complexes and thus decrease mIPSC frequency, inhibitory weights, and IPSC amplitude. The biochemical link from GPR55 to gephyrin provides an area for future investigation. In one scenario, GPR55 activates RhoA/ROCK and thus regulates actin (Maekawa et al., 1999), which in turn modulates gephyrin positioning or stability (Bausen et al., 2006; Giesemann et al., 2003; Hanus et al., 2006). Alternatively, GPR55 might provoke kinases (ERK) or phosphatases (CaN) to alter gephyrin phosphorylation (Tyagarajan and Fritschy, 2014). This might mobilize the same pathway for dispersion of gephyrin + GABA_A_R clusters as found for elevated synaptic input (Bannai et al., 2009).

GPR55-mediated changes in GABAergic signaling will contribute strongly to regulation of E:I ratio, promoting information transfer but also potentially exacerbating seizure pathophysiology. GABA_A_R γ_2_, the target of GPR55 action, has previously been implicated in both clinical and preclinical studies of seizures. Surface levels of γ_2_ are decreased in patients with temporal lobe epilepsy (Loup et al., 2000) and in hippocampal tissue of rodents displaying Li-PLC-induced status epilepticus (Goodkin et al., 2008). Reduced surface γ_2_ and GABAergic current occur with γ_2_ point mutations that cause pediatric epilepsy with febrile seizures (Baulac et al., 2001; Frugier et al., 2007; Wallace et al., 2001). This suggests that LPI-mediated reduction in levels of γ_2_ and gephyrin could predispose brain circuits to a proconvulsive state, phenocopying acquired and genetic mutations in γ_2_ seen in epilepsy. Further, LPI/GPR55 signaling could lower the ceiling on GABAergic synaptic strength and thereby limit the maximal impact of positive allosteric modulators (PAMs) of GABA_A_R, which are notoriously ineffective in many forms of epilepsy and status epilepticus (Burman et al., 2022). By opposing GPR55 activation and restoring the abundance of GABA_A_R clusters, CBD might remove both functional and therapeutic limitations of reduced benzodiazepine efficacy in chronic seizures, thus contributing to its clinical efficacy in combination therapy with GABA_A_R PAMs (Anderson et al., 2019; Chuang et al., 2021).

## Author Contributions

E.R., S.C., M.B., B.W., H.S., and R.T. designed the research. E.R., S.C., M.B., E.N., X.W., S.M., S.J., S.G., M.W., A.S., S.B., P.P., R.R., performed experiments and analyzed results. N.C. provided imaging analysis code. S.S. provided graphical abstract and figure edits. E.R. and R.T. wrote the paper, with contributions from S.C., M.B., D.J., G.B., O.D., G.W., H.S., and B.W.

## Acknowledgements

We thank Ken Mackie for generously providing the GPR55 KO (*B6;129S-Gpr55^tm1Lex^/Mmnc*) mice. We gratefully acknowledge GW Research Ltd, Cambridge, UK for furnishing the cannabidiol (Epidiolex ®) used in the study. We thank Pablo Castillo, Jayeeta Basu, and Niels Ringstad for valuable comments on an early draft of this manuscript. Additional analysis code was generously provided by Benjamin Suutari, Natasha Tirko, and Katherine Eyring. We thank the NYU Metabolomics Core and Experimental Pathology Research Laboratory Core (both supported by the Cancer Center Support Grant P30CA016087 at NYU Langone’s Laura and Isaac Perlmutter Cancer Center) for their help in acquiring and analyzing the data presented in this report. This work is supported by funding from the Ruth L. Kirschstein National Research Service Awards (NRSA) for Individual Pre-doctoral MD/PhDs (F30 NS100293), the NYU MSTP Training Grant (T32GM007308), and grants to RWT from the NIMH (5R37MH071739), NIDA (DA040484-01), the Simons Foundation, and the *Vulnerable Brain Project*. S.C., A.S., and O.D. supported by funding from FACES, Finding a Cure for Epilepsy & Seizures. S.C. supported by a Charles H. Revson Senior Fellowship in Biomedical Science and a postdoctoral fellowship from the Fonds de Recherche du Québec - Santé (FRQS) and a K99/R00 Pathway to Independence Award from NIMH (1K99MH126157-01).

