## Supplemental Figures and Methods for "Cannabidiol targets a modulatory system for excitatory-inhibitory synaptic coordination, contributing to its anti-seizure action"

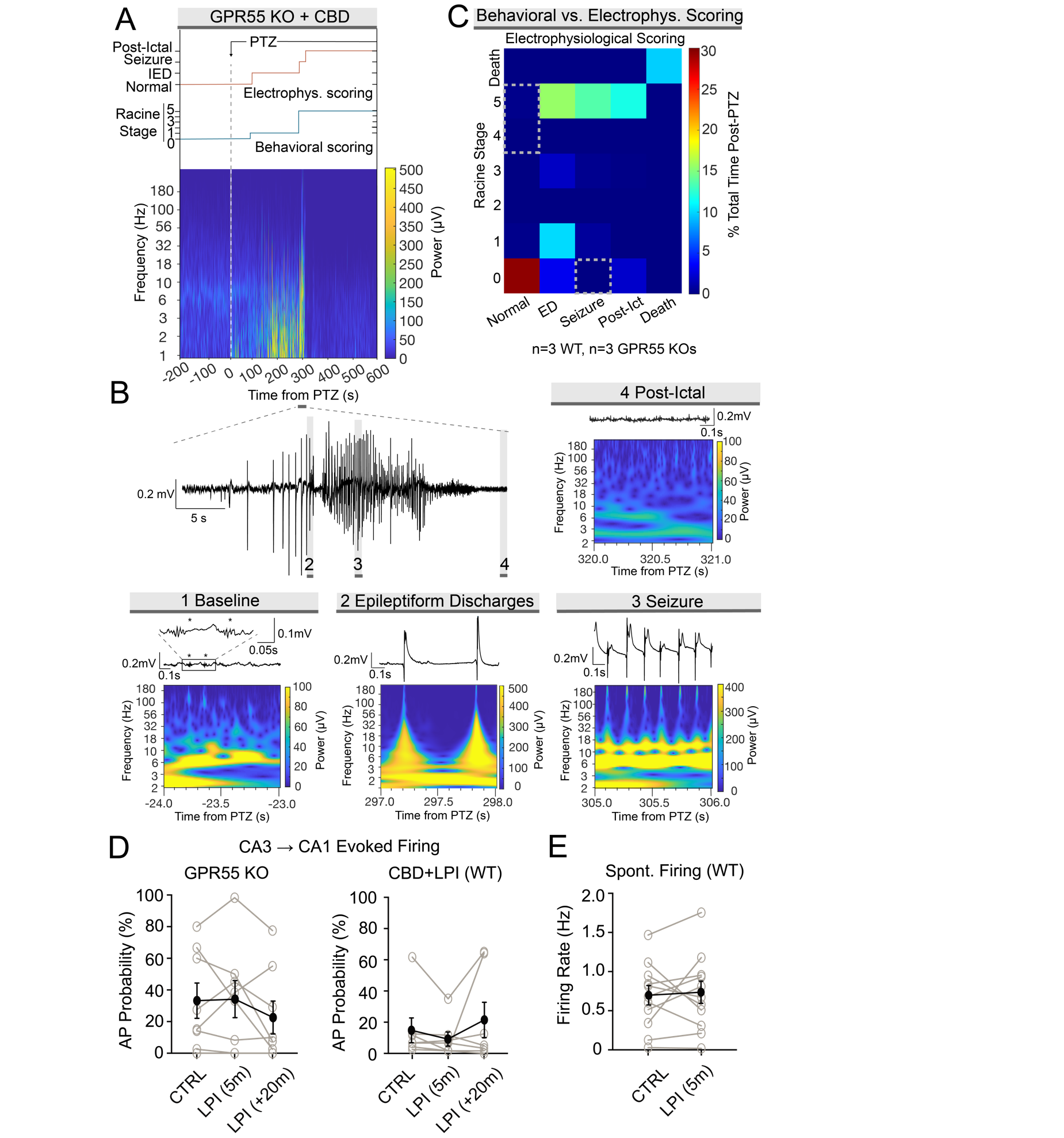
***Supplemental Figures***

**Suppl Fig. 1, Related to Fig. 1: Effects of CBD on *in vivo* pentylenetetrazole (PTZ)-induced seizures and *ex vivo* evoked and spontaneous spiking** (A) Robust correlation between electrographic and behaviorally observed PTZ (105 mg/kg, i.p.)-induced seizures in a representative GPR55KO mouse treated with CBD 200 mg/kg, i.p. 1h prior to PTZ induction. (Top) orange line: electrographic seizures as recorded with tungsten electrodes targeted at hippocampal area CA1 were classified as “Normal” (low amplitude baseline), “ED” (epileptiform discharges), “Seizure” (rhythmic, or semi-rhythmic, high amplitude population spikes), or “Post-ictal” (depressed signal following seizure activity). (Middle) blue line: seizures were scored according to Racine scale (see methods, (Pohl and Mares, 1987)). (Bottom): spectrogram demonstrating EEG power and frequency during recording. Time zero indicates PTZ i.p. injection, as indicated by vertical dotted gray line. (B) Representative electrographic traces and spectrograms of EEG recordings from 290-320 s post-PTZ injection in spectrogram in (A), designated by expansion of horizontal black bar. “Baseline” recordings isolated from -24-23 s prior to PTZ injection, demonstrating sharp wave ripples as indicated by asterisks. (C) Contingency plot comparing behavioral scoring (Racine scale, y-axis) with electrographic seizure scoring (x-axis) from n=3 WT+CBD and n=3 KO+CBD animals treated with PTZ. Heat plot demonstrates % of total time post-PTZ occupied by each condition. Of note, no animals with behaviorally observed tonic-clonic seizures (Racine stage 4-5) had “Normal” EEG scoring (dotted gray rectangle), and no animals with “Seizure” electrographic scoring had behavioral Racine stage 0 (dotted gray square). (D) LPI did not modulate CA3 → CA1 evoked firing in slices from GPR55 KO mice (p=0.89, n=8) or in slices pre-treated with CBD (p=0.19, n=7). (E) LPI did not change the rate of spontaneous firing in slices from WT mice (p=0.74, n=11).

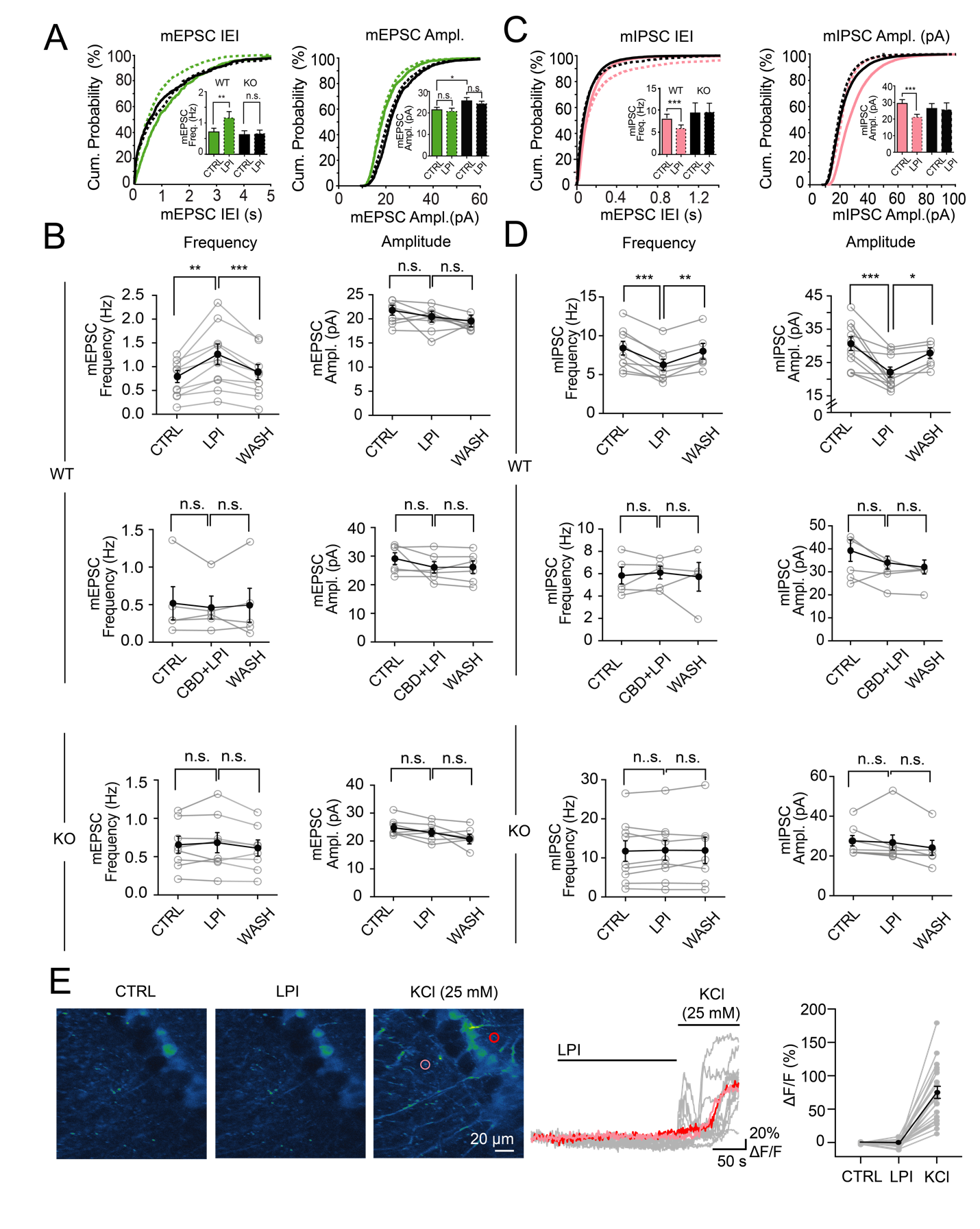

**Suppl Fig. 2, Related to Fig. 2: LPI elevates excitatory synaptic strength and decreases inhibitory strength via GPR55, effects blocked by CBD** (A, left) LPI (4 μM) shifted the population distribution of mEPSC interevent intervals (IEI) leftward, consistent with an elevation in average mEPSC frequency in WT animals (p=0.0026 vs. baseline, n=10), but not KO animals (p=0.48, n=8). WT-basal = solid green line, WT-LPI = dashed green line, KO-basal = solid black line, KO-LPI = dashed black line. (A, right) LPI did not significantly alter mEPSC amplitude in either WT (p=0.47) or KO (p=0.10) mice. However, in genotype comparison, mEPSCs from KO mice were significantly larger than those from WT mice (p=0.018). mEPSC “LPI” events averaged during a 5 min window, 5 min post LPI application. (B) LPI acutely elevated mEPSC frequency (p=0.0026), an effect reversed upon LPI removal (p=0.0008). However, this effect was blocked by pre-treatment with 1 μM CBD (n=5, p=0.91 vs. baseline), and absent in GPR55 KO mice (p=0.50 vs. baseline). (C, left) LPI shifted the distribution of mIPSC IEIs to the right, in line with a reduction in average mIPSC frequency in WT animals (p=0.0006, n=9), but not KO (p=0.78, n=9). WT-basal = solid pink line, WT-LPI = dashed pink line, KO-basal = solid black line, KO-LPI = dashed black line. (C, right) LPI treatment drove the mIPSC amplitude distribution to the left, indicating a reduction in mIPSC amplitude in WT (p=0.0008, n=11), but not KO (p=0.66). There was no significant difference in average mIPSC amplitude between WT and KO animals (p=0.36). (D) LPI curtailed both mIPSC frequency (p=0.0006) and amplitude (p=0.0008), effects that washed out post-LPI removal (Freq: p=0.0015, Ampl: 0.042). CBD prevented the effect of LPI (n=5, Freq: p=0.57, Ampl: p=0.85), as did deletion of GPR55 in the KO animal (Freq: p=0.64, Ampl: p=0.66). mIPSC “LPI” events averaged during a 5 min window, 25 min post LPI application. (E) (Left) Representative images of CA3🡪CA1 hippocampal GCaMP6f Ca^2+^ responses in PV+ interneuron terminals from acute slices of PV-Cre x Ai-148 mice. Slices were perfused with LPI (4 μM, 3 min), followed by KCl (25 mM, 1 min) as a positive control to assess intact axonal Ca^2+^ flux in terminals. (Middle) Ca^2+^ responses (ΔF/F) from ROIs (pink and red), as indicated in representative images on the left. (Right) Pooled data showing that LPI (4 μM) had no effect on presynaptic Ca^2+^ measured in PV-expressing inhibitory terminals, while high [K] stimulation caused significant Ca^2+^ elevation.

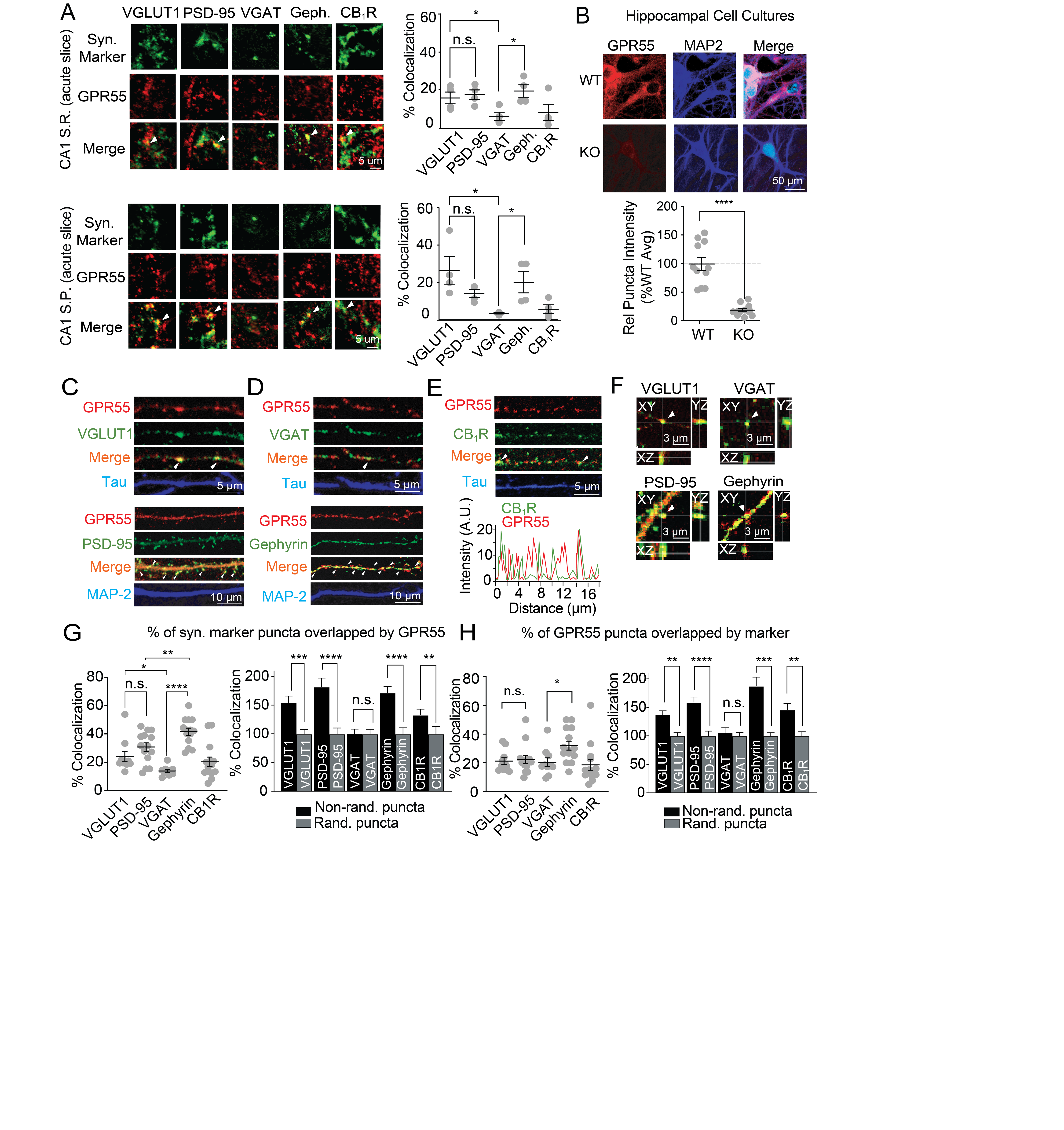

**Suppl. Fig. 3, Related to Fig. 2: GPR55 expression at excitatory and inhibitory synapses in *ex vivo* hippocampal slices and *in vitro* hippocampal cell cultures.**

(A) (Above): Representative confocal images (63X, mean projection from z-stacks) demonstrating GPR55 (red) colocalization with synaptic markers (green) in the CA1 *stratum radiatum* region of *ex vivo* hippocampal slices. Arrowheads depict putative colocalized puncta. GPR55 colocalized with both presynaptic excitatory terminals marked by VGLUT1 (16±3% of all VGLUT1+ terminals, n=4 slices) and excitatory postsynaptic densities labeled with PSD-95 (18±2%, n=4) (unpaired t-test VGLUT1 vs. PSD-95: p=0.68). However, GPR55 overlapped more with inhibitory postsynaptic densities identified by gephyrin (20±3%, n=4) than presynaptic inhibitory terminals marked by VGAT (6±2%, n=4) (p=0.014). GPR55 colocalized with VGLUT1+ terminals more than VGAT+ terminals (p=0.044), and overlapped with 8±4% of all CB_1_R terminals (n=4). (Below): GPR55 colocalization with synaptic markers in the CA1 *stratum pyramidale* region. While GPR55 colocalized similarly with VGLUT1 (44±12%, n=4) and PSD-95 (23±4%, n=4) puncta (p=0.15 VGLUT1 vs PSD-95), GPR55 displayed a larger degree of overlap with gephyrin puncta (34±9%, n=4) in comparison to VGAT puncta (6±1, n=4) (p=0.026 gephyrin vs VGAT). GPR55 also colocalized with more VGLUT1 puncta compared with VGAT puncta (p=0.020), and colocalized with 9±4% of all CB_1_R terminals (n=4). (B) GPR55 expression (red) is significantly reduced in neuronal cell cultures derived from GPR55 KO mice in comparison to WT controls (WT GPR55 intensity, relative to average 100±11%, n=11 vs KO 19±3%, n=12, unpaired t test: p<0.0001). (C-D) Representative confocal images (63X, mean projection from z-stacks) revealed GPR55 (red) colocalization (arrowheads) with synaptic markers (green) in hippocampal cell cultures. (E) Representative images and line scan of GPR55 (red) with CB_1_Rs (green) along a tau-labeled axon demonstrating alternating pattern along an axon with occasional overlap. (F) Higher resolution scans demonstrate GPR55 puncta colocalization with synaptic markers in three dimensions. (G) (Left): Quantification of % of total synaptic marker puncta that is colocalized with GPR55 puncta. While GPR55 colocalized with markers of both pre- and postsynaptic excitatory synaptic compartments (GPR55 with VGLUT1: 24±4% colocalization, n=10 coverslips; GPR55 with PSD-95: 31±3%, n=14; unpaired t test, p=0.16), GPR55 had greater colocalization with gephyrin+ inhibitory postsynaptic densities than presynaptic VGAT+ terminals (VGAT 14±1%, n=9; gephyrin 42±3%, n=14; unpaired t test, p<0.0001). Levels of GPR55 colocalization were higher in VGLUT+ terminals than VGAT+ terminals (p=0.022), and comparably greater in postsynapses labeled with gephyrin than PSD-95 (p=0.0092). GPR55 colocalized with 20±3% of CB_1_R puncta (n=14). (Right) In comparison to scrambled GPR55 pixels along a 2D axon/dendrite, GPR55 colocalized better than chance with VGLUT (non-randomized colocalization vs. randomized, as a % of randomized mean: 155±11%, paired t-test, p=0.0002), PSD-95 (182±15%, p<0.0001), gephyrin (171±11%, p<0.0001), and CB_1_R (133±10%, p=0.0061) but not VGAT (101±7%, p=0.93). See Materials and Methods for more details. (H) (Left): Quantification of % of total GPR55 puncta overlapped by synaptic marker puncta: VGLUT1 (21±2%, n=10), PSD-95 (22±3%, n=14), VGAT (20±3%, n=10), gephyrin (32±3%, n=14), and CB_1_R (19±4%, n=14). VGLUT1 colocalization with total GPR55 puncta was not significantly greater than PSD-95 colocalization with total GPR55 puncta (p=0.81). However, gephyrin colocalization with total GPR55 was greater than VGAT (p=0.017). (Right): Total GPR55 puncta colocalized with the following synaptic markers greater than scrambled GPR55 pixel controls: VGLUT1 (138±6%, p=0.0013), PSD-95 (159±9%, p<0.0001), gephyrin (188±16%, p=0.0001), and CB_1_R (146±11%, p=0.0069). However, VGAT did not colocalize with total GPR55 to a greater extent than GPR55 randomized pixel controls (p=0.55). Values represent colocalization as a % of mean scrambled pixel control.

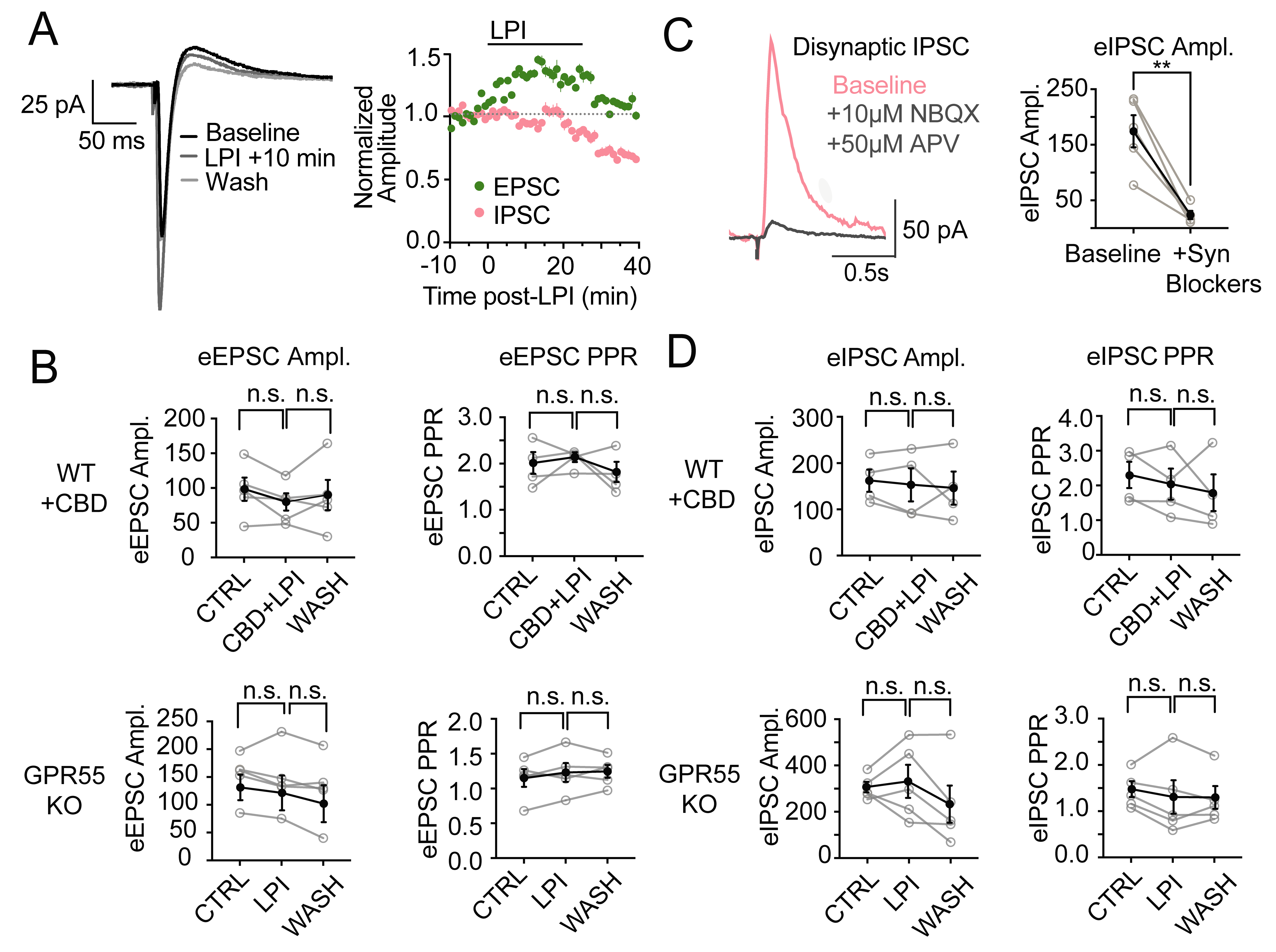

**Suppl. Fig. 4, Related to Fig. 3: Experimental controls for LPI-mediated changes in evoked synaptic transmission.** (A) In compound EPSC/IPSC recordings (n=9 pyramidal neurons held at -60 mV), LPI induces an early increase in SC-evoked EPSC amplitude (green) followed by a delayed reduction in IPSC amplitude (pink). 4 μM LPI was washed on for 30 min before a 10 min ACSF washout. (B) In the presence of 1 μM CBD, 4 μM LPI did not alter evoked (e)EPSCs (n=5, p=0.095 vs baseline) or the paired pulse ratio (PPR) between the second and first eEPSC (p=0.59). Additionally, LPI had no effect on eEPSC amplitude (p=0.51) or PPR (p=0.26) in GPR55 KO mice. (C) Disynaptic IPSCs were significantly reduced by the glutamatergic synaptic blockers NBQX (10 μM) and APV (50 μM) (n=5, p=0.0051). (D) Addition of 1 μM CBD prevented LPI-induced changes in eIPSC amplitude (n=4, p=0.54) and PPR (n=4, p=0.37), as did genetic deletion of GPR55 (n=5, Ampl: p=0.67, PPR: p=0.44).

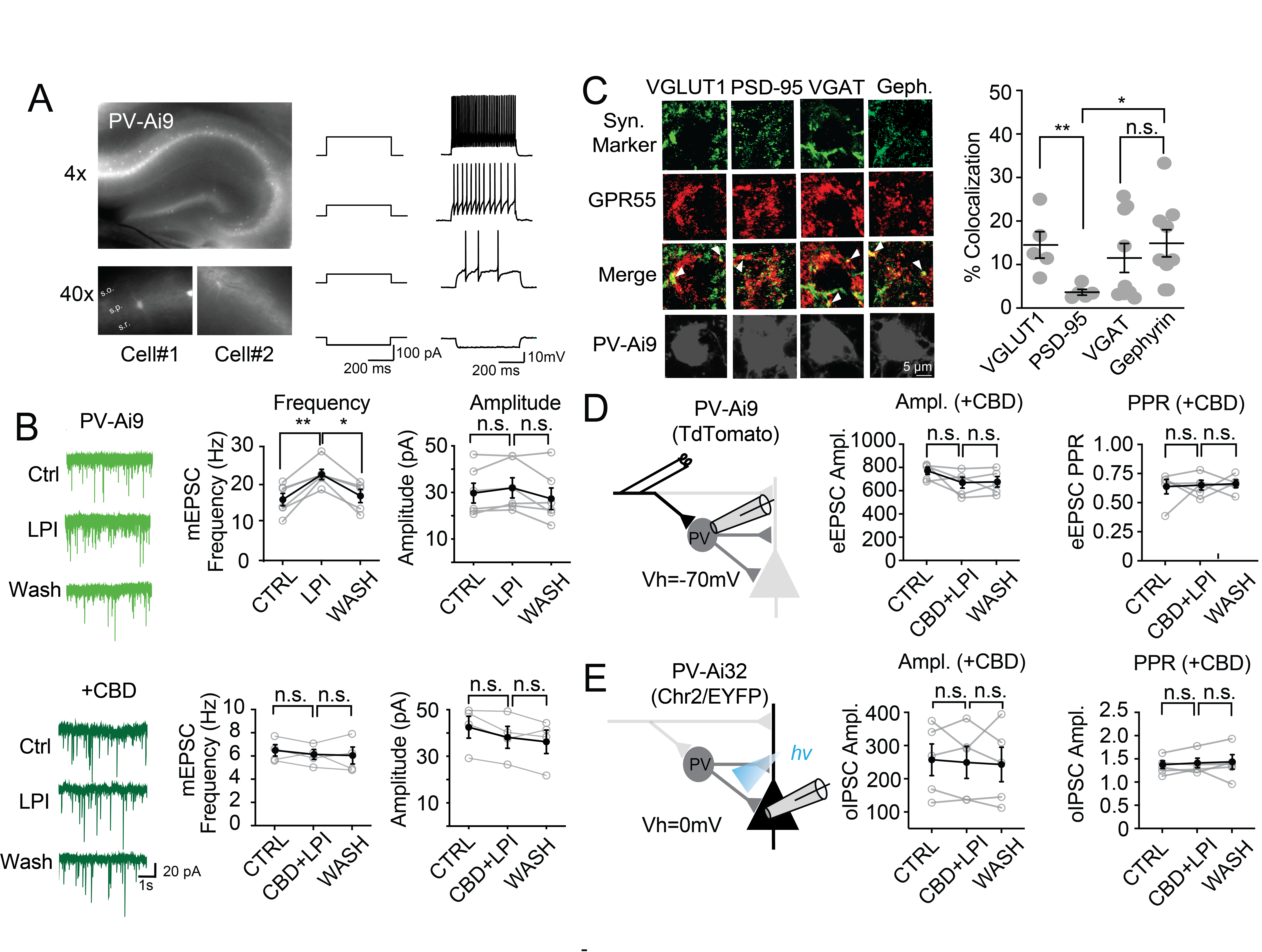

**Suppl. Fig. 5, Related to Fig. 4: CBD blocks LPI effects at both E→I and I→E synapses.** (A) Labeled PV neurons in slices from PV-Cre X Ai9/TdTomato mice at 4x and 40x magnification. Whole cell recordings from genetically labeled PV neurons demonstrated characteristic fast spiking behavior. (B) LPI (4 μM) transiently elevated mEPSC frequency in PV interneurons (n=6, p=0.0033 CTRL vs. LPI, p=0.019 LPI vs. WASH), blocked by pre-treatment with 1 μM CBD (p=0.35 CBD+LPI vs. baseline). (C) Representative images (left) and quantification (right) of GPR55 (red) colocalization (arrowheads) with synaptic markers (green) in PV-Ai9 genetically labeled neurons (gray). Quantification of the % of individual synaptic marker puncta that are colocalized with GPR55 puncta: GPR55 puncta colocalized more with VGLUT1+ (15±3%, n=5) vs PSD-95 (4±1%, n=5) (unpaired t-test, p=0.0083). However, GPR55 colocalized to a similar level with VGAT (12±3%, n=9) and gephyrin (15±3%, n=9) (p=0.47). GPR55 overlap with gephyrin was greater than that with PSD-95 (p=0.023). (D) Recording in PV+ interneurons, CBD impeded LPI-mediated elevations in eEPSCs (see Fig. 4; CBD+LPI vs. baseline p=0.74) and reductions in PPR (p=0.50). mEPSC and eEPSCs both acquired during a 5 min window, 5 min post LPI application. (E) CBD prevented the LPI-mediated reduction of optogenetically evoked PV monosynaptic IPSCs (see Fig. 4: PV-Cre X Ai32/ChR2 mice, n=5, CBD+LPI p=0.14 vs. baseline), without changing the PPR (p=0.88). Analysis was performed in a 5 min window, 25 min post LPI application.

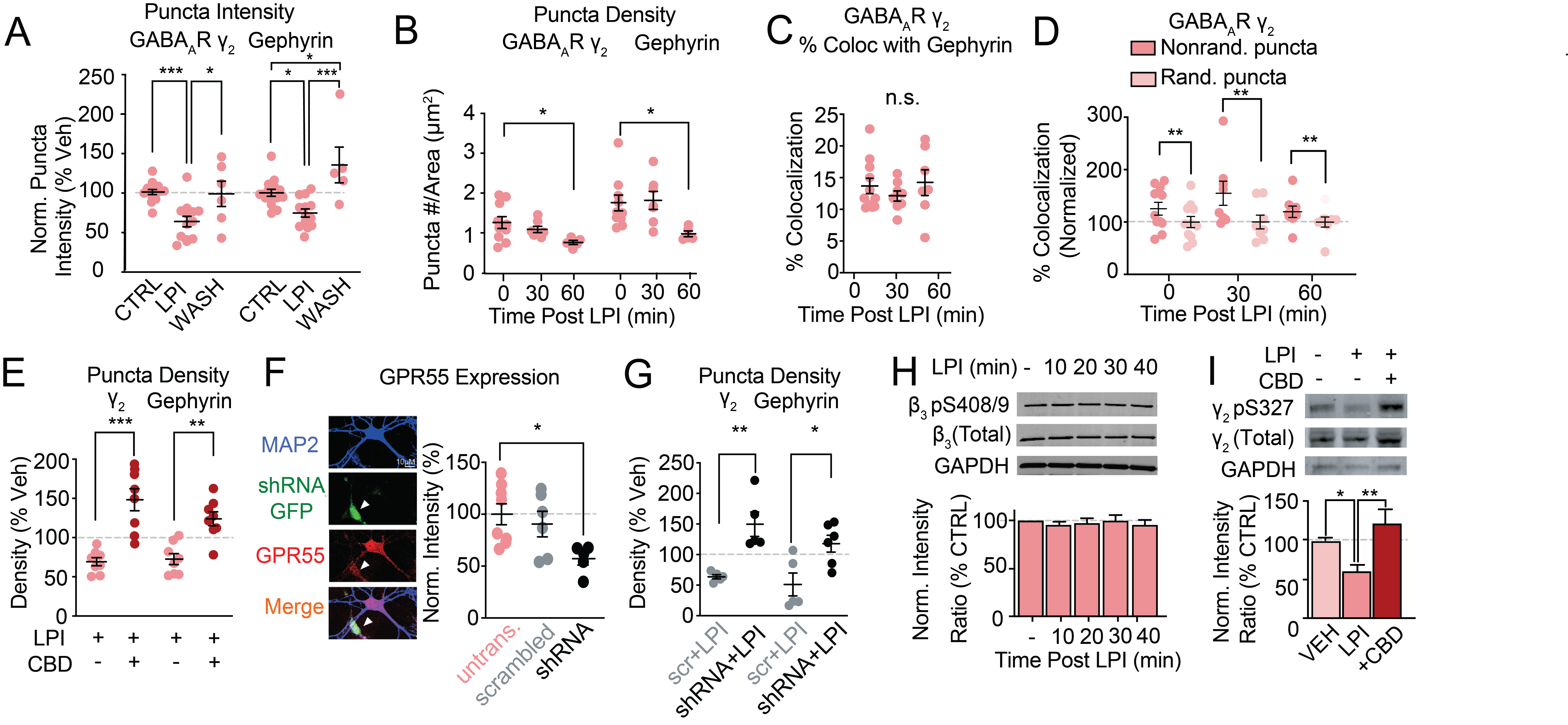

**Suppl. Fig. 6, Related to Fig. 5: LPI reversibly decreases GABA_A_R γ_2_ and gephyrin puncta intensity and density in a GPR55-dependent manner.** (A) LPI-mediated reduction in GABA_A_R γ_2_ puncta intensity reverses following 10 min wash with ACSF (intensity as % of average control: CTRL 100±3%, n=14; LPI 62±7%, n=13; WASH 98±16%, n=5; one-way ANOVA p=0.0006: VEH vs LPI p=0.0008, LPI vs WASH p=0.014, CTRL vs WASH p=0.98). Following 10 min of ACSF wash, intensity of gephyrin puncta increased to levels greater than pre-LPI administration (CTRL 100±4%, n=15; LPI 74±5%, n=13; WASH 135±23%, n=5; one-way ANOVA p=0.0002: VEH vs LPI p=0.028, LPI vs WASH p=0.0002, VEH vs WASH p=0.027). (B) LPI decreased the density of GABA_A_R γ_2_ puncta per unit area at 60 min but not at 30 min post-application (VEH 1.3±0.1/µm^2^, n=10; LPI-30 1.1±0.1/µm^2^, n=7; LPI-60 0.8±0.1/µm^2^, n=7; one-way ANOVA: p=0.048, VEH vs LPI-30 p=0.51, VEH vs LPI-60 p=0.029). Similarly, LPI reduced the density of gephyrin puncta at only 60 min post-treatment (VEH 1.7±2/µm^2^, n=10; LPI-30 1.8±0.2/µm^2^, n=7; LPI-60 1.0±0.1/µm^2^, n=7; one-way ANOVA: p=0.038, VEH vs LPI-30 p=0.92; VEH vs LPI-60 p=0.042). (C) No significant change in GABAR γ_2_ and gephyrin puncta colocalization following LPI treatment (% of gephyrin puncta overlapped by GABA_A_R γ_2_ following LPI treatment: 0 min (13.7±1%, n=10), 30 min (12±1%, n=7), 60 min (14±2%, n=7); 1-way ANOVA: p=0.57). (D) At all points following LPI treatment, GABAR γ_2_ colocalization with gephyrin puncta was greater than GABAR γ_2_ scrambled pixel controls: 0 min (p=0.0011), 30 min (p=0.0070), 60 min (p=0.0013). Values represent colocalization as a % of mean scrambled pixel control. (E) 1 μM CBD blocked the LPI-driven decrease in GABA_A_R γ_2_ puncta (LPI 69±5% relative to vehicle controls, n=7 vs CBD+LPI 148±14%, n=8; p=0.0001, unpaired t-test) and gephyrin puncta per unit area (LPI 73±7%, n=7 vs CBD+LPI 124±9%, n=8; p=0.0005, unpaired t-test), (F) Lentiviral-mediated shRNA targeting GPR55 decreased GPR55 expression to 57±6% of untransfected levels (n=5, p=0.034), effects not seen with scrambled shRNA controls (91±12%, n=6, p=0.89 vs untransfected). (G) Compared to scrambled shRNA controls (“scr”), in cultures transfected with GPR55 shRNA, LPI failed to reduce GABA_A_R γ_2_ density (scr+LPI 64±3% relative to vehicle controls, n=5 vs shRNA+LPI 150±20%, n=5; p=0.0028). Similarly, shRNA knockdown of GPR55 blocked LPI-mediated decreases in gephyrin density relative to scrambled shRNA (scr+LPI 51±19%, n=5 vs shRNA+LPI 118±14%, n=5; p=0.017). (H) LPI did not change β_3_ S408/409 phosphorylation at any time post-LPI treatment (one-way ANOVA p=0.83). (I) 1 μM CBD prevented LPI-mediated decrease in GABA_A_R γ_2_ intensity (VEH 100±3%, n=4; LPI 60±8%, n=11; LPI+CBD 122±20%, n=4; one-way ANOVA p=0.0028: VEH vs LPI p=0.047, LPI vs LPI+CBD p=0.0027, VEH vs CBD+LPI p=0.50).

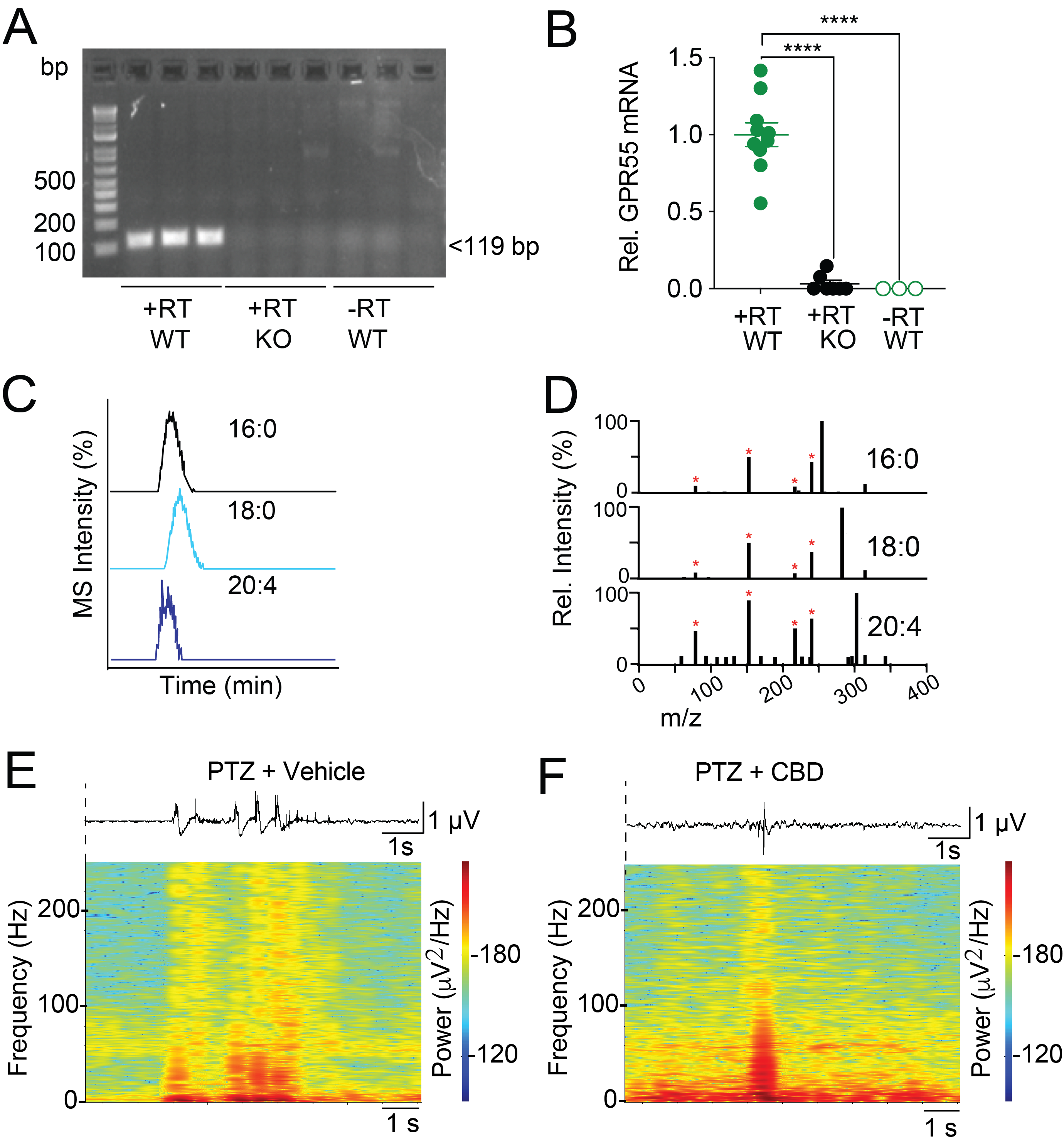

**Suppl. Fig. 7, Related to Fig. 6: Controls for qPCR measurements of GPR55 and HPLC-MS assay for LPI.** (A) DNA gel indicating the major PCR product (119 bp) from WT hippocampal lysates prepared with reverse transcriptase (+RT/WT), absent in both age-matched KO controls with RT (+RT/KO), and WT controls without RT (-RT/WT). (B) Quantification of qPCR results, indicating that detection of GPR55 mRNA is significantly reduced in hippocampi from GPR55 KO mice, and in the absence of RT (+RT/WT 1.00±0.08, n=10; +RT/KO 0.03±0.02, n=7; -RT/WT 0±0, n=3; one-way ANOVA p<0.0001: +RT/WT vs +RT/KO p<0.0001, +RT/WT vs. -RT/WT p<0.0001). All PCR products were confirmed by Sanger sequencing. (C-D) Chromatographic separation of LPI isoforms (16:0 black, 18:0 light blue, 20:4 purple) by liquid chromatography (C) and structural confirmation by MS/MS fragmentation (D), lowest concentration standard shown for each. (E-F) Representative EEG traces and power spectra from matched mice treated with PTZ (60 mg/kg) and either vehicle (E) or CBD (200 mg/kg) (F), 1 hour prior to seizure induction.

|  | **PTZ-induced Seizures (mice)** | | | **Lithium-Pilocarpine Epileptogenesis (rat)** | | |
| --- | --- | --- | --- | --- | --- | --- |
|  | **16:0 LPI** | **18:0 LPI** | **20:4 LPI** | **16:0 LPI** | **18:0 LPI** | **20:4 LPI** |
|  | **nmol/g tissue** | | | **nmol/g tissue** | | |
| **Non Seizure - 1** | 43.23325667 | 59.8540892 | 30.96121 | 0.10089827 | 1.49556247 | 0.16854048 |
| **Non Seizure - 2** | 112.760734 | 104.148024 | 32.08553 | 0.08490841 | 0.99412524 | 0.12995733 |
| **Non Seizure - 3** | 46.65924831 | 60.6071709 | 50.71104 | 0.03453195 | 0.4143992 | 0.55865199 |
| **Non Seizure - 4** | 126.7829041 | 119.655144 | 43.68935 | 0.03787319 | 0.9527208 | 0.04326028 |
| **Non Seizure - 5** | 62.26182941 | 14.8855624 | 47.9851 | 0.02781658 | 0.56509434 | 0.06312231 |
| **Non Seizure - 6** | 41.1331132 | 42.0983307 | 31.15213 | 0.01313484 | 0.51794224 | 0.23814269 |
| **Non Seizure - 7** | 63.81042006 | 120.026382 | 29.41262 | - | - | - |
| **Non Seizure - 8** | 48.12298467 | 60.1828995 | 39.89212 | - | - | - |
| **PTZ + Veh - 1** | 107.5209821 | 170.228296 | 30.81271 | 0.09693066 | 1.26923239 | 0.19181266 |
| **PTZ + Veh - 2** | 109.0377524 | 209.441582 | 42.66049 | 0.02337486 | 0.46157969 | 0.20723709 |
| **PTZ + Veh - 3** | 36.65704982 | 59.5783127 | 30.1657 | 0.00342776 | 0.28385636 | 0.10330653 |
| **PTZ + Veh - 4** | 63.81042006 | 113.503209 | 33.75079 | 0.10062834 | 0.55089549 | 0.28897336 |
| **PTZ + Veh - 5** | 58.76159028 | 144.326527 | 35.08725 | 0.05968903 | 0.62179749 | 0.0954903 |
| **PTZ + Veh - 6** | - | - | - | 0.02384377 | 0.23414265 | 0.32523686 |
| **PTZ + CBD - 1** | ∅ | ∅ | ∅ | 0.02583883 | 0.65064933 | 0.07354184 |
| **PTZ + CBD - 2** | 146.8085146 | 49.1094157 | ∅ | 0.04864788 | 0.49449746 | 0.19589096 |
| **PTZ + CBD - 3** | 40.47549251 | 40.5921671 | 31.50215 | 0.06702039 | 0.53334739 | 0.24358883 |
| **PTZ + CBD - 4** | 55.78108362 | 52.5884413 | 28.98834 | 0.04702657 | 0.54318937 | 0.07330862 |
| **PTZ + CBD - 5** | 41.7058796 | 35.0978524 | 30.12327 | 0.01008256 | 0.42333291 | 0.01474997 |
| **PTZ + CBD - 6** | - | - | - | 0.03059547 | 0.21165581 | 0.0910401 |

**Supplemental Table 1, related to Fig. 6, 7: LPI concentrations (nmol/g tissue) PTZ- and Li-PLC induced seizures.** “∅” indicates undetectable, “-“ indicates no sample

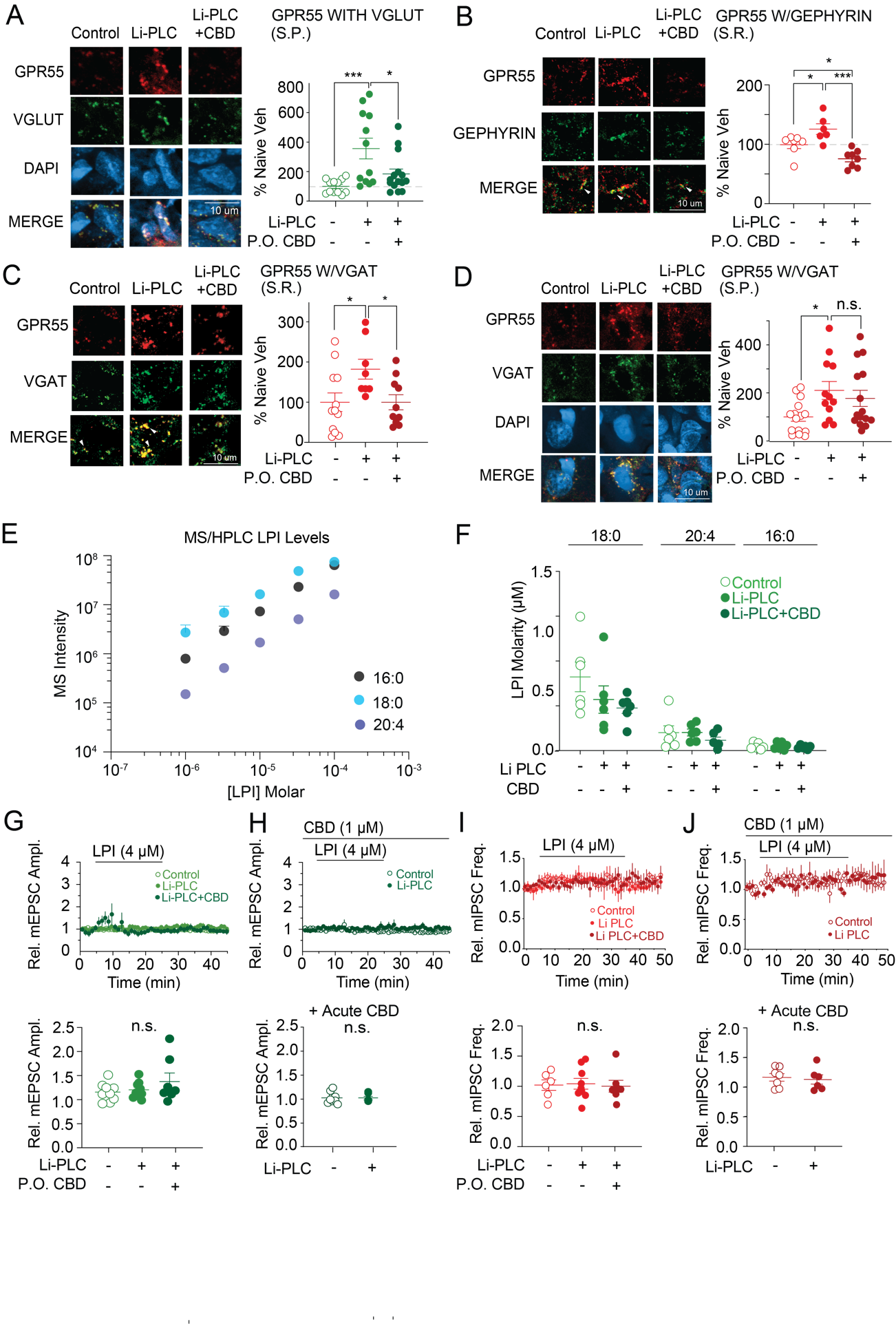

**Suppl. Fig. 8, Related to Fig. 7: Li-PLC epileptogenesis elevates GPR55 expression in both excitatory and inhibitory synapses*,* and electrophysiological negative controls.** (A) CBD (200 mg/kg, p.o.) prevents the Li-PLC-induced rise in the intensity of GPR55 puncta colocalized with VGLUT1 in the hippocampal *stratum pyramidale* (S.P.) region (control 100±14% as % of average, n=12; Li-PLC 357±70%, n=12; Li-PLC+CBD 184±34%, n=15; one-way ANOVA p=0.0012: control vs Li-PLC p=0.0007, Li-PLC vs Li-PLC+CBD p=0.017). (B) Li-PLC epileptogenesis increases GPR55 puncta intensity at inhibitory synapses labeled by gephyrin in the *stratum radiatum*, prevented by chronic CBD treatment (control 100±7%, n=7; Li-PLC 126±9%, n=6; Li-PLC+CBD 76±5%, n=15; one-way ANOVA p=0.0003: control vs Li-PLC p=0.041, Li-PLC vs Li-PLC+CBD p=0.0002, control vs Li-PLC+CBD p=0.047). (C-D) Intensity of GPR55 puncta colocalized with VGAT, a presynaptic marker of inhibitory synapses, was nearly doubled following Li-PLC administration (one-way ANOVA p=0.032, control 100±23%, n=12 vs Li-PLC 182±25%, n=8; p=0.035), effects prevented by CBD p.o. in S.R. (C, Li-PLC+CBD 100±19%, n=10; p=0.043) but not in S.P. (D, p=0.69). (E) Linear relationship in standard control curves for 3 LPI isoforms (16:0 black, 18:0 light blue, 20:4 purple) across increasing concentrations of LPI. (F) Levels of LPI isoforms (18:0, 20:4, and 16:0) remained unchanged following Li-PLC epileptogenesis (ANOVA comparing control, Li-PLC, Li-PLC+CBD changes in each isoform: 18:0 p=0.20; 20:4 p=0.45, 16:0 p=0.76), (G-H) Acute wash-on of 4 μM LPI alone (G) or 1 μM CBD + 4 μM LPI (H) does not change mEPSC amplitude in *ex vivo* hippocampal slices non-epileptogenic controls (unfilled green circles), Li-PLC-treated animals (filled light green circles), or Li-PLC-treated animals with chronic CBD p.o. (filled dark green circles) (one-way ANOVA p=0.26). (I-J) Acute wash-on of 4 μM LPI alone (I) or 1 μM CBD + 4 μM LPI (J) does not change mIPSC frequency in *ex vivo* hippocampal slices from either control animals (unfilled red circles) and Li-PLC-treated rats (filled light red circles), or Li-PLC-treated rats with chronic CBD p.o. (filled dark red circles) (one-way ANOVA p=0.95).

***Supplemental Methods***

**Animals**

All procedures involving animals were approved by the Institutional Animal Care and Use Committee at the New York University Langone Medical Center, and in accordance with guidelines from the US National Institutes of Health. GPR55 KO (*B6;129S-Gpr55^tm1Lex^/Mmnc*) mice were generously provided by Prof. Ken Mackie, Indiana University, and backcrossed to a C57Bl/6J (WT, +/+) background at NYU to expand the colony. For cell culture experiments, Sprague-Dawley rat pups were utilized.

PV interneurons were identified with parvalbumin (PV)-Cre (*B6;129P2-Pvalb^tm1(cre)Arbr^/J*) line crossed with a Ai9*/*TdTomato expressing lines (*B6.Cg-Gt(ROSA)26Sor^tm9(CAG-tdTomato)Hze^/J*) (Jackson Labs). For optogenetic experiments involving PV+ -mediated inhibition, PV-Cre mice were crossed with an Ai32 line expression channelrhodopsin 2 EYFP fusion protein *(B6;Cg-Gt(ROSA)26Sor^tm32(CAG-COP4*H134R/EYFP)Hze^/J*) (Jackson Labs).

Male Wistar-Kyoto rats (>P21) were used for studies involving lithium pilocarpine-induced status epilepticus at the University of Aston (UK). All rats were maintained in 12h:12h dark:light cycle, a room temperature of 21°C and humidity of 50±10 %, with *ad libitum* access to food and water. The experiments were performed in accordance with UK Home Office regulations (Animals Scientific Procedures Act, 1986). Male Swiss mice weighing 18-25g (Taconic, Germantown, NY) were used for electrographic pentylenetetrazole (PTZ)-induced seizures (Fig. 7) as previously described (Vilela et al., 2017).

**Induction of pentylenetetrazole (PTZ)-induced seizures in mice (Fig. 1)**

During a pre-experimental baselining period, animals were randomized for treatment and treatment groups balanced within litters to the extent permitted by individual litter size, sex, and genotype distribution. Male GPR55^+/+^ (n=40) and GPR55^-/-^ (n=40) mice, aged P24-28 and weighting 25-40g, were injected intraperitoneally (i.p.) with vehicle (ethanol: kolliphor^®^: 0.9% saline=1:1:18; Sigma-Aldrich, UK) 1h before receiving i.p. administration of PTZ (Sigma-Aldrich, UK) at 85, 95, 105 or 115 mg/kg (n=10 per group) dissolved in saline (0.9% (w/v) NaCl). During the 30 min following PTZ injection, the number of animals exhibiting fully developed tonic-clonic seizures with loss of righting reflex and/or death was calculated.

The acute experimental phase of the study utilized a similar design, but included administration of plant-derived, highly purified cannabidiol (CBD, generously provided by GW Research Ltd, Cambridge, UK) at doses of 50, 100, or 200 mg/kg, or vehicle, given 30 min before injection of PTZ (105 mg/kg, i.p) or vehicle. Animals were sacrificed within 30 min of PTZ injection or if premature death occurred following status epilepticus.

**EEG recordings and analysis for high dose PTZ Seizures (Fig. 1, Suppl. Fig. 1)**

**Implants:** Mice (n=3 WT, n=3 GPR55 KO, 3-4 months old) were anesthetized with 1.5-2% isoflurane (2 L/min) and provided with a local anesthetic to the incision site (bupivicaine at .05 mg/kg, 2.5 mg/ml, s.c.). The skull was cleaned with saline and hydrogen peroxide and ground wires (bare stainless steel) were positioned intracranially over the cerebellum. The skull was then coated with Optibond (Kerr Dental, Brea, CA) and a craniotomy (~0.25 x 0.25 mm) was performed at AP -2.2, ML -2.0 (left hemisphere). The dura was pierced, and the tungsten wires were implanted ~0.9mm into the brain. The wires with custom driver were cemented to the skull with C & B Metabond Quick Adhesive Cement (Parkell) and Unifast Trad acrylic (GC America). The craniotomy was capped with a mixture of mineral oil (one part) and dental wax (three parts), and a Faraday cage was constructed using copper mesh and connected to the cerebellar ground wire. Following surgery, an opioid analgesic was injected (Buprenex at 0.06 mg/kg, 0.015 mg/ml, i.m.) and given as needed for the next 1-3 days.

**Recording:** Three insulated tungsten wires (0.002” California Fine Wire, CA) were glued together and cut at a 45⁰ angle. These were then mounted to a screw and soldered to a header pin (Mill-Max, NY). Signals were digitized at 30 KHz with the Intan amplifier board (RHD2132/RHD2000 Evaluation System, Intan). Mice were allowed at least a one week post-implantation recovery before electrodes were lowered into place. CA1 ripples (150 – 200Hz oscillation) were used to identify correct placement prior to experimentation. At least one day was allowed between electrode placements and seizure induction. On the day of seizure induction, mice were placed in their home cage, and the position was monitored by a USB webcam synchronized with the neural recordings through a blinking LED. First, CBD (200 mg/kg i.p.) was injected, followed 1 h later by injection of PTZ (105 mg/kg i.p.). Recordings were made for at least 30 minutes or until death.

**Analysis:** Behaviorally observed seizure state was classified independently based on neural activity and on behavior using the Racine scale (Racine, 1972). Electrographic events were characterized in 30 second time blocks as one of four classifications: **(***1) Normal:* normal EEG background without epileptiform discharges, seizures, or slowing; *(2) Epileptiform discharges* (Gelinas et al., 2016) clusters of >2 spikes, with absolute value of LFP amplitude larger than 4 standard deviations of the LFP recorded throughout, spike duration less than 75 ms in duration as measured by the width at half max, and no evolution over the duration of 30 s time window; **(***3) Seizures* (Van Erum et al., 2019) high amplitude, rhythmic trains of high frequency complex discharges lasting for a minimum of 5 s; (4) *Post-ictal suppression:* LFP values in the bottom decile of LFP values recorded throughout the experiment. Spectrograms were made in MATLAB v2018b (Mathworks, MA) using Morlet wavelets on the down-sampled (1250 Hz) local field potential from the recording site with the largest amplitude ripples.

**Induction of lower dose pentylenetetrazole (PTZ)-induced seizures in mice (Fig. 7)**

Methods were performed as previously described (Vilela et al., 2017). In brief, male Swiss Mice weighing 18-25 g (Taconic, Germantown, NY) were used. Four total electrodes were placed over left frontal, right occipital and bilateral hippocampi. Animals were treated subcutaneously (s.c.) with either vehicle (ethanol: kolliphor^®^: 0.9% saline=1:1:18; Sigma-Aldrich, UK) or 200 mg/kg CBD (plant-derived highly purified, generously provided by GW Research Ltd, Cambridge, UK). One hour later, animals were injected with pentylenetetrazol (60 mg/kg s.c.; Sigma-Aldrich, UK). During seizure induction, mice were monitored using a webcam synchronized with the EEG recordings. Seizure behavior was analyzed by a blinded observer using Racine scale as described (Racine, 1972). Electrographic activity was analyzed with Sirenia (v2.2.1; Pinnacle) or Spike 2 (v7.12; Cambridge Electronics). Recordings were analyzed for latency to the first abnormal event (epileptiform activity or seizure), seizure duration, and power. Power Spectrograms were made in MATLAB (v2018b; Mathworks, MA) using Morlet wavelets of the down-sampled (1250 Hz) EEG from the recording site.

**Induction of chronic epilepsy in Wistar-Kyoto rats**

Using a recently described model of low-mortality, high-morbidity status epilepticus (Modebadze et al., 2016), male (P>21) Wistar-Kyoto rats (Charles River) were injected subcutaneously (s.c.) with lithium-chloride (127 mg/kg) 24 hours before pilocarpine treatment, to sensitize them to the seizurogenic effects of pilocarpine. 30 minutes prior to pilocarpine treatment, animals were injected with methylscopolamine (Sigma-Aldrich, UK; 1 mg/kg; s.c.), a peripherally restricted muscarinic receptor antagonist that minimizes the peripheral effects of pilocarpine (salivation etc.). Pilocarpine (Sigma-Aldrich, UK; 25 mg/kg; s.c.) was administered at 30-min intervals until bilateral rearing and falling (Racine stage 3.5) was observed. Xylazine (Sigma-Aldrich, UK; 2.5 mg/kg; s.c.) was then injected to inhibit motor movement during seizures. Following 60 minutes of sustained seizure activity, animals were injected with “STOP” solution (1 ml/kg; s.c.), consisting of diazepam (Sigma-Aldrich, UK; 2.5 mg/kg), 2-methyl-6-(phenylethynyl) pyridine (Sigma-Aldrich, UK; 20 mg/kg), and dizoclipine (MK-801; Sigma-Aldrich, UK; 0.1 mg/kg) to prevent further seizures. During the 2 weeks after induction, regular animal welfare checks were performed. Epileptogenesis was determined using a previously validated post-seizure behavioral battery (PSBB) test, and only animals with PSBB scores >10 following a 10 week period were used for electrophysiology and immunohistochemistry (Modebadze et al., 2016). In the weeks following the establishment of epileptogenesis, rats were randomly divided into two groups to receive either 200 mg/kg CBD or vehicle (3.5% kolliphor ® HS, Sigma) as per (Patra et al., 2019). All drugs were administered in drinking water to reduce the incidence of spontaneous seizures with epileptic animals during frequent handling.

**Two-Pulse Kainic Acid (KA) Seizure Induction**

For the “KA1” pulse, WT C57Bl6 mice (2-3 months) received either CBD (200 mg/kg) or CBD vehicle (ethanol: kolliphor^®^: 0.9% saline=1:1:18; Sigma-Aldrich), followed by kainic acid (KA, Sigma, 24 mg/kg in normal saline, s.c.) 1 h later. Behavioral seizures were then scored based on a modified Racine scale (Racine, 1972): 0 – Grooming, moving, normal; 1 – Freezing, splayed limbs, sloppy movements; 2 – Ear twitching, head nodding; 3 - Unilateral arm jerks; 4 - Bilateral arm jerks, with stages 3-4 identifying convulsive seizures. Animals were then given diazepam (Hospira, Inc, 10 mg/kg, s.c.) 2 hours later to prevent status epilepticus. For qPCR experiments, 48 h following KA1 injection, animals were sacrificed and brains were rapidly frozen for mRNA extraction. For those receiving the “KA2” pulse, 48 h following KA1, a second dose of CBD or vehicle (concentrations as above), followed by KA (as above) was given. Behavioral seizures were again assayed as per the modified Racine Scale.

**Electrophysiology slice preparation (for mice, all except Fig. 7)**

GPR55 KO or littermate control mice 2-3 months old were anesthetized with a mixture of ketamine/xylazine (150 mg/kg and 10 mg/kg, respectively) and perfused transcardially with an ice-cold sucrose solution containing (in mM): 206 Sucrose, 11 D-Glucose, 2.5 KCl, 1 NaH_2_PO_4_, 10 MgCl_2_, 2 CaCl_2_ and 26 NaHCO_3_. Following perfusion and decapitation, brains were removed and placed in the cold sucrose for sectioning before gluing to the stage of a Leica VT 1000S Vibratome. Transverse, 350 μm sections of left and right hippocampi were cut and transferred to an oxygenated, 34°C recovery chamber filled with artificial cerebrospinal fluid (ACSF) containing (in mM): 122 NaCl, 3 KCl, 10 D-Glucose, 1.25 NaH_2_PO_4_, 2 CaCl_2_, 1.3 MgCl_2_, and 26 NaHCO_3_. Slices were allowed to recover for 1 h at 34°C and were then maintained at room temperature in oxygenated ACSF for 1-6 h before recording.

**Electrophysiological recordings (for mice, all except Fig. 7)**

Hippocampal slice recordings were performed in a submerged chamber maintained at 32-34°C with a constant bath perfusion of ACSF (with or without pharmacological agents) at ~4 mL/min. Slices equilibrated in the chamber for >10 minutes before recording. Whole cell and cell-attached recordings were made with borosilicate glass pipettes pulled on a Sutter Instrument P-97 micropipette puller. Tip resistance ranged between 2-5 MΩ following fire polishing to enhance seal quality.

For voltage clamp recordings of spontaneous and evoked EPSCs, the intracellular solution contained (in mM): 130 CsMeSO3, 6 CsCl, 1 MgCl2, 10 HEPES, 0.3 EGTA, 10 Tris-Phosphocreatine, 4 Mg-ATP and 0.3 Na-GTP. Spontaneous IPSCs onto pyramidal cells were recorded in voltage clamp using a high Cl- internal solution containing (in mM): 70 CsMeSO_3_, 35 CsCl, 15 TEA-Cl, 1 MgCl_2_, 0.2 CaCl_2_, 10 HEPES, 0.3 EGTA, 10 Tris-Phosphocreatine, 4 Mg-ATP and 0.3 Na-GTP, with ACSF glutamatergic blockers 10μM NBQX and 50 μM APV. For mEPSC and mIPSC recordings, 1μM TTX was added to ACSF. For disynaptic IPSC experiments, cells were held at 0mV and stabilized for at least 10-15 minutes prior to recording. For synaptic stimulation recordings, stimulating electrodes were placed in the *stratum radiatum*, and a 10x 10Hz frequency train was delivered, with either 15s (eEPSCs) or 30s (eIPSCs) between trains.

Whole-cell patch clamp recordings were performed on neurons in the CA1 region of the hippocampus, identified visually with an upright microscope (Zeiss Axioskop 2 FS Plus) using infrared differential interference contrast (IR-DIC) optics. For all experiments involving lysophosphatidylinositol (LPI) application, cells were recorded in regular ACSF for 10-15 minutes, followed by LPI (4 μM, Sigma in DMSO vehicle) for 30 minutes, and a subsequent 10-15 minute washout period. In a subset of experiments, plant-derived highly purified CBD (GW Research Ltd, Cambridge, UK, 1μM in DMSO vehicle) was added for 20 minutes prior to concomitant LPI treatment (CBD+LPI).

Data were recorded with a MultiClamp 700B amplifier (Axon Instruments), filtered at 10 kHz using a Bessel filter and digitized at 20 kHz with a Digidata 1322A analogue-digital interface (Axon Instruments). mEPSCs and mIPSCs were analyzed using Clampfit software, and evoked EPSCs/IPSCs were identified offline using a custom analysis script developed in MATLAB (Mathworks). Passive properties were continuously monitored during recordings, and all cells with V_m_ <-55 mV, Series Resistance (R_s_) > 25MΩ, or significant changes in R_s_ during the course of the recording, were excluded from analysis.

**Electrophysiology slice preparation (for rats, Fig. 7 only)**

Healthy and epileptic rats (p>66) were decapitated under deep isoflurane anaesthesia (4%^w^/_v_ in O_2_), and their brains were rapidly removed and placed in ice-cold artificial cerebrospinal fluid (ACSF) solution containing (in mM): 130 NaCl, 24 NaHCO_3_, 3.5 KCl, 1.25 NaH_2_PO_4_, 2.5 CaCl_2_, 1.5 MgSO_4_, 10 glucose saturated with 95% O_2_, 5% CO_2_, at pH 7.3. Transverse slices (350 μm thickness) including hippocampus were cut using a Vibroslice (Campden instruments Ltd., Loughborough, Leicestershire, UK) and transferred to a nylon mesh where they were maintained submerged in a chamber containing ACSF at 33˚C for 30 min. Slices were then maintained at room temperature (18-22˚C) in ACSF (95% O_2_ / 5% CO_2_).

Acute slices were secured under a nylon mesh, submerged, and superfused with ACSF in a chamber mounted on the stage of an upright microscope (Scientifica, UK). Slices were visualized with a 40×/0.1 NA water-immersion objective coupled with infrared and differential interference contrast (DIC) optics linked to a video camera (digital camera Orca 03G, Hamamatsu, Hamamatsu City, Japan).

**Electrophysiological recordings (for rats, Fig. 7 only)**

Somatic whole-cell patch-clamp recordings (at ~33°C) were made from visually identified cells using borosilicate glass capillaries (GC150F-10; Harvard Apparatus Ltd., Kent, UK) using a P1000 Flaming Brown Micropipette puller (Sutter Instruments Co., California, USA) and filled with a filtered intracellular solution consisting of (in mM): 120 K-gluconate, 4 KCl, 4 Mg-ATP, 10 HEPES, 0.3 Na_2_-GTP, 10 Na_2_-phosphocreatine, pH adjusted to 7.2 with KOH. Resistance of the patch pipettes was 5-6 MΩ. Recordings were accepted only if the initial seal resistance was >1 GΩ and series resistance did not change by more than 20% throughout the recording period. No correction was made for the junction potential between the pipette and the ACSF. Pyramidal neurons were distinguished from interneurons by the localization of their somata in CA1 *stratum pyramidale*, a lower input resistance, a higher membrane constant, a smaller fast after-hyperpolarization and adapting, <20 Hz maximum firing rates. Intracellular signals were digitised to a computer with an A–D converter (Digidata 1500, Molecular Devices) and monitored during experiments with Clampex software (Molecular Devices). All electrophysiological signals were amplified (Multiclamp 700B, Molecular devices), low pass filtered at 10 kHz, digitized at 20 kHz.

In recordings of miniature excitatory postsynaptic currents (mEPSCs; voltage-clamp), the AMPA receptor component was isolated by adding 0.1 μM CGP 55845 (Abcam, UK), 100 μM D-APV (Abcam, UK), 500 μM MCPG (Abcam, UK), 1 μM strychnine (Sigma Aldrich, UK), 20 μM Bicuculline (Abcam, UK), 500 nM AM281 (Sigma Aldrich, UK) and 1 μM TTX (Tocris, UK) in the ACSF. GPR55 receptors were activated by applying 4 μM LPI (Sigma Aldrich, UK). In some experiments plant-derived highly purified CBD (GW Research Ltd, Cambridge, UK) was added to the ACSF at a final concentration of 1 μM. The DMSO concentration never exceeded 0.01%^w^/_v_.

**Two-Photon Calcium Imaging**

To image GCaMP6f confined to PV+ interneuron terminals (Suppl. Fig. 2E) we used acute hippocampal slices derived from PV-Cre x Ai148 mice. Images were focused at the *stratum pyramidale* and *stratum radiatum* in the CA1 hippocampus. The field of view was imaged at 30 Hz by using a resonant scanner-based two-photon microscope (Sutter), and individual frames were subsequently averaged before analysis. 4 μM LPI was washed onto slices for ~3 min, followed by 25 mM KCl for ~1 min as a positive control to ensure the integrity of axon terminals to flux Ca^2+^. Calcium responses were measured as a ratio of ΔF/F_0_, with F_0_ representing basal fluorescence measured before LPI application.

**Cell Culturing**

Hippocampal neurons were cultured from postnatal day 0 male and female Sprague-Dawley rat pups. The hippocampus was isolated in ice-cold HBSS (Corning) containing 20% fetal bovine serum (FBS), and washed in HANKS without serum. Following washing, hippocampi were digested for 8 min. in a 1 ml papain solution (Papain dissociation System, Worthington). 50 units of DNase I (Millipore Sigma) and 0.5 μM MgCl_2_) was added at the end of Papain digestion. Digestion was stopped by adding 5 ml of modified HBSS containing 20% fetal bovine serum. After additional washing, the tissue was dissociated using Pasteur pipettes of decreasing diameter. The cell suspension was pelleted and plated on 10 mm coverslips coated with poly-D-lysine in 24-well plates for immunocytochemistry, or directly onto PDL-coated 12 well plates for western blot studies. The cultures were maintained in NbActiv4 (BrainBits, Springfield, IL). A 50% medium change was performed at 7 days, and once per week thereafter. Neurons were used for experiments 12-14 days in vitro (DIV) after plating.

A subset of cultures were transfected at 3DIV (MOI=5) with 4 custom designed lentiviral, GFP+ shRNA constructs (Origene) targeting GPR55 with the following sequences, along with scrambled controls:

5’-TGAGTCAGCTAGACAGTAACAACTGCTCG-3’

5’-CAACCTGGCTGTCTTCGACTTACTGCTTG-3’

5’-CTGGACCATTGCTACCAATCTTGTCGTCT-3’

5’-CTCAATGTAGTTCAGCCATAGCAGAATGA-3’

**Immunostaining**

For all cell culture experiments involving LPI treatment, cells were treated for 1 hour with pre-warmed 4mM K Tyrode’s solution consisting of (in mM): 150 NaCl, 4 KCl, 2 MgCl_2,_ 2 CaCl_2_, 10 HEPES, 10 glucose, pH 7.4, and synaptic blockers 1μM TTX, 10μM NBQX, and 50 μM APV. 4μM LPI, or DMSO vehicle, was then applied in Tyrode’s solution with synaptic blockers. Cultured cells were then fixed at 30 or 60 minutes post drug application in ice-cold 4% paraformaldehyde in PBS supplemented with 20 mM EGTA and 4% (w/v) sucrose. Fixed cells were then permeabilized with 0.1% Triton X-100, blocked with 5% normal donkey serum, and incubated overnight with primary antibodies (see table below). The next day, cells were washed with PBS, incubated at room temperature for 50 min with Alexa secondary antibodies (1:1000, Molecular Probes), washed with PBS (3x5 min) and mounted using with ProLong Gold Antifade Mountant with or without DAPI (ThermoFisher Scientific).

For all experiments involving *ex vivo* hippocampal brain slices, animals were perfused with PBS followed by 4% paraformaldehyde in PBS. Whole brains were dissected and fixed overnight in 4% PFA, and then sucrose-protected overnight in 30% sucrose in PBS before embedding and freezing in optimal cutting temperature compound (Tissue-Tek O.C.T.) for sectioning. Frozen sections were cut on a cryostat at 16 μm and collected on HistoBond coated slides (VWR) for staining. Sections were blocked for 2-3 hours at room temperature in 0.2% Triton X-100 and 5% normal serum, then incubated at 4°C in primary antibodies (see chart below).

Sections were rinsed and then incubated in species-appropriate Alexa Fluor conjugated secondary antibodies (Molecular Probes, 1:500) for 2-3 hours, and finally mounted with ProLong Gold Antifade Mountant with or without DAPI (ThermoFisher Scientific). Fluorescent images were acquired on a Zeiss LSM 510 meta Imager.M1 confocal microscope at 10x, 20x, or 63x magnification.

Images were analyzed for both average puncta intensity and colocalization using custom analysis scripts in Icy (<http://icy.bioimageanalysis.org>) software (de Chaumont et al., 2012). Regions of interest (ROI) were traced around apical dendrites labeled by MAP2 or axons labeled by tau. For puncta colocalization analyses, immunoreactive puncta for GPR55 and synaptic markers (e.g. VGLUT1, VGAT) were separately detected with sub-pixel accuracy, in 3D, and in an automated manner using the ‘Spot Detector’ plugin of ICY (scale 2, threshold = 100). Puncta were determined to be “colocalized” with a threshold of 250 nm in the horizontal (XY) plane and 500 um in the depth (Z) plane, based on average pixel sizes of 120 nm XY and 370 nm Z, respectively. The 3D colocalization ratio was then computed as: (#colocalized GPR55 puncta + #colocalized marker puncta)/(total # GPR55 puncta + total #marker puncta). In order to gain a statistical insight on the estimated correspondence between puncta, we adopted a Monte Carlo simulation approach in which we aimed at building the probabilistic distribution of the colocalization ratio under the null hypothesis that the protein GPR55 is uniformly distributed along the axon and that colocalization with synaptic markers occurs only by chance. We randomly redistributed the location of GPR55 immunoreactive puncta along the axon in 2D after detection, while keeping their number fixed. Briefly, using the tau (axonal) or MAP2 (dendritic) fluorescence ROIs in 2D, we traced a series of linked 2D segments and orthogonally projected the detections for the two proteins along them so that a detection with 3D coordinates (x, y, z) is then represented by a single value (t) corresponding to the 1D distance to the tip of the traced axon/dendrite. We then computed the colocalization ratio as before in this coordinate system (colocalization threshold = 250 nm). For simulated data, we performed 10^4^ trials in which the location of synaptic marker puncta remained fixed to preserve the spatial distribution of the protein, while GPR55 detections were randomly uniformly redistributed along the axonal segment. For each axonal segment, we compared the averaged simulated colocalization ratio with the measured one and we statistically pooled data from multiple coverslips using a paired t-test.

**Immunoblotting**

Cells were harvested and lysed (Lysis Buffer A, Thermo) with EDTA-free protease and phosphatase inhibitors (Thermo) following treatment with 4μM LPI (see immunostaining above for description). In a subset of studies, samples were treated with pharmacological blockers (see table below), and lysed at various times post LPI application. Sample proteins were equalized using the BCA Protein Assay Kit (Pierce) and loaded into 10-20% SDS-PAGE gel with 5% beta-mercaptoethanol following heating at 95^o^C for 5 min. Cellular protein was transferred to Immobilon transfer membrane (Millipore). The membrane was then blocked at room temperature for 2 hrs in Odyssey blocking buffer (Li-Cor Biosciences, Lincoln, NE), and incubated with primary antibodies (see table below). The following day, the membrane was washed with 0.1% Tween 20 in PBS and incubated with an IRDye-labeled secondary antibody (Li-Cor) for 1 hr. After incubation, the membrane was washed with PBS and imaged with Odyssey imaging systems. All the bands were analyzed with Image Studio software.

**Total RNA extraction and qPCR**

For the PTZ (105mg/kg, i.p.)-induced seizure model, one hippocampus was dissected out on ice and used for total RNA extraction at the time point either immediately after seizure-induced death, or 30 minutes after PTZ/vehicle treatment, whichever came first. For KA-induced seizure model, one hippocampus was dissected out on ice 48 hours following either KA (24 mg/kg, i.p., “KA1”) or vehicle. For both PTZ and KA experiments, CBD (200 mg/kg) was administered 1 h prior to chemoconvulsant induction in a subset of animals. RNeasy Mini Kit (Qiagen, Cat#74104) was used to extract total RNA as per manufacturer instructions. Total RNA was eluted in 30 μl of RNase-free water and concentration was measured using nanodrop. SuperScript III Reverse Transcriptase (ThermoFisher Scientific #18080044) was used for reverse transcription of 1 μg (for the PTZ experiments) or 500 ng (for the KA experiments) total RNA from each sample following the user manual. qPCR was performed with SsoAdvanced Universal SYBR Green Supermix (Bio-Rad# 1725272) using 1 μl cDNA as template. Mouse GPR55 qPCR primers were designed to span an intron and were confirmed with non-RT control to make sure there is no genomic DNA interference (Suppl. Fig. 7A-B), and all PCR products were confirmed by Sanger sequencing. Two internal control genes (*Hmbs* and *Sdha*) were used with geometric averaging to ensure accurate normalization (Vandesompele et al., 2002).

For the qPCR assay of Li-PLC-induced seizure model in rats, hippocampi were dissected 3-6 months following establishment of epileptogenesis with age-matched non-epileptic controls, and flash frozen in liquid nitrogen. One half of a hippocampus (~30 mg) from each rat was used to extract total RNA using the RNeasy Mini Kit. To eliminate genomic DNA contamination, column-purified RNA were treated with RNase-free DNase I (Ambion, AM2224) at 37 ^o^C for 30 minutes followed by 75 ^o^C inactivation for 10 minutes and column cleaning. Non-RT controls, in which no reverse transcriptase was added in the reverse transcription reaction, were run for all samples to monitor the level of genomic DNA contamination in the case of all-isoform *Gpr55* qPCR. We tested different promoter usages of rat *Gpr55* and used primer 10&11 (below) in Fig. 7B.

Primers used for qPCR:

| ID | Primer | Sequence | Notes |
| --- | --- | --- | --- |
| 1 | mHmbs-qPCRF | gagaaagttcccccacctgg | 1&2: mouse internal control 1 |
| 2 | mHmbs-qPCRR | ccaggacgatggcactgaat |  |
| 3 | mSdha-qPCRF | tgcggctttcacttctctgt | 3&4: mouse internal control 2 |
| 4 | mSdha-qPCRR | cgcctacaaccacagcatca |  |
| 5 | mGpr55-qPCRF | gcttggggacagaagtgtga | 5&6: mouse *Gpr55*, flanking intron; isoform-specific; **(Fig. 6C, Suppl Fig. 7A-B)** |
| 6 | mGpr55-qPCRR | gctgcaaggttctggtaagc |  |
| 7 | ratHmbs-qPCRF | ggacctggttgttcactccc | 7&8: rat internal control 1 |
| 8 | ratHmbs-qPCRR | ggcaaggtttccagggtctt |  |
| 9 | ratSdha-qPCRF | cgctcacatactgttgcagc | 9&10: rat internal control 2 |
| 10 | ratSdha-qPCRR | tcagagcctttcacggtgtc |  |
| 11 | ratGpr55-qPCRF2 | tcagcccgagaaggaactgcttc | 11&12: rat Gpr55, flanking intron; isoform-specific; corresponding to 5&6 in mouse **(used in Fig. 7B)** |
| 12 | ratGpr55-qPCRR2 | tggtcaggttgtccacgaaaacgaa |  |

**HPLC/MS**

*Extraction of LPIs from brain tissue* – Prior to extraction, samples were moved from

-80 °C storage to dry ice and weighed into 2.0 mL screw cap vials containing ~100 µL of disruption beads (Research Products International, Mount Prospect, IL). Each sample was normalized to 46 mg/mL using 100% methanol (Fisher Scientific) and homogenized for 10 cycles on a bead blaster homogenizer (Benchmark Scientific, Edison, NJ). Cycling consisted of a 30 sec homogenization time at 6 m/s followed by a 30 sec pause. In glass LC vials, 80 µL of Optima LC/MS grade water (Fisher Scientific, Waltham, MA) was combined with 80 µL of homogenate. Further extraction was performed by adding 160 µL of 100% chloroform and vortexing for 2 min. Entirety of sample was transferred to glass inserts and spun at 21,000 g for 3 min at 4 °C. For analysis, 20 µL of each sample was transferred to LC/MS vials containing glass inserts.

*LC-MS/MS targeted LPI method* – Standards of 16:0, 18:0 and 20:4 LPI (Avanti Polar Lipids) were prepared in ethanol at 1uM, 3uM, 10uM, 30uM, and 100uM and used to validate retention time and fragmentation pattern for each LPI. Samples were analyzed by UPLC-MS/MS with a targeted product reaction monitoring (PRM) method for 16:0, 18:0 and 20:4 LPI. The LC column was a Waters^TM^ BEH-C18 (1.0 x 50 mm, 1.7 μm) coupled to a Dionex Ultimate 3000^TM^ system and the column oven temperature was set to 25^o^C for the gradient elution. The flow rate of 0.1 mL/min was used with the following buffers; A) 60:40 acetonitrile:water, 10 mM ammonium formate, 0.1% formic acid and B) 90:10 isopropanol:acetonitrile, 10 mM ammonium formate, 0.1% formic acid. The gradient profile was as follows; 50-100%B (0-2.0 min), hold at 100%B (0.5 min), 100-50%B (2.5-3 min), hold at 50%B (3.0 min). Injection volume was set to 1 μL for all analyses (6 min total run time per injection). MS analyses were carried out by coupling the LC system to a Thermo Q Exactive HF^TM^ mass spectrometer operating in heated electrospray ionization mode (HESI). Method duration was 6 min with a PRM scan in negative mode only. Global list for PRM contained [M-H]^-^ theoretical masses for targeted LPIs. Spray voltage for was 3.5kV and capillary temperature was set to 320^o^C with a sheath gas rate of 25, aux gas of 10, and max spray current of 100 μA. Tandem MS scans utilized 30,000 resolution with a maximum IT of 100 ms, isolation window of 0.4 m/z, isolation offset of 0.1 m/z, and normalized collision energies (nCE) of 35. The minimum AGC target was 2e5 with an intensity threshold of 1e6. All data were acquired in profile mode. Duplicate standard curve points and samples were randomized for sequence injection. Standard curves were generated for each LPI. For quantitation, a background of 3x the average blank signal plus 10,000 counts was applied. Sample quantities were calculated using linear regressions to determine on column amount of each LPI. Assuming 100% extraction efficiency, calculated values were used to determine amount of LPI per mg of tissue.

**Statistics and Analysis**

Data analyses were performed using Clampfit (Molecular Devices, UK), Icy (http://icy.bioimageanalysis.org), and Prism 6 (GraphPad Software, USA) software. Data are presented as means ± SEM. For PTZ-induced seizure and mortality incidence, Chi-squared tests or Fisher's exact tests were used. For electrophysiological and molecular biological data, distributions passing Shapiro-Wilk test for normality were compared using a two-way student’s t-test. Non-Gaussian distributions were compared using the non-parametric tests Wilcoxon signed rank test. For recordings pre- and post-drug application within a given cell, paired t-tests were used for comparisons. Events were normalized to the mean PSC frequency or amplitude during a 5 min window of baseline. For *in vitro* LPI experiments, intensities were normalized to the average of the vehicle-treated condition. For 3 or more independent comparisons (including electrophysiological recordings, HPLC, qPCR, immunocytochemistry, and immunoblots), a one-way ANOVA followed by post-hoc testing (e.g. Dunnett’s or Tukey’s) was performed. Differences were considered significant at p < 0.05.

Antibodies:

| **Antibody** | **Company** | **Catalog #** | **Concentration** |
| --- | --- | --- | --- |
| Rb α GPR55* | Cayman Chemicals | 10224 | 1:500 |
| MS α VGLUT1 | Synaptic Systems | 135 311 | 1:100 |
| MS α VGAT | Synaptic Systems | 131 011 | 1:100 |
| GT α PSD-95 | Abcam | Ab12093 | 1:1000 |
| CK α Gephyrin | Abcam | Ab136343 | 1:200 |
| MS α CB_1_R | Synaptic Systems | 258 011 | 1:500 |
| GP α Tau | Synaptic Systems | 314 004 | 1:1000 |
| GP α MAP2 | Synaptic Systems | 188 004 | 1:1000 |
| RB α GABA_A_R Gamma_2_ | Synaptic Systems | 224 003 | 1:500  (WB, ICC) |
| RB α GABA_A_R Gamma_2_ phospho-S327 | Phospho-solutions | P1130-327 | 1:500 (WB) |
| RB α GABA_A_R Beta_3_ | Synaptic Systems | 224 404 | 1:500 (WB) |
| RB α GABA_A_R Beta_3_ phospho-S408/409 | Phospho-solutions | P1130-4089 | 1:500 (WB) |
| RB α Beta Actin | Cell Signaling | 4970 | 1:10,000 (WB) |
| MS α GAPDH | GeneTex | GTX627408 | 1:10,000 (WB) |

*Cayman chemicals GPR55 antibody targets an internal cytoplasmic region of GPR55 protein, amino acids 7 to 20

Pharmacological blockers

| **Compound** | **Target** | **Company/**  **Catalog#** | **Working concentration** |
| --- | --- | --- | --- |
| Y-27632 | Rho-associated protein kinase (ROCK) | Cayman Chemicals  #10005583 | 10 μM |
| U 73122 | PLC | Tocris  #1268 | 10 μM |
| Bisindolyl-  maleimide II | PKC | Tocris #1138 | 0.1 μM |
| Thapsigargin | ER Ca^2+^-ATPase | Tocris #1138 | 10 μM |
| (-)-Xesto-  spongin C | IP3R | Cayman Chemicals  #64950 | 1 μM |
| Cyclosporin A  (CsA) | Calcineurin | Tocris  #1101 | 10 μM |
| Tautomyectin | PP1 α | Tocris  #2305 | 5 nM |
| CID16020046 | GPR55 | Cayman  #15247 | 2.5 μM |
| AACOCF3 | PLA2 | Cayman  # 62120 | 20 μM |
| YM 26734 | PLA2 | Cayman #17631 | 10 μM |

***Supplemental References***

de Chaumont, F., Dallongeville, S., Chenouard, N., Herve, N., Pop, S., Provoost, T., Meas-Yedid, V., Pankajakshan, P., Lecomte, T., Le Montagner, Y.*, et al.* (2012). Icy: an open bioimage informatics platform for extended reproducible research. Nat Methods *9*, 690-696.

Gelinas, J.N., Khodagholy, D., Thesen, T., Devinsky, O., and Buzsaki, G. (2016). Interictal epileptiform discharges induce hippocampal-cortical coupling in temporal lobe epilepsy. Nature medicine *22*, 641-648.

Modebadze, T., Morgan, N.H., Peres, I.A., Hadid, R.D., Amada, N., Hill, C., Williams, C., Stanford, I.M., Morris, C.M., Jones, R.S.*, et al.* (2016). A Low Mortality, High Morbidity Reduced Intensity Status Epilepticus (RISE) Model of Epilepsy and Epileptogenesis in the Rat. PloS one *11*, e0147265.

Patra, P.H., Barker-Haliski, M., White, H.S., Whalley, B.J., Glyn, S., Sandhu, H., Jones, N., Bazelot, M., Williams, C.M., and McNeish, A.J. (2019). Cannabidiol reduces seizures and associated behavioral comorbidities in a range of animal seizure and epilepsy models. Epilepsia *60*, 303-314.

Pohl, M., and Mares, P. (1987). Effects of flunarizine on Metrazol-induced seizures in developing rats. Epilepsy Res *1*, 302-305.

Racine, R.J. (1972). Modification of seizure activity by electrical stimulation. I. After-discharge threshold. Electroencephalogr Clin Neurophysiol *32*, 269-279.

Van Erum, J., Van Dam, D., and De Deyn, P.P. (2019). PTZ-induced seizures in mice require a revised Racine scale. Epilepsy & behavior *95*, 51-55.

Vandesompele, J., De Preter, K., Pattyn, F., Poppe, B., Van Roy, N., De Paepe, A., and Speleman, F. (2002). Accurate normalization of real-time quantitative RT-PCR data by geometric averaging of multiple internal control genes. Genome Biol *3*, 7 RESEARCH0034.

Vilela, L.R., Lima, I.V., Kunsch, E.B., Pinto, H.P.P., de Miranda, A.S., Vieira, E.L.M., de Oliveira, A.C.P., Moraes, M.F.D., Teixeira, A.L., and Moreira, F.A. (2017). Anticonvulsant effect of cannabidiol in the pentylenetetrazole model: Pharmacological mechanisms, electroencephalographic profile, and brain cytokine levels. Epilepsy & behavior *75*, 29-35.
